# Phenotypic divergence and genomic architecture between parallel ecotypes at two different points on the speciation continuum in a marine snail

**DOI:** 10.1101/2023.09.21.558824

**Authors:** Francesca Raffini, Aurélien De Jode, Kerstin Johannesson, Rui Faria, Zuzanna B. Zagrodzka, Anja M. Westram, Juan Galindo, Emilio Rolán-Alvarez, Roger K. Butlin

**Affiliations:** Ecology and Evolutionary Biology, School of Biosciences, The University of Sheffield, Sheffield S10 2TN, UK; Department of Life Sciences and Biotechnology, University of Ferrara, 44121, Ferrara, Italy; Department of Biology and Evolution of Marine Organisms, Stazione Zoologica Anton Dohrn, Villa Comunale, 80121, Naples, Italy; Department of Marine Sciences, University of Gothenburg, Tjärnö Marine Laboratory, 45296 Strömstad, Sweden; Department of Biological Sciences, University of Alabama, Tuscaloosa, AL, 35487, USA; Dauphin Island Sea Lab, Dauphin Island, AL, 36525, USA; CIBIO, Centro de Investigação em Biodiversidade e Recursos Genéticos, InBIO Laboratório Associado, Campus de Vairão, Universidade do Porto, 4485-661 Vairão, Portugal; BIOPOLIS Program in Genomics, Biodiversity and Land Planning, CIBIO, Campus de Vairão, 4485-661 Vairão, Portugal; Faculty of Biosciences and Aquaculture, Nord University, 8049 Bodø, Norway; Institute of Science and Technology Austria, 3400 Klosterneuburg, Austria; Centro de Investigación Mariña, Universidad de Vigo, Departamento de Bioquímica, Genética and Immunología, 36310 Vigo, Spain

## Abstract

Speciation typically occurs over too long a time frame to be observed directly. A way forward is to compare pairs of ecotypes that evolved in parallel in similar contexts but have reached different degrees of reproductive isolation. Such comparisons are possible in the marine snail *Littorina saxatilis* by contrasting barriers to gene flow between parallel ecotypes in Spain and Sweden. In both countries, divergent ecotypes have evolved to withstand either crab predation or wave action. Here, we explore transects spanning contact zones between the Crab and the Wave ecotypes using low-coverage whole-genome sequencing, morphological and behavioural traits. Despite parallel phenotypic divergence, distinct patterns of differentiation between the ecotypes emerged: a continuous cline in Sweden indicating a weak barrier to gene flow, but two highly genetically and phenotypically divergent, and partly spatially-overlapping clusters in Spain suggesting a much stronger barrier to gene flow. Absence of Spanish early-generation hybrids supported strong isolation, but a low level of gene flow is evident from molecular data. In both countries, highly differentiated loci were located in both shared and private chromosomal inversions but were also present in collinear regions. Despite being considered the same species and showing similar levels of phenotypic divergence, the Spanish ecotypes are much closer to full reproductive isolation than the Swedish ones. Barriers to gene flow of very different strengths between ecotypes within the same species might be explained by dissimilarities in the spatial arrangement of habitats, the selection gradients or the ages of the systems.

Graphical abstract.
Distinct patterns of divergence between parallel ecotypes in Sweden and Spain.

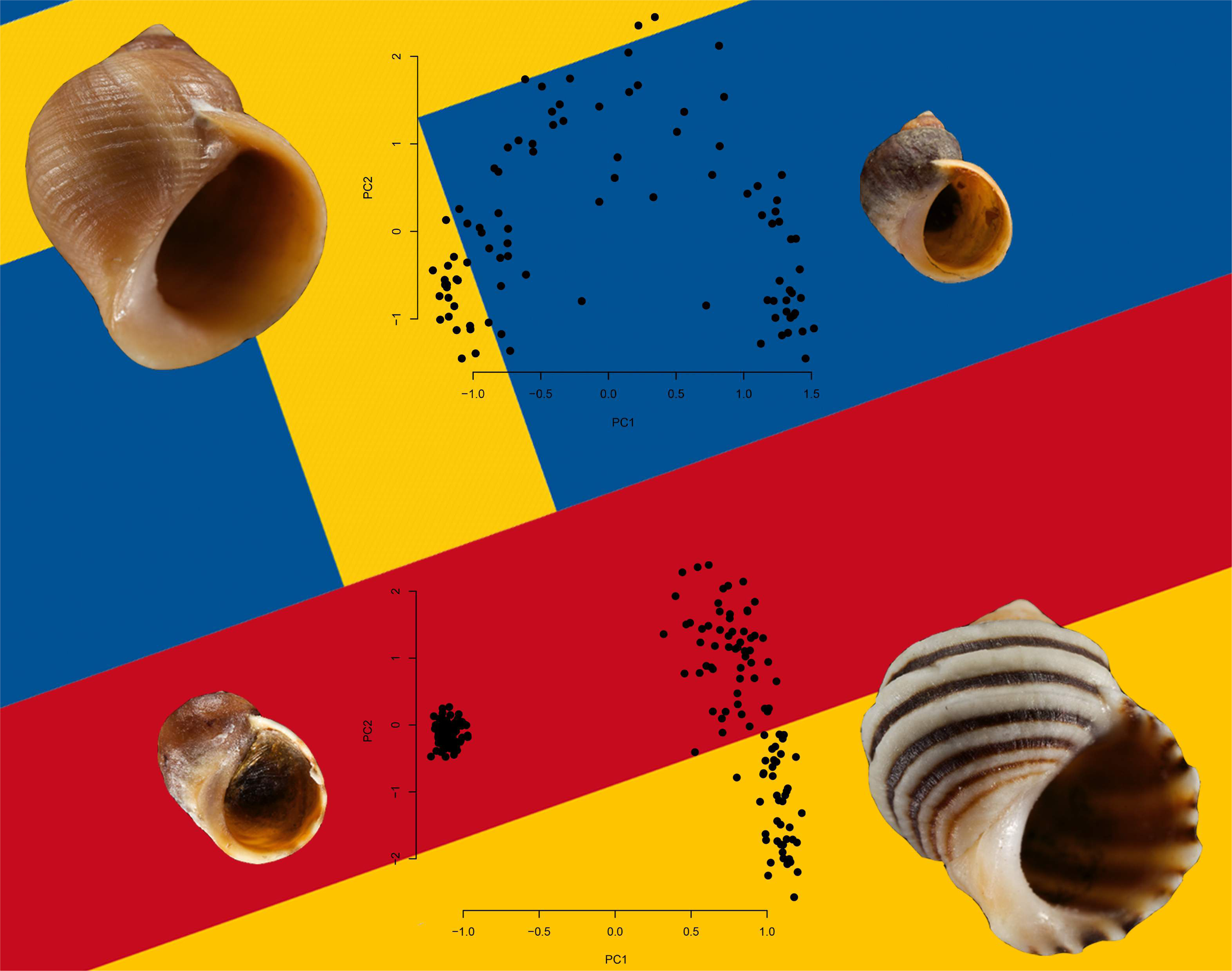

## Introduction

The process of speciation involves the build-up of reproductive isolation. The precise meaning of ‘reproductive isolation’ has recently been debated (Westram *et al*., 2022 and associated commentaries); here, we use the term in a general sense to include both reduction in interbreeding and reduction in gene exchange. The time-scale over which speciation occurs is highly variable (Coyne & Orr, 2004) but it is typically too long for direct observation. Therefore, inferences about the accumulation of reproductive isolation often depend on comparing contemporary pairs of populations at different points on a “speciation continuum” (Seehausen & Wagner, 2014; Stankowski & Ravinet, 2021). Due to the complex set of processes involved in speciation, it can be helpful to think about this continuum in multiple dimensions (Bolnick *et al*., 2023; K. Johannesson *et al*., 2024). This approach can provide many insights, for example in considering whether there are early- and late-evolving components of reproductive isolation, or how patterns of gene exchange across the genome are modified at different stages of divergence (Feder *et al*., 2012). However, a monotonic progression from weak to strong isolation cannot be assumed (Bolnick *et al*., 2023; Stankowski & Ravinet, 2021). More studies of population pairs across the speciation continuum are needed, particularly within clades so that comparisons among different levels of reproductive isolation are not confounded with differences among taxa that may not be relevant to the speciation process (Seehausen & Wagner, 2014). Significant progress has been made in this direction, especially in terms of genome-wide patterns of genetic differentiation (e.g., Butlin & Faria, 2024; Fang *et al*., 2020, 2021; Jiggins, 2017; K. Reid *et al*., 2021; Riesch *et al*., 2017). The number of investigations in different study systems is, however, still limited and many questions about the mechanisms leading to completion of speciation remain open, particularly in the case of speciation with gene flow.

The appearance of the first components of reproductive isolation is often relatively well understood, but explaining the later accumulation of additional components of isolation and the final cessation of gene flow remains challenging (Butlin *et al*., 2008; Butlin & Smadja, 2018; Kulmuni *et al*., 2020). Many factors have been suggested that might lead to stronger reproductive isolation. Time is needed for the accumulation of incompatibilities (e.g., Guerrero *et al*., 2017), and possibly also for response to divergent selection if this is mutation limited (Barrett & Schluter, 2008), and so older population pairs might be more strongly isolated, as is often observed (Coyne & Orr, 1989; Matute & Cooper, 2021). Stronger divergent selection, for example between more distinct habitats, should also lead to stronger isolation (Funk *et al*., 2006). Some spatial arrangements of populations might be more conducive to the evolution of reproductive isolation than others, because gene flow opposes divergence (Coyne & Orr, 2004) but contact also provides the opportunity for reinforcement (Servedio & Noor, 2003; Yukilevich, 2021). Furthermore, cycles of population expansion and contraction can act to bring components of reproductive isolation together (Butlin & Smadja, 2018; Hewitt, 1989). Opportunities for the evolution of assortative mating, either through mating behaviour or due to habitat association, can lead to strong reproductive isolation particularly when they are aligned to divergent selection (Kopp *et al*., 2018). Finally, one-allele barrier effects (Butlin *et al*., 2021; Felsenstein, 1981), pleiotropy and multiple-effect traits (Servedio *et al*., 2011; Smadja & Butlin, 2011) might promote the evolution of reproductive isolation. Likewise, genomic architecture, i.e., numbers and effect sizes of barrier loci, their genomic distribution and patterns of recombination, including the effects of structural variants, may be important because the coupling of individual barrier loci and of barrier effects can be opposed by recombination (Butlin & Smadja, 2018; Dopman *et al*., 2024; Felsenstein, 1981).

The intertidal gastropod genus, *Littorina*, is a model system in which several of these issues can be addressed (K. Johannesson *et al*., 2010a; K. Johannesson, 2016; K. Johannesson *et al*., 2017, 2024; Rolán-Alvarez *et al*., 2015). *Littorina saxatilis* is widespread and abundant on North Atlantic shores. Its ovoviviparous reproduction and resulting short lifetime dispersal have allowed it to adapt to many different habitat types (D. G. Reid, 1996). Ecotypes adapted to habitats with a high risk of crab predation but less direct wave action (the ‘Crab’ ecotype), or habitats with strong wave action but low predation risk (the ‘Wave’ ecotype), occur in close proximity on many shores and form contact zones where habitats meet. These ecotypes differ in a suite of adaptive traits including size, shell shape and thickness, and behaviour (B. Johannesson & Johannesson, 1996). Demographic models suggest that they evolved in parallel in multiple countries (Butlin *et al*., 2014; Carvalho, Faria, *et al*., 2023) although this is likely to have involved repeated use of at least some adaptive genetic variants (Morales *et al*., 2019), a proportion of which are found in chromosomal inversions (Faria, Chaube, *et al*., 2019; Koch *et al*., 2021, 2022; Reeve *et al*., 2023; Westram *et al*., 2021). Background levels of genetic divergence between the Crab and Wave ecotypes and patterns of genomic differentiation determined using pooled sequencing data suggested that the strength of the barrier to gene flow between ecotypes varies widely among locations in Europe (Morales *et al*., 2019).

In this study, we compared and contrasted the patterns of divergence between Crab and Wave ecotypes in two geographic regions: the Swedish west coast, colonised since the last glaciation (divergence estimated ∼15 kya), and the Galician coast in Spain (divergence estimated ∼57 kya), where older populations survived the Pleistocene glaciations *in situ* (Bosso *et al*., 2022; Butlin *et al*., 2014; Carvalho, Faria, *et al*., 2023; Carvalho, Morales, *et al*., 2023; Doellman *et al*., 2011; Panova *et al*., 2011). In Sweden, the tidal range is small, the Crab-Wave axis is parallel to the shore-line (horizontal zonation), and the two ecotypes hybridize in narrow contact zones where sheltered boulder fields abut rocky headlands. There is a genome-wide barrier to gene exchange, but it is weak: background F_ST_ is about 0.04 (K. Johannesson *et al*., 2024), clinal changes in SNP frequency are wide-spread in the genome but fixed differences are rare, loci putatively contributing to barrier effects occur both within and outside polymorphic inversions (Koch *et al*., 2022; Westram *et al*., 2018, 2021). The barrier in Sweden appears to be due primarily to local adaptation, without noticeable intrinsic incompatibilities, but with some contribution of size-assortative mating (Hollander *et al*., 2005; K. Johannesson *et al*., 2008; Perini *et al*., 2020).

In Spain, the tidal range is much greater and the two ecotypes are distributed on a perpendicular up-down shore axis (vertical zonation) with the Crab ecotype primarily in the barnacle belt in the high shore, where predation is most intense, and the Wave ecotype among blue mussels in the low shore, where wave action is strongest. The Spanish ecotypes overlap in a contact zone in the mid shore, characterized by a mosaic distribution of barnacle and mussel patches (K. Johannesson *et al*., 1993, 1995; Rolán-Alvarez *et al*., 1997, 1999). There are some indications that the barrier to gene flow between ecotypes is stronger in Spain than in Sweden: background F_ST_ is higher, around 0.1 (Butlin *et al*., 2014; Morales *et al*., 2019; Westram *et al*., 2021), and only a few intermediate genotypes were observed in putatively hybrid samples using reduced representation (RADseq) data (Kess *et al*., 2018). Yet, an investigation of the pattern of differentiation in Spain with a combination of genome-wide data and detailed spatial coverage of the contact zone has been lacking.

Here, we take a step further into understanding the drivers of speciation, particularly at the molecular level. We describe the phenotypic and genomic patterns of differentiation between Crab and Wave snails in Spain using low-coverage whole-genome sequence data, shell features and behavioural traits for snails from dense transects across the contact zone. We make a direct comparison with a transect in Sweden as well as with published analyses based on pooled or capture sequencing. Despite studying populations of the same species and similar phenotypic divergence (Crab and Wave ecotypes), we report very different patterns of genomic differentiation in the contact zones of the two countries: in Spain, ecotypes partly overlap in space with evidence for only limited, almost unidirectional introgression, while in Sweden there is a gradual phenotypic as well as genetic transition from one ecotype to the other. Thus, despite being populations of the same species (*L. saxatilis*), pairs of ecotypes in different environmental and demographic contexts have very different positions on the speciation continuum, and we discuss potential reasons for this.

## Materials and Methods

### Sampling

Snails were sampled to include the typical habitats of both the Crab and Wave ecotypes and the contact zone in between, in both Spain and Sweden. In Sweden, snails were sampled in a single transect along the shore from a boulder field (‘Crab’ habitat) to a rocky headland (‘Wave’ habitat) on the island of Ramsö (58°49′27.8″N, 11°03′45.3″E; a re-sampling of transect CZA_right from Westram *et al*., 2021). Note that the tidal amplitude is only 35 cm in Sweden, but high and low water level also vary with air pressure and wind direction up to a maximum amplitude of 1.5 m. In all parts of the transect, snails were collected from positions scattered throughout their vertical distribution (∼1m). Seven hundred snails were sampled in June 2015 along the transect, without reference to phenotype but aiming to avoid juveniles. The position of each snail was recorded in three dimensions using a total station (Trimble M3). For spatial analysis, we placed each snail on a ‘least cost path’ (as described in Westram *et al*., 2021) and distances were transformed to start from zero at the Crab ecotype end of the transect.

In Spain, snails were collected from Centinela on the west coast of Galicia (N 42° 4’ 38.06”, W 8° 53’ 47.47”) in spring (March) and autumn (September) of 2017. Each sample consisted of approximately 600 snails collected in the same way as in Sweden from two transects perpendicular to the shore, and about 5m wide and separated by 2-10 m, stretching from the upper limit of the *L. saxatilis* distribution in the splash zone to its lower limit close to low water of spring tides (tidal range ∼4m). Sampling was more dense in the lower part of the shore where hybridization between the two previously described ecotypes was expected (Galindo *et al*., 2013). For spatial analysis, we used the position of each snail on a single shore-position axis running through the middle of each transect. Distances were transformed such that each transect started from the top of the distribution (defined as zero) and ended at roughly low water level as indicated by the lowest collected snails. This corresponds to a vertical range of approximately 3 m and shore height was recorded relative to the position of the lowest individual sampled. To include habitat information, the presence or absence of barnacles (*Chthamalus* spp., typical of the mid to upper shore), mussels (*Mytilus galloprovincialis,* from mid to lower shore), and goose barnacles (*Pollicipes pollicipes*, lower shore) was recorded within 5 cm of each snail position and for approximately uniformly distributed positions across the area containing the two sampling transects. Snails were stored in individual tubes, moistened with seawater and kept at 4°C until phenotyping was complete. Then, the head and foot of each snail was dissected and preserved in 99% ethanol.

### Sequencing and read processing

Seventy-three adult snails from Spain in spring, 114 from Spain in autumn and 96 from Sweden were selected randomly relative to phenotype to cover the full transect range within each country. DNA was extracted from foot tissue using a modified CTAB protocol (Panova *et al*., 2016). In-house, high-throughput genomic DNA library preparation and whole genome sequencing (Illumina HiSeq4000, 150bp, eight lanes, paired-end) were performed by The Oxford Genomics Centre with a target coverage of 3x based on the estimated genome size of 1.35Gb (Westram *et al*., 2018).

Raw reads were trimmed with Trimmomatic v. 0.38 (Bolger *et al*., 2014), retaining reads with a minimum length of 70bp after filtering, and mapped to the *L. saxatilis* draft reference genome (Westram *et al*., 2018) using bwa mem v. 0.7.17 (Li, 2013) and default settings. Positions with base or map quality lower than 20 were discarded using Samtools v. 1.7 (Li, 2011; Li *et al*., 2009). PCR duplicates and overlap between paired-end reads were removed with Picard v. 2.0.1 (http://broadinstitute.github.io/picard/) and bamUtil (Jun *et al*., 2015). As specimens were sequenced in paired-ends in eight lanes, resulting in 16 outputs for each snail, the files belonging to the same individual were sorted and merged with Samtools.

To explore patterns of diversity within Spain and Sweden, variants were called separately in each country using samtools mpileup and bcftools call v. 1.11 (Li, 2011) including only the longest contigs covering 90% of the reference genome. Allelic read counts for each SNP rather than genotype calls were retained due to the low coverage. The resulting vcf files were filtered to retain only biallelic SNPs, positions where at least 50% individuals had a read depth between one and 18 irrespective of the reference or alternative allele, and a minor allele frequency higher than 0.05 using vcftools v. 0.1.14 (Danecek *et al*., 2011) and vcffilterjs (Lindenbaum & Redon, 2018). Additionally, we retained only positions on the *L. saxatilis* linkage map (Westram *et al*., 2018), i.e. within 1,000 bp of a SNP that could be positioned on this map.

Two types of datasets were generated for each country: unthinned and thinned. The first one included all the SNPs resulting from the processing described above without any further filtering. The second one was obtained by retaining one random SNP in each genomic window of 1,000 bp to reduce the impact of linkage disequilibrium and avoid overweighting regions with high SNP density. To achieve this aim, the reference genome was sliced into bins of 1,000 bp and one SNP was randomly picked in each window using the R v. 4.0.3 (R Core Team, 2020) package GenomicAlignments v. 1.26.0 (Lawrence *et al*., 2013), vcftools and custom scripts. This random SNP subsampling was repeated three times in each country to create a total of six random SNP subsets from the unthinned datasets.

To investigate overall divergence between countries, variants were called and filtered jointly in Sweden and Spain and a thinned dataset was generated using the procedures described above.

In all datasets (within and between countries, thinned and unthinned), the reference and alternative allele read depth was extracted from the vcf files and one random read per position and individual was subsampled using vcftools and custom scripts. As for the random SNP subsets, the random read subsampling was repeated three times to create replicates for each SNP dataset and subset. These ‘single read’ data sets formed the basis of population genomic analysis, thereby avoiding the issues associated with calling genotypes from low-coverage data (Nielsen *et al*., 2011).

### Genomic analyses

Overall patterns of divergence within countries were explored in the thinned datasets through PCA and DAPC in the R packages adegenet (Jombart, 2008; Jombart & Ahmed, 2011) and factoextra v. 1.0.7 (https://github.com/kassambara/factoextra) treating individuals as haploid. If more than one genetic group was identified, the within-group PC1 scores among subsets were tested for normality using the Shapiro test (Shapiro & Wilk, 1965). We compared group (if any) assignment of each individual between subsampled data sets and computed the correlation of the within-group PC1 scores using the Pearson or Spearman’s coefficients, according to the data distribution, using the R package GGally v. 2.1.2 (https://ggobi.github.io/ggally). Loci with an allele frequency difference between groups (if any) > 0.5 were used to compute a Hybrid Index for each individual (mean over genotypes expressed as 1 for the allele more common in the ‘Crab’ environment and 0 for the allele more common in the ‘Wave’ environment). Individual ancestries were estimated through a maximum likelihood approach and cross validation procedures using ADMIXTURE v. 1.3.0 (Alexander *et al*., 2009), PLINK v. 1.90b6.5 (Purcell *et al*., 2007) and custom scripts. We tested associations between genetic groups detected in Spain and environmental variables using Chi-squared or t-tests in R.

While the analyses described in the previous paragraph took advantage of the thinned datasets, the unthinned ones were used to compute global and per-locus F_ST_ (Weir & Cockerham, 1984) between genetic groups (in Spain) or transect ends (in Sweden, defined as positions before 37 and after 92 m as in Koch *et al*., 2022), without imputation for missing values, using the R package hierfstat v. 0.5-7 (Goudet, 2005). At a larger geographic scale, divergence between countries was explored using the joint thinned datasets through PCA and DAPC as described above.

Chromosomal inversions are known to contribute to ecotype differentiation in both Sweden and Spain (Faria, Chaube, *et al*., 2019; Koch *et al*., 2022; Morales *et al*., 2019; Westram *et al*., 2021, 2023). To investigate patterns of differentiation along the genome and identify additional chromosomal inversions in Spain that might or might not overlap with known ones, the unthinned datasets were analysed using two approaches. First, per locus F_ST_ values between genetic clusters were plotted along each linkage group (LG hereafter) to produce Manhattan plots using custom scripts in R. Genomic inversions differing in frequency between clusters are expected to generate blocks of higher differentiation compared to non-inverted regions (Le Moan *et al*., 2024; Mérot, 2020). To test for differences in F_ST_ between inverted regions and collinear regions we used a permutation approach. For each LG, we computed the size of inversions in number of SNPs and a block of equal size was randomly positioned on the LG. Then, the difference between the average F_ST_ in that block and the average F_ST_ in the rest of the LG was computed. This procedure was repeated 1000 times and the observed F_ST_ difference between the inversion position and the rest of the LG was compared to the distribution of the permuted F_ST_ differences. If more than one inversion was present in a LG and the inversions overlapped, we considered them as one block in the permutation procedure. If the inversions did not overlap, two blocks were shuffled on the LG without being allowed to overlap and the F_ST_ difference was computed between the average of the two blocks and the rest of the LG. When the observed F_ST_ difference was higher than the 95^th^ quantile of the distribution, we considered the F_ST_ difference between the inversions and the collinear regions to be significant. Second, we computed PCAs by map position using the same procedure illustrated above and custom scripts to extract the SNPs falling on each linkage map position. PC1 scores for each individual and map position were extracted and transformed by reversing the sign of the score when needed so that individuals belonging to the same genetic group always clustered on the same side of the PC1 axis. PC1 scores were then plotted along each LG and individuals were coloured according to their genetic group (Crab or Wave). The PCAs by map position were used to detect inversions without using prior knowledge regarding their positions in the genome. However, in some cases this approach failed to identify inversions in the positions of the original Swedish inversions, probably due to a lack of power. Therefore, additional PCAs were computed, in both Spain and Sweden independently, for each genomic region known to carry inversions in *L. saxatilis* and/or its sister species *L. arcana* (Reeve *et al*., 2023). We refer to these PCAs as ‘PCAs by inversion’. Typically, a single chromosomal inversion would result in the presence of three distinct clusters on a PCA plot, with the two most distant groups comprising individuals that are homozygotes for one arrangement or the other and the cluster in between comprising individuals carrying both arrangements (Reeve *et al*., 2023). In the presence of overlapping inversions, the PCA plot displays a characteristic triangular pattern of six clusters with the three apical groups comprising homokaryotype individuals (Mérot, 2020). An inversion was considered to be present when both the Manhattan plot and PCA by map position or by inversion displayed an inversion signal. Furthermore, to explore if the two ecotypes shared the same inversion arrangement in the two countries, loci that were present in both the Swedish and Spanish datasets were extracted and individuals from Sweden were projected into the PCA space defined by the Spanish individuals using the suprow function in the *ade4* R package v. 1,7-18 (Dray & Dufour, 2007), labelled projected PCA hereafter.

According to the patterns observed in both the independent and projected PCAs by inversion, these structural variants were classified in two categories: (i) regions showing a clear and shared inversion pattern in both Spain and Sweden, and (ii) regions showing different patterns between countries or no obvious inversion signal. In the former cases, we computed arrangement frequencies in each country using the country-specific PCA for each single inversion. In both countries, association between arrangements and genetic group (Spain) or end of the transect (Sweden) were tested using a Chi-squared test in the rstatix R-package v. 0.7.2 (Kassambara, 2023) and we used the projected output to determine whether the same arrangement had a higher frequency in the same ecotype in both countries. To further explore the relationship between inversion genotype and shore position within the heterogeneous Spanish Crab ecotype, the correlation between the frequency of the arrangement most abundant in the Wave ecotype and shore position was tested in the Spanish Crab ecotype using a Kendall rank test in rstatix. For simple inversions, PC2 was associated with within-arrangement differentiation between the ecotypes. In some cases, homozygote individuals for the arrangement most abundant in the Wave ecotype were split into two distinct sub-clusters along PC2 corresponding to the two ecotypes. In most cases, those sub-clusters showed some overlap. We investigated the presence of gene flow between ecotypes within arrangement by plotting the PC2 scores against shore positions for individuals carrying the arrangement most abundant in the Wave ecotype, using the ggplot2 R package v. 3.4.4 (Wickham, 2016).

To investigate the effect of chromosomal inversions on the observed patterns of divergence between ecotypes within countries, the genetic analyses described above were repeated excluding SNPs falling within known inversions (Faria, Chaube, *et al*., 2019; Koch *et al*., 2022; Reeve *et al*., 2023; Westram *et al*., 2021) from the original Swedish and Spanish thinned datasets using vcftools.

Consistency among the randomly-sampled subsets of SNPs and reads was tested using Kendall’s coefficient of concordance, W, computed using only the complete cases of the rankings with the R package DescTools v. 0.99.48 (Kendall, 1948; Signorell, 2023).

### Phenotypic analyses

Parallel morphological differentiation of ecotypes in Sweden and Spain has been described previously (Butlin *et al*., 2014; K. Johannesson *et al*., 2010a). Continuous variation in multiple phenotypic traits across the Swedish transect has also been described previously in genotyped individuals (Koch *et al*., 2021, 2022; Larsson *et al*., 2020, Westram *et al*. 2021). Here, we focused on quantitative phenotypic divergence in the Spanish transects, where the following traits were recorded in each snail: sex, wet weight, shell thickness (using NeoteckDTI Digital Dial Indicator Probe, 0.001mm resolution at the widest point of aperture and average over three measures), shell ridging (presence/absence), shell striping (presence/absence), and boldness. To measure boldness, snails were disturbed to induce retraction and time until they emerged again (out time) and until they got back on the foot (crawl time) were recorded. This test was repeated three times for each snail and the average score for both out and crawl times between the three trials was used for subsequent analyses. Scores were attributed according to the out or crawl times in minutes, using the following categories for the two measures, respectively: [0,<1], score=1; [0-1,1-5]=2; [1-5,5-10]=3; [5-10,10-15]=4; [10-15, >15]=5. Each trial was stopped after 15 minutes independently of the snail’s response. Details of this method deviate slightly from previous studies (Koch *et al*., 2021) but result in a similar distinction between ‘bold’ (low score) and ‘wary’ (high score) behaviours.

Shell length and growth parameters were obtained from standardized pictures of each snail. These pictures were analysed using *ShellShaper* (Larsson *et al*., 2020) and aperture position parameters (r0_scaled=r0/shell_length, z0_scaled=z0/shell_length), aperture shape (extension factor c0/a0), aperture size (a0_scaled=a0/shell_length), the relative shell thickness (thickness/a0), height growth (log_gh), width growth (log_gw) and convexity (abs(log_gh-log_gw)) were computed as in previous studies (Koch *et al*., 2021; Larsson *et al*., 2020). For each phenotypic trait, we tested differences between the two genetic groups identified using the clustering analysis described above. Additionally, multivariate patterns were investigated through a PCA using the *prcomp* function from the R package *stats* v. 4.3.0 and the following phenotypes: a0_scaled, r0_scaled, z0_scaled, log_gh, log_gw, relative thickness, weight, thickness, and shell length. Finally, to investigate whether genetic groups and/or shore position had an effect on phenotypes, linear models were built using either phenotypic PCs or individual phenotypic variables as response variable.

To compare phenotypes between Sweden and Spain, we retrieved phenotypic data from Koch *et al*. (2022) for the “CZA”sampling of the same transect in Ramsö (Sweden) a few years earlier (Westram *et al*., 2021). That dataset included the following variables: shell length, wet weight, aperture size (a0_scaled), aperture position (r0_scaled), aperture shape (c0/a0), height growth (gh), width growth (gw) and relative thickness. All these variables were scaled in each country and used in a PCA including all individuals from both CZA and Spain. In CZA, individuals found in the boulder habitat at least 15 meters away from the boulder-rock transition were considered Crab ecotype, and individuals found in the rocky habitats at least 40 meters away from the boulder-rock transition were considered Wave ecotype, following Koch *et al*. (2022). Finally, average relative differences between ecotypes were computed as the difference between the Crab average and the Wave average divided by the Crab average for each variable both in Sweden and Spain and bootstrap (10,000 iterations) confidence intervals were computed in R.

## Results

### Distinct patterns of genetic divergence in Sweden and Spain

We explored genetic divergence along the transects spanning the Crab-hybrid-Wave axis in Sweden (boulder field to rocky headland) and Spain (high shore to low shore) using low-coverage whole-genome resequencing data in 283 snails. The Spanish transects did not show seasonal patterns (Fig. S1 in Supplementary Information), thus the samples from spring and autumn were merged in the subsequent steps. A total of 311,549 (unthinned datasets) and 21,250 (thinned datasets) SNPs were obtained in Sweden, while the Spanish unthinned and thinned data included 339,614 and 21,212 SNPs, respectively. The joint thinned datasets, including SNPs polymorphic both within and among countries used to investigate overall divergence between Sweden and Spain, included 7,577 SNPs. As our filters were quite stringent, sites had to have reads in more than 50% of all 283 individuals to be included in this joint dataset. This resulted in a lower number of loci but unbiased against divergent sites between countries. We first describe genome-wide analyses, including both collinear loci and chromosomal inversions. Thereafter, we present results without inversions.

Sweden and Spain showed distinct patterns of genetic differentiation. In Sweden, snails at the Crab and Wave ends of the transect were distinguished by PC1 (3% of variation), but with a continuous range of intermediates distributed clinally along the transect so that cluster analysis identified only a single genetic group (Fig. 1, Fig. S2). This pattern was consistent among the random SNP and read subsets (Table S1) and in line with previous studies (Westram *et al*., 2018, 2021). In contrast to the Swedish population, the Spanish snails did not show a clinal pattern along the transect but instead formed two genetically distinct groups: one more genetically variable and spanning almost all of the transect (here referred to as the Crab ecotype for consistency with previous studies, see Discussion), while the other one was more homogeneous and mostly localized in the lower part of the shore (Wave ecotype). No intermediate genotype was observed between these two groups (Fig. 1, S2-S3). However, some Spanish Crab individuals showed a hybrid index value close to 0.6, indicating admixture and significant contributions of alleles typical of the Wave ecotype in their genome, while individuals of the Wave ecotype had little or no evidence of Crab ancestry. None of the analysed Spanish snails had a Crab ancestry proportion between 0.1 and 0.6, or a hybrid index between 0.2 and 0.55, whereas such individuals were common in the Swedish transect (Fig. 1, Table S2). These results were consistent among the random SNP and read replicates (Fig. S3, Table S1), including the assigned group membership, i.e., the same individuals were consistently classified as belonging to the Crab or Wave ecotype. Within-group PC1 scores were highly correlated among the random SNP and read subsets (range: 0.7999 - 0.9965, Fig. S4). The two ecotypes in Spain exhibited different associations with habitat features (Fig. S5). The Wave ecotype was more often associated with the goose barnacle *Pollicipes*. The *Mytilus* zone defined the overlap between the Crab and Wave ecotypes. Overall, the Wave ecotype was confined to the lower shore while the Crab ecotype was distributed over the whole vertical transect although at low densities in the lowest positions (Fig.1, Fig. S5).

**Figure 1.**
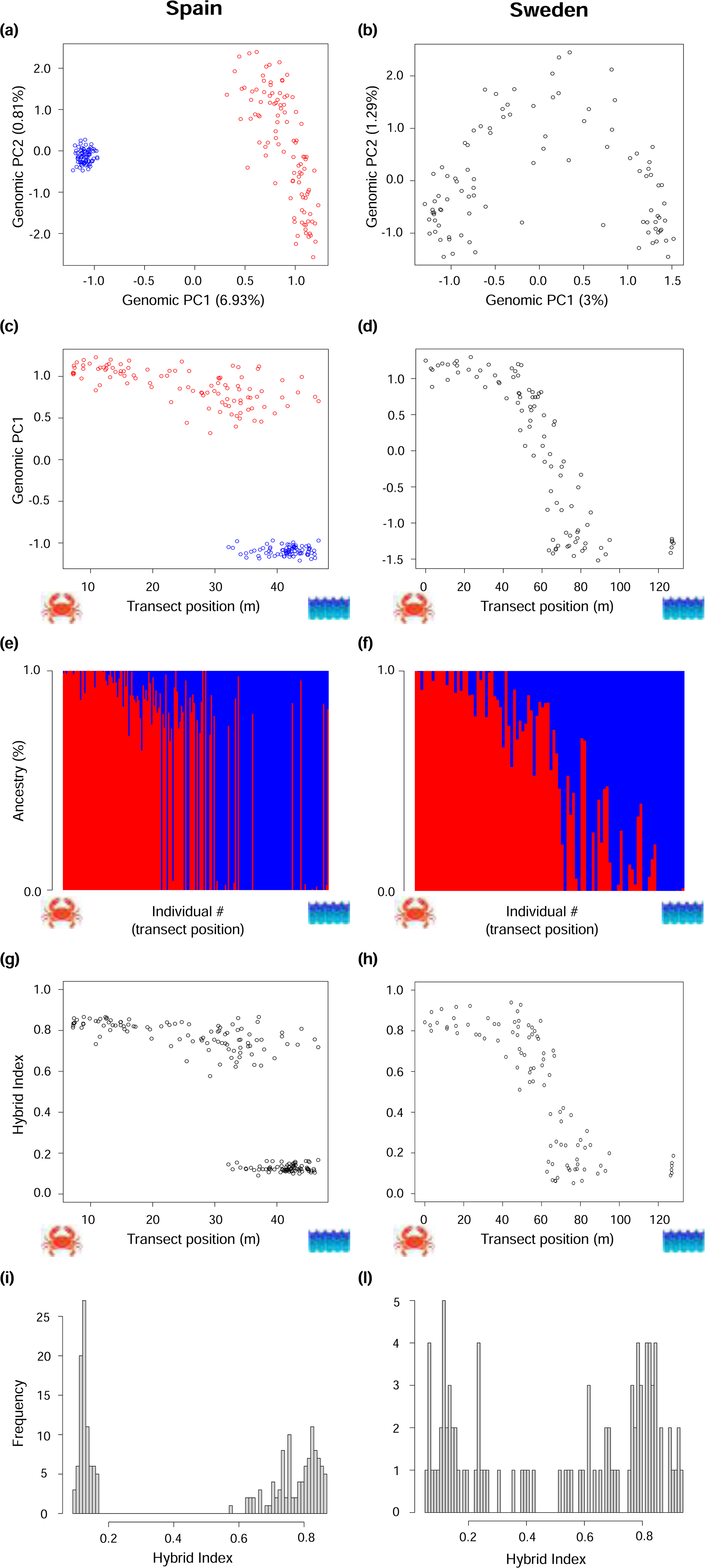
Patterns of genomic divergence in Spain (**a, c, e, g, i**) and Sweden (**b, d, f, h, l**) as shown by the genome-wide PCA (**a, b, c, d**), admixture (**e, f**) and the Hybrid Index (**g-l**). Transect positions closer to zero correspond to the high shore (Spain) or boulder field (Sweden), where the Crab ecotype is typically found, while higher values represent the low shore (Spain) or rocky headland (Sweden), that typically hosts the Wave ecotype. Snails in the admixture plots (e, f) were ordered according to their position along shore. In Sweden, a sampling gap occurred at the transect positions 90-120 as seen in the plot of PCA1 and Hybrid Index along the shore (d, h) and a single genetic cluster was identified statistically but two groups were forced in the admixture plot (f) to facilitate comparisons with Spain. The Spanish Crab and Wave ecotype are coloured in red and blue, respectively.

At a larger spatial scale, the joint analyses consistently identified three genetic groups: Sweden and the two Spanish ecotypes. Most of the differentiation was explained by geographic separation between the two regions, followed by the Crab-Wave axis in Spain, and the divergence between the two Spanish ecotypes was larger than divergence in the whole Swedish transect (Fig. S6-S7).

### Differentiation along the genome and chromosomal inversions

The genome-wide average differentiation between the two genetic groups was higher in Spain compared to the ends of the transect in Sweden (global F_ST_ values of 0.11 and 0.08, respectively, Table S3). The average F_ST_ by LG was higher in Sweden than in Spain in LG8 and LG15; similar between the two countries for LG7, LG11, LG13 and LG16; and higher in Spain than in Sweden in all the other LGs (Table S3).

Some regions displayed higher F_ST_ in Sweden compared to Spain, while other regions presented high differentiation in both countries, both in the form of islands or narrow peaks of high differentiation relative to adjacent regions (Fig. 2, S8-S9). A total of 53 contigs (4,711,312 bp) exhibited high differentiation in both countries (average F_ST_ per contig > 0.3 in both countries, Fig. S10) accounting for approximately 43% (53/122) and 11% (53/476) of contigs with F_ST_ values exceeding 0.3 in Sweden and Spain, respectively. Highly differentiated genomic regions in both countries were located on LG1, LG2, LG3, LG5, LG6, LG8, LG9, LG12, LG14 and LG17, with LG6 and LG14 containing 26 and 10 of these contigs, respectively (Table S4). Forty-one (3,468,966 bp) of the 53 shared highly-differentiated contigs were located in known inversions that were polymorphic between ecotypes in Sweden (Faria, Chaube, *et al*., 2019; Hearn *et al*., 2022; Westram *et al*., 2021) on LG6, LG9, LG12 and LG14.

**Figure 2.**
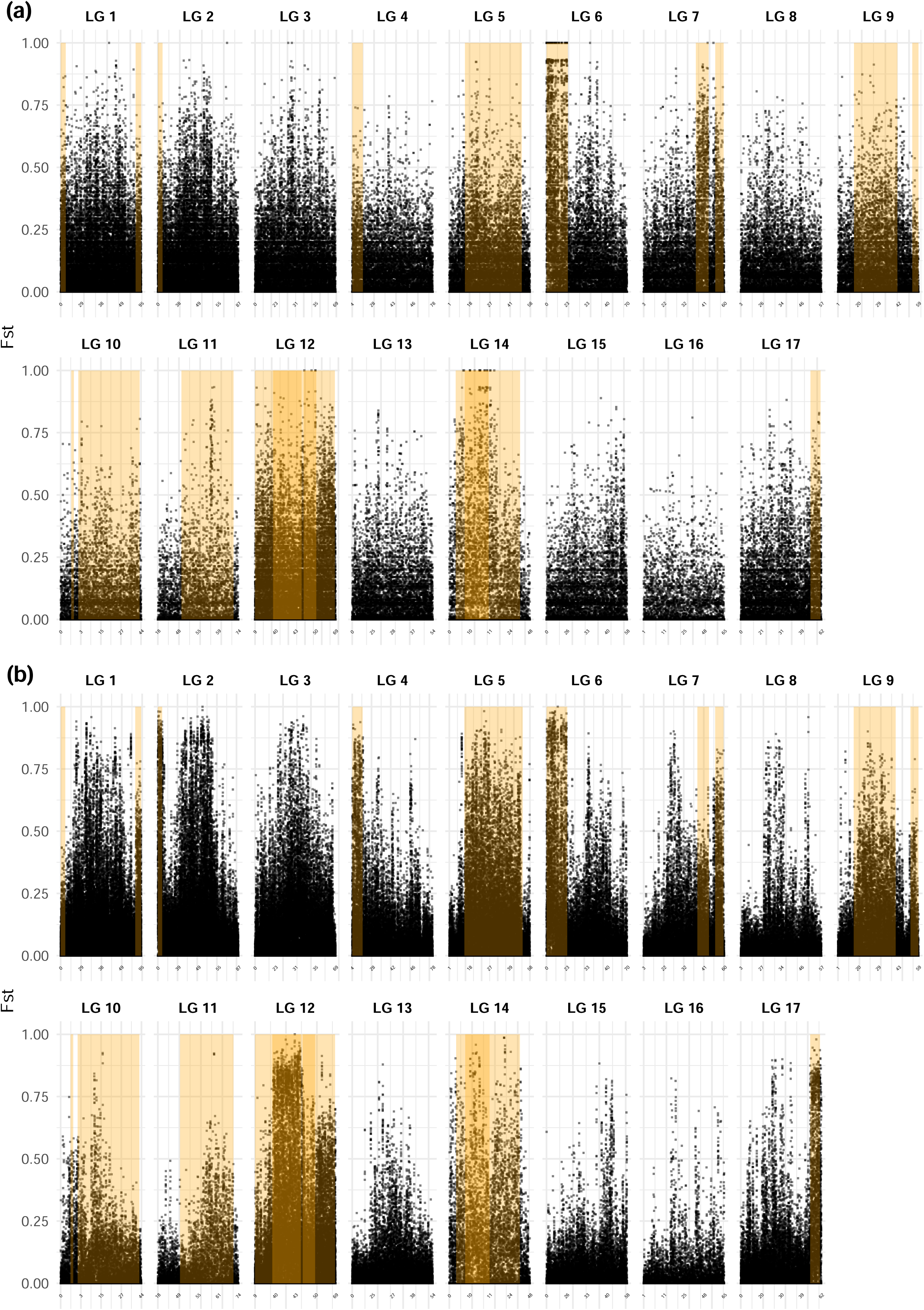
Manhattan plots of per locus F_ST_ values along the genome in Sweden (**a**) and Spain (**b**). The linkage groups and positions in centimorgans are indicated at the top and the bottom, respectively. Chromosomal inversions identified in previous studies are highlighted in orange. Overlapping rectangles indicate the presence of overlapping inversions.

To investigate the presence of inversions along the genome, we used Manhattan plots and PCA by map position or by inversion along each of the 17 LGs. Multiple chromosomal inversions were identified in both countries in the same positions across the genome. Inversion patterns were congruent among methods (F_ST_ and PCA by map positions) and generally more pronounced in Spain than in Sweden, with Manhattan plots showing blocks of high F_ST_ that were more differentiated from the background and PCA plots showing more distinct clusters (Fig. 2, S8-S9,S11-S12). While the map-based approaches produced inconclusive patterns for LGC5.1, LGC6,1/2, LGC9.2, LGC10.1, LGC10.2, LGC11.1, LGC12.1/2/3/4 and LGC14.1/2, the PCAs by inversion confirmed the presence of these inversions in our dataset (Fig. 3, S13).

**Figure 3.**
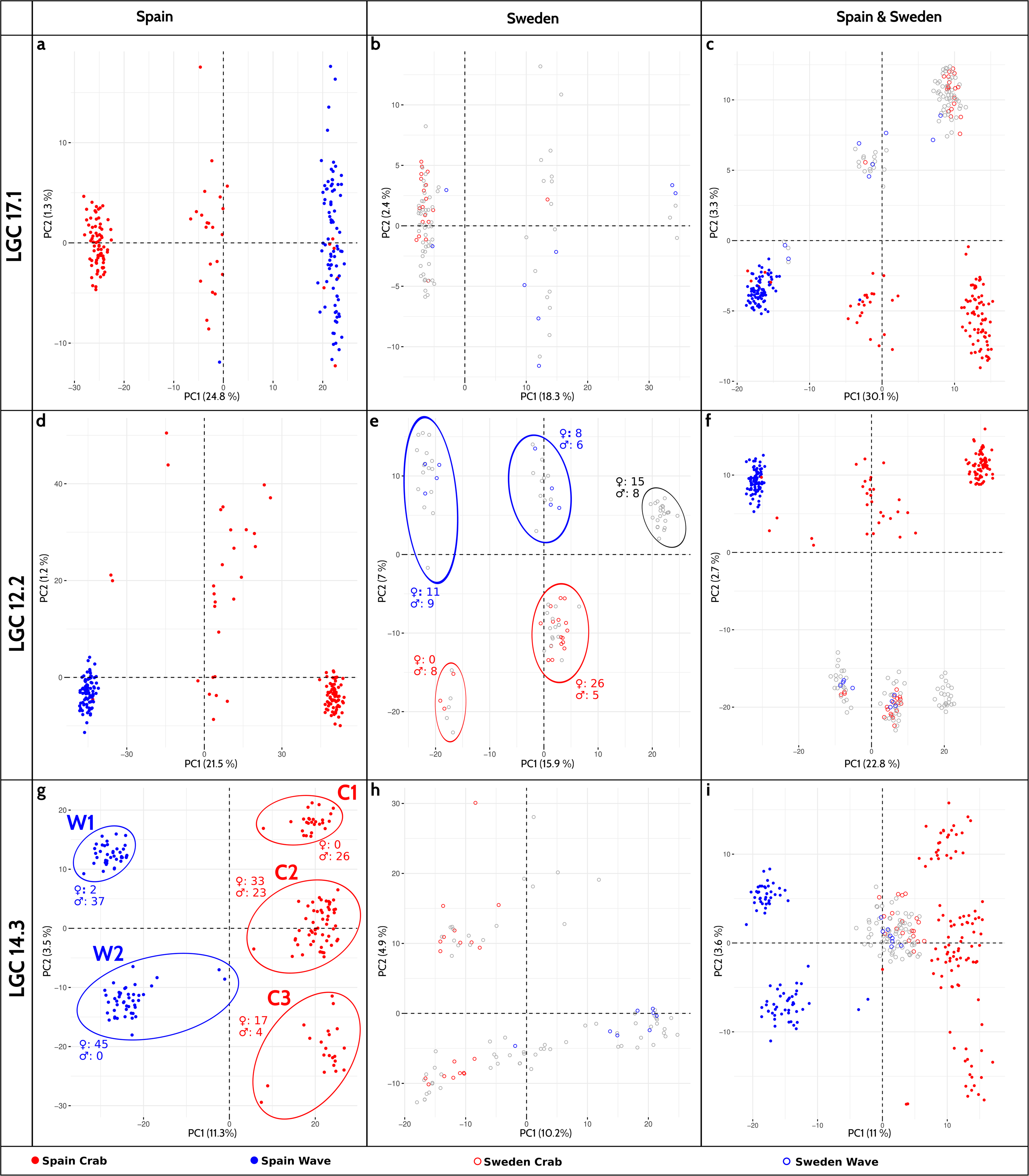
Genetic PCAs by inverted region in Spain (**a,d,g**), Sweden (**b,e,h**), and both countries (suprow approach; **c,f,i**) showing examples of inversions with similar (simple: LGC17.1, **a,b,c;** or complex: LGC12.2; **d,e,f)** or contrasting (LGC14.3; **g,h,i**) patterns between countries. Filled dots represent the Spanish individuals, empty circles the Swedish ones. The Crab and Wave ecotype (Spain) or ends of the transect (Sweden) are denoted in red and blue, respectively, while grey circles represent individuals in or close to the Swedish contact zone (middle portion of the transect).

Overall, a total of 19 inversions were established in our datasets, 14 shared between the two countries, 4 unique to Spain and 1 unique to Sweden (Table 1, Fig. 3, S13). Compared to the set of previously known inversions (Reeve *et al*., 2023), our study did not detect any polymorphic rearrangements in LGC10, where two have been recorded elsewhere, and did not detect any new rearrangements. Inverted regions were generally significantly more differentiated than collinear regions, with similar patterns in Spain and Sweden (Fig. S14-S15). However, in a few LGs, the differentiation gap between inverted and collinear regions was higher in Spain than in Sweden (Fig. S8-S9, S14-S15), SI: Differentiation between inverted and collinear regions).

**Table 1.**
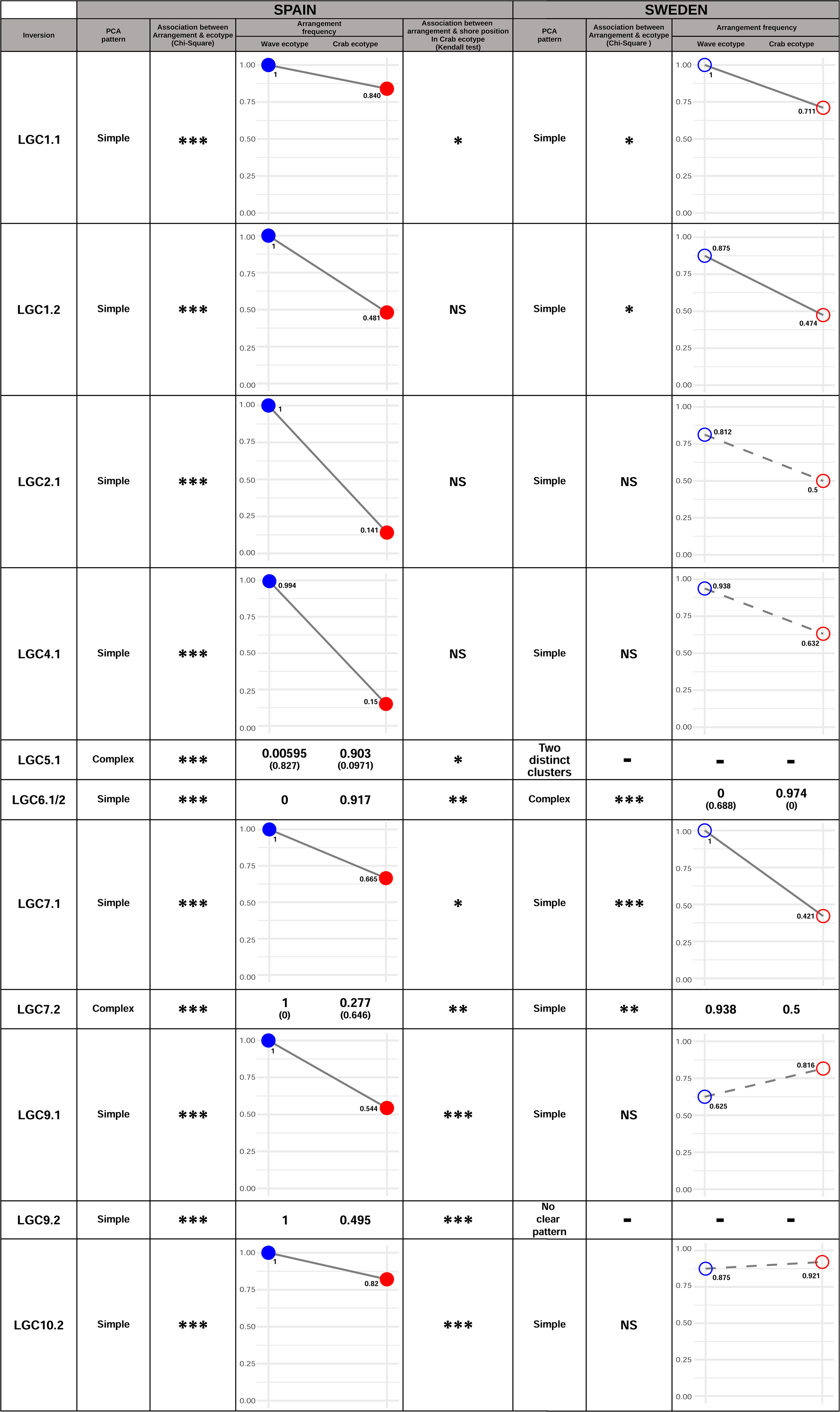
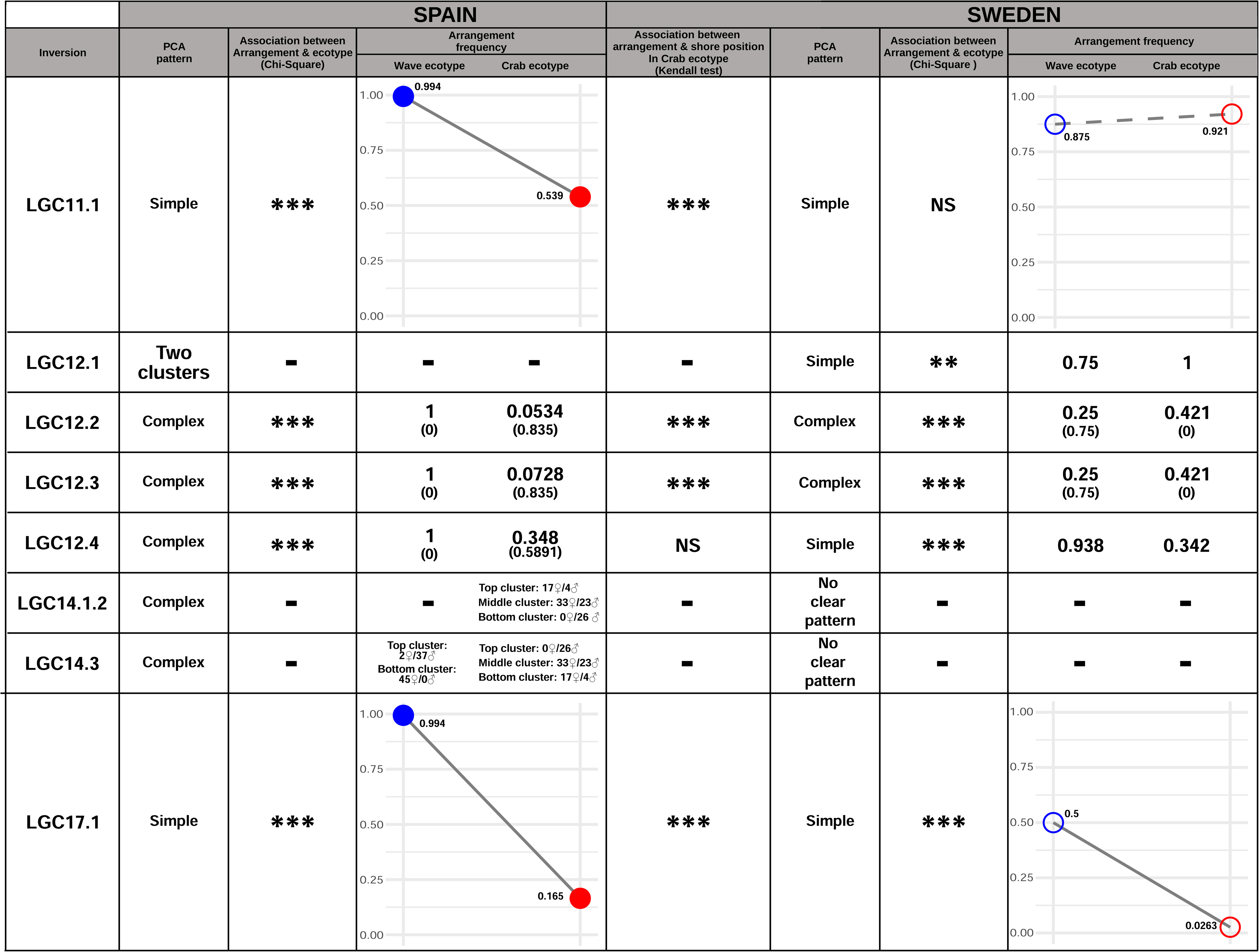
Inversions identified in Spain and Sweden through the PCA by inversion approach in this study. When the projected PCAs (Fig. S13) showed clear inversion patterns, we plotted the frequencies of the same arrangement in both countries. In the miniplots, solid lines indicate statistically significant arrangement frequency differences between ecotypes, while dashed lines represent non-significant differences. When the projected PCAs did not allow identifying the same arrangement across countries, we reported the frequencies of the most abundant arrangement in each country. For complex inversions, with three arrangements present, the frequency of the next most abundant arrangement is given in parentheses. In the Spanish LG14, the number of male (♂) and female (♀) individuals in each ecotype is reported. Dashes denote untested cases due to unclear inversion signals. The statistical significance level is indicated using NS (P-value > 0.05), * (P-value < 0.05), ** (P-value < 0.01), *** (P-value < 0.001).

A mix of simple and complex inversions was detected, with patterns not always consistent between Spain and Sweden (Fig. 3, Table 1, SI: Inversion results). In Spain, arrangement frequencies were associated with ecotypes for all inversions, whereas this was true for only a subset of inversions in Sweden. In all except three cases (LGC9.1, LGC10.2 and LGC11.1, where arrangement frequency differences between ecotypes were not significant in Sweden) the arrangement with higher frequency in Wave was the same in both countries. In Spain, for many inversions, one arrangement was fixed in the Wave ecotype while arrangement frequencies varied with shore position in the Crab ecotype, with Crab individuals in the low shore more likely to carry the Wave arrangement (Fig. S16). In some cases, the Crab and Wave ecotype homozygote individuals carrying the Wave arrangement formed distinct clusters in the PCA-by-inversion analyses, along PC2. This suggests that, while both Crab and Wave individuals can carry the Wave arrangement, the content of this arrangement differs between ecotypes, indicating low or absent gene flow. However, for other inversions Crab and Wave individuals were spread along PC2 and for LGC1.2, the arrangement found in Crab individuals was more Wave-like in the lower part of the shore (Fig. S17), suggesting that gene flow might occur in this particular arrangement between the Crab and Wave ecotypes. In Sweden, arrangements for LGC12.2 were associated with sex in the Crab ecotype, as previously reported (Hearn *et al*. 2022), but this was not true in Spain. Instead, arrangements in LG14 showed associations with sex (Fig. 3, Fig. S13) suggesting that the sex determining locus is on different chromosomes in the two countries. More detailed descriptions of these inversion patterns are provided in Supplementary Information: Inversion results.

To investigate the contribution of chromosomal inversions to genetic divergence along transects, the genetic analyses illustrated in Fig. 1, S2, and S4 described above were repeated using the thinned datasets and removing the SNPs in known inversions. In Sweden, clinal patterns remained consistent after removing the positions within known inversions (Fig. S18-S19, Table S2). A similar trend was observed in Spain, where no substantial differences were detected between the patterns with and without inversions: both showed two distinct genetic groups without intermediate genotypes, and the down-shore variation was still present within the Crab ecotype (Fig.1, S2-S3, S18-S19). The Wave ecotype was less homogeneous, and the percentage of variance attributed to the divergence between the two genetic groups (Wave and Crab) in PC1 was slightly lower (5.5%) compared to the analyses with inversions (6.93%, Fig. 1, S18). As in the analyses with inversions, results were consistent among the random subsets, which showed the same individual assignment to the Crab or Wave ecotype and a high correlation of the within-ecotype PC1 scores (Fig. S20, Table S1). After removing inversions, genome-wide average F_ST_ decreased both in Spain and Sweden (Table S3). The genome-wide variability of F_ST_ (standard deviation of F_ST_ in the genome) was higher in Spain than in Sweden and decreased in both countries with the removal of the inversions (from 0.162 to 0.138 in Spain and from 0.137 to 0.121 in Sweden). It remained high in Spain and Sweden, indicating heterogeneity in the barrier to gene flow over and above the direct contribution of inversions.

### Phenotypic divergence

Analyses of individual traits in 185 Spanish snails revealed significant phenotypic differences between the two genetic groups. Some phenotypes could not be measured in all snails (see sample sizes in Fig. S21). Overall, the Crab snails were bigger, heavier, and possessed thicker shells with lower height and width growth, a higher aperture height, a smaller aperture size and higher size-independent relative thickness compared to Wave (Fig. S21). Most of the Crab shells (93%) displayed ridges while they were present only in a single Wave snail (Fig. S21). In Sweden, ridging is observed in Crab but it is much less pronounced compared to Spain and infrequently present in Wave (Castillo *et al*., 2023). Striped shells, a phenotype never present in Sweden, were observed in most snails except eleven individuals belonging to the Wave ecotype. Stripes were mostly black in the Crab snails, while they were both black and brown in Wave (Fig. S21). Behavioural tests indicated that the Crab snails were more wary than the Wave ecotype (Fig. S21). In fact, more than half of the Crab snails took more than ten minutes to come out of their shell (median out boldness score of 4.33) whereas less than five minutes (median out boldness score of 1.33) were needed for most of the Wave individuals (Fig. S21). Using data from Koch *et al.* (2022) for the CZA transect, relative differences between average individual phenotypic traits in Crab and Wave varied between Spain and Sweden. Except for aperture shape and aperture position in Spain, relative differences between ecotypes were all different from zero. Wherever differences between ecotypes were found in both countries, they were in the same direction. While relative differences between average wet weight and shell length were similar between ecotypes in Sweden and Spain, there were greater and significant differences in average aperture size, aperture position, aperture shape, relative thickness, width growth and height growth between ecotypes in Sweden compared to Spain (Fig. S22).

The multivariate analysis indicated a more continuous variation in phenotypic than genomic data in the Spanish samples, and partial overlap between the Crab and Wave ecotypes (Fig. 4). The first axis of the PCA, accounting for around 51% of the observed variation, was highly correlated with shell length parameters while the second axis, describing around 19% of variance, was highly correlated with the aperture position (r0 scaled and z0 scaled, Fig. 4a). Overall, the two ecotypes were phenotypically different. However, one individual (a small Crab snail) showed discordance between its genetic background and phenotype, i.e., phenotypic features that were typical of the other ecotype. The Crab snails showed a down-shore cline, as in the genomic analyses, with some individuals overlapping spatially and phenotypically with the Wave ones: individuals located closer to the lower shore exhibited more Wave-like phenotypic characteristics (Fig. 4b).

**Figure 4.**
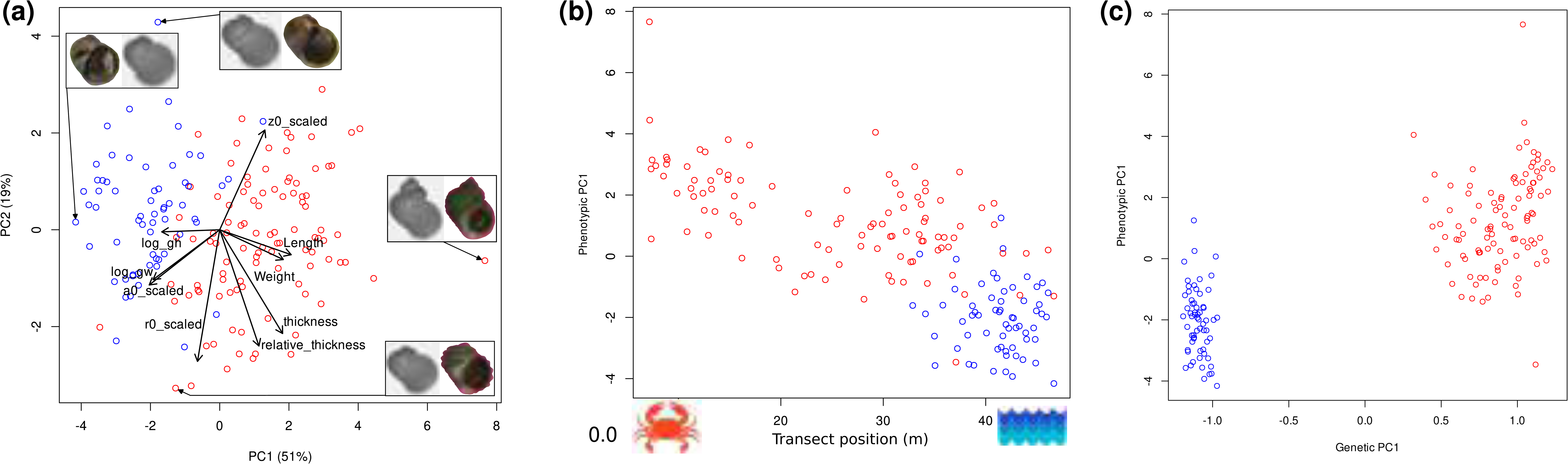
Multivariate phenotypic divergence in Spain as shown by the first two PCA axis (**a**), PC1 along the transect (**b**), and its relation to genome-wide differentiation (**c**). Transect positions closer to zero correspond to the high shore while higher values represent the low shore. The Crab and Wave ecotype are coloured in red and blue, respectively

Linear models revealed that both the genetic group and transect position had a significant effect on the PC1 phenotype. However, a closer look at the patterns within groups indicated that shore position had a significant effect on the PC1 phenotype only within Crab (Fig. S23, Table S5, Table S6). The PC2 phenotype was only influenced by the genetic group. The genetic group had a significant effect on every individual phenotypic trait except aperture shape, convexity and z0 scaled. The transect position had a significant effect on most individual phenotypic traits (Table S5).

The joint phenotypic PCA, including the common subset of traits from both the Spanish snails and the Swedish CZA dataset from Koch *et al*. (2022), also showed continuous phenotypic variation between individuals (Fig. S24). Divergence between Crab and Wave ecotypes was very similar in direction in both countries but higher in Sweden than in Spain. Ecotype difference was mostly carried by the first axis which accounted for around 66% of the total variation and was highly correlated with aperture size, aperture shape, and height growth. Wave ecotype snails appeared more similar between countries than Crab ecotype snails and the Crab ecotype in Spain was, on average, more “Wave-like” than the Crab ecotype in Sweden.

## Discussion

*Littorina saxatilis* in Sweden and Spain illustrate the evolution of replicated reproductive isolation in two independent lineages of the same species, i.e., parallel speciation (Schluter & Nagel, 1995). The two systems analysed here have a relatively recent common ancestry (50-278ky, Carvalho, Faria, *et al*., 2023; Panova *et al*., 2011)), and share some genomic patterns of differentiation (Morales *et al*., 2019; Reeve *et al*., 2023). Hence the speciation processes of both should be promoted or limited by roughly the same genomic potentials and constraints. Moreover, there is evidence that divergence into Crab and Wave ecotypes has evolved without extended periods of isolation, that is, mostly in the presence of gene flow in both countries (Butlin *et al*., 2014; Carvalho, Faria, *et al*., 2023). These natural replicates offer an outstanding opportunity to investigate the impact of different factors contributing to reproductive barriers in the two countries.

While much of the genetic and phenotypic structure of this Swedish contact zone was known from earlier studies (Westram *et al*., 2018, 2021), our understanding of the Spanish ecotypes was based on reduced representations of the genome and analysis of parental and hybrid groups identified from phenotypic traits, rather than genetically-assigned individuals(K. Johannesson *et al*., 1993, 1995; Kess *et al*., 2018; Rolán-Alvarez *et al*., 1997). This could not reveal the full pattern of differentiation, either spatially or genetically, and so restricted understanding of gene flow. Our dense and, with respect to phenotype, random sampling in a Spanish contact zone in Centinela revealed two distinct genetic clusters and no F1 hybrid individuals or potential backcrosses to the Wave ecotype. A lack of admixture, consistent with our observations, was previously described in a SNP-based study with targeted sampling of phenotypically intermediate individuals (Kess *et al*., 2018). The data we provide here show a homogeneous Wave ecotype, very similar in phenotype to the Swedish Wave, and in Spain found exclusively in the lowest part of the shore. Conversely, the Spanish Crab ecotype, phenotypically resembling the Swedish Crab one, is mostly present in the high shore but is also distributed throughout the shore and overlaps spatially (but not genetically) with the Wave ecotype in the

lowest zone. Previous work shows that the pattern observed in the Swedish site studied here is repeated across multiple sites (Westram *et al*., 2021)(. Although no other sites have been studied in a similar way in Spain, the results of Kess *et al*. (2018) for three other sites in North Western Spain strongly suggest the presence of distinct genetic groups with few intermediates and measurements of assortative mating between ecotypes at multiple locations have demonstrated isolation indices between 0.64 and 1.0, including Centinela that was one of the sites with the lowest value (Rolán-Alvarez *et al*., 1999). Our insights suggest that, while the two regions share many similarities, they also show significant differences and represent different points on the speciation continuum.

The systems have evolved parallel phenotypic divergence resulting in similar Crab and Wave ecotypes driven by strong divergent natural selection established by high predation from crabs in one environment and strong physical stress from wave action in the other micro-habitat (Janson, 1983; K. Johannesson *et al*., 2010b; Koch *et al*., 2022; Rolán-Alvarez *et al*., 1997). Snails from these two sites located in distant countries share a similar genomic architecture that shows high levels of differentiation between genetic groups or ends of transects (this study, Koch *et al*., 2022), including genomic loci located within chromosomal inversions (this study, Morales *et al*., 2019; Reeve *et al*., 2023). Moreover, we here show that most of these inversions present the same arrangement at higher frequency in the same ecotype (Crab/Wave) in both countries, indicating common sets of adaptive alleles within inversions that might have a shared origin. Inversions are key components in divergence and local adaptation especially under gene flow (Barth *et al*., 2017; Faria, Johannesson, *et al*., 2019; Kirkpatrick & Barton, 2006; Wellenreuther & Bernatchez, 2018), and they host genes contributing to traits under selection in *L. saxatilis,* at least in Sweden (Koch *et al*., 2021, 2022). Shared genetic loci may have allowed for allele reuse in response to similar environmental pressures and kick-started the adaptive and diversifying processes in parallel, while reduced recombination has contributed to overcome the homogenizing effects of gene flow and aided the establishment of separate evolutionary paths in the face of permeable reproductive barriers. Furthermore, size-assortative mating seems to be present in both locations (K. Johannesson *et al*., 1995; Perini *et al*., 2020) and provides a potential additional barrier to gene flow, although, as discussed below, this might not be more than marginally important. Interestingly, inversions alone did not fully explain the two distinct genetic clusters observed in Spain, as is also true in Sweden (this study, Koch *et al*., 2022; Westram *et al*., 2021). In fact, the two groups in Spain remained clearly discrete after removing inversions, suggesting that gene flow between the Spanish Crab and Wave ecotypes is highly restricted throughout the genome. Yet, genome-wide divergence between ecotypes is low in both countries, potentially due to their recent origin or ongoing gene flow, albeit weak.

However, some country-specific patterns emerged in the two target sites. Our results support the presence of two private inversions in these Spainish transects (LGC5.1 and LGC9.2), previously identified as “new putative inversions” (Faria, Chaube, *et al*., 2019; Reeve *et al*., 2023) that were not reported in the Swedish site, hinting that the contribution of structural variants to evolution might differ between populations. Sex determination is strongly though not perfectly linked to inversions in both lineages, as previously reported in several organisms (Bachtrog, 2013; Blumer *et al*., 2024; Peichel *et al*., 2020). Remarkably, our findings suggest that sex determination in Spain and Sweden did not involve the same loci, a rare example of intra-specific variation in the genetic determination of sex. In Sweden, LGC12.2 and LGC12.3 showed a strong association with sex in Crab (this study, Hearn *et al*., 2022; Koch *et al*., 2021). Conversely, LGC14.1/2 and LGC14.3 are related to sex in Centinela (Spain), with a particularly strong relationship displayed by Wave individuals in the latter inversion. Additionally, Wave in Sweden presents the highest abundance of the LGC9.1 inversion arrangement found in the Spanish Crab snails (this study, Morales *et al*., 2019). This locus is most likely implicated in local shore height-related adaptation to temperature and/or desiccation stress as Crab in Spain and Wave in Sweden are found higher on the shore than Wave in Spain and Crab in Sweden.

A key distinction between the two lineages is that reproductive isolation between Crab and Wave ecotypes is substantially stronger in Spain than in Sweden (this study, Kess & Boulding, 2019; Morales *et al*., 2019). The overall level of genome-wide divergence between ends of transects or genetic groups is lower in the site in Sweden than Spain. The Swedish contact zones show a continuous, unimodal pattern with frequent hybrids at the contact (this study, Westram *et al*., 2021); in contrast, the Spanish contact zone analysed here is genetically discrete, bimodal with no early hybrids and evidence of only weak, unidirectional gene flow. Chromosomal inversions showed more distinct and ecotype-associated differences in frequencies in Spain, with several arrangements fixed in the Wave ecotype, while most of them remain polymorphic at transect ends in Sweden (this study, Westram *et al*., 2021). In some Spanish inversions, distinct haplotypes in the Crab and Wave ecotype were observed within the same arrangement (“sub-clusters” in PCAs), indicating very low levels of gene exchange between these ecotypes for a substantial period and/or additional divergence due to some form of selection or genetic drift. Secondary contact between the Spanish ecotypes could also contribute to their divergence, although the available evidence does not support extended periods of previous isolation (Carvalho, Faria, *et al*., 2023) and short vicariance events are unlikely to contribute to strong reproductive barriers. Taken together, these differences indicate that the Swedish barriers to gene flow are considerably weaker than those in Spain; the Spanish ecotypes are closer to completion of speciation than the Swedish ones.

The Spanish snails belonging to the genetic Crab group become phenotypically more Wave-like in a down-shore gradient while still retaining their genetic group identity (this study, Rolán-Alvarez *et al*., 1997). The two Spanish genetic groups analysed here are mostly distinguished by shell ridging and colour especially in the overlap zone, as earlier described (Johannesson *et al.*, 1993, 1995; Rolán-Alvarez *et al.*, 1997, 1999). The Wave-like phenotypic appearance of the Crab individuals in the low shore is also accompanied by an increase of genetic similarity in collinear loci and in the frequency of inversion arrangements typically found in the Wave snails. This genetic and phenotypic convergence of the Crab and Wave ecotypes in the low shore is likely driven by microhabitat-related selection, supported by rare unidirectional weak gene flow from Wave to Crab, favouring some introgressed variants but working against others. Introgression would provide an important source of standing variation that can facilitate adaptation of Crab ecotype individuals to the Wave environment in the low shore Alternatively, and possibly complementarily, the more Wave-like background in Crab in the low shore could have emerged from ancestral polymorphisms facilitating local adaptation. The absence of fixed arrangements in any inversion in the Spanish Crab (contrasting with high rates of fixation in the Spanish Wave, Fig. S16) further lends support to the role of chromosomal inversions in adaptation of this group over a much more heterogeneous environment than Wave in Spain (but not in Sweden where environmental heterogeneity is greater in the Wave habitat). Which again raise a concern with the conventional naming of the Spanish genetic group distributed from high to low shore as a “Crab ecotype”.

Why do these two systems exhibit such different extents of reproductive isolation? We discuss five, non-mutually exclusive, potential explanations:

(1) The Swedish ecotypes are much younger than the Spanish ones (∼15ky and ∼57ky, respectively, Carvalho, Faria, *et al*., 2023). The Spanish populations survived the Pleistocene glaciations *in situ*, whereas the Swedish populations are the result of post-glacial colonisation (Bosso *et al*., 2022; Doellman *et al*., 2011; Panova *et al*., 2011; Stankowski *et al*., 2023). If the divergence is limited by available genetic variation, a longer time of divergence would have increased the chances for addition of mutations that increased the local adaptation and hence, by involving more loci, the barrier strength.
(2) The two contact zones are arranged differently in the shore space: a narrow and gradual environmental transition from boulder field to rocky headland in Sweden *versus* a large mosaic of barnacle and mussel patches in Spain. Individual life-time local cruising range is rather limited in this species (a few meters, Cruz *et al*., 2004; Erlandsson *et al*., 1998; Janson, 1983; Westram *et al*., 2018). In the Swedish contacts, most snails will never come close to the other ecotype, while in Spain the majority of snails live close to or in the contact zone. Selection for reinforcement of reproductive barriers would not take place except in the contact zone (Fernández-Meirama *et al*., 2022), and thus in Sweden only a small proportion of the population would be under selection for reinforcement of barriers. By contrast, in Spain many snails would be involved, and reinforcement alleles would be more likely to be established.
(3) Selection reinforcing prezygotic barriers to gene flow might be weak in Sweden. Preliminary data suggest that hybrid snails in Sweden are as fit in the intermediate habitat of the contact zone as pure ecotypes (Janson, 1983; Sadedin *et al*., 2009). The survival of the Spanish hybrids has not been measured in this study as they are rare. However, fertility data of hybrids from laboratory cross-breeding of Spanish Crab and Wave individuals show that F1 hybrids have very high rates of embryo abortion (60%, Figure S25) compared to what is found in Swedish hybrids present in contact zones (12%, K. Johannesson, 2020; Sá-Pinto *et al*., 2013). Low fertility of Spanish hybrids would represent an additional component of reproductive isolation in itself and would favour selection reinforcing prezygotic barriers in Spain.
(4) Size is a multiple-effect trait that may be conducive to coupling of reproductive barriers (Butlin & Smadja, 2018; Smadja & Butlin, 2011), and size-assortative mating is present in both Sweden and Spain (K. Johannesson *et al*., 1995; Perini *et al*., 2020). However, in Sweden where snail size changes gradually across the contact zone, the barrier effect of the size-assortment becomes minor due to the extensive number of hybrids (Perini *et al*., 2020). This might also apply to Spain, though less strongly, due to the convergence in size in the low shore. However, in Spain there is assortative mating caused by Crab and Wave being non-randomly distributed in the patchy environment and snails of divergent size overlapping in the mid-shore (Boulding *et al*., 2017; Rolán-Alvarez *et al*., 1999). Earlier experimental work based on phenotypic identification of ecotypes suggested micro-habitat choice causing assortative mating in the Spanish contact zone (Cruz *et al*., 2004; K. Johannesson *et al*., 1995), but these experiments need be repeated with individuals in which the mating pattern is confirmed by genotyping. In Sweden, such *habitat* choice would be difficult to achieve as the contact zone is more a continuous environmental transition than patchy, such that most snails experience only one habitat type (Janson, 1983; Westram *et al*., 2018).
(5) The Swedish genetic and phenotypic transitions are formed by mainly one selection gradient running from crab selection in boulder fields to wave selection on rocky cliffs. A second axis of selection runs from high to low shore with strong gradients in temperature and desiccation (Sokolova *et al*., 2000), but in Sweden this axis is perpendicular to the crab-wave selection axis. In Spain, the two selection gradients are parallel and act in synergy as both the temperature/desiccation axis and the Crab-Wave axis run from high shore to low shore. The distribution of inversion arrangements somewhat reflects the effects of the different selection axes in Sweden and Spain. In six of the inversions that showed differences between ecotypes in both Spain and Sweden, the same arrangement was more frequent in Wave than Crab in both countries (e.g., inversions on linkage groups LG6, LG12 and LG14). This is consistent with previous findings (Morales *et al*., 2019), and genes involved in shell traits that discriminate between Crab and Wave ecotypes are present in these inversions (Koch *et al*., 2021, 2022). However, one inversion (LGC9.1) showed contrasting arrangement frequencies between Crab and Wave ecotypes in the two countries, supporting earlier findings that some loci are instead under divergent selection for adaptation over the high-low shore environmental axis (Morales *et al*., 2019). Interestingly, we also found a clear association between arrangement frequency and position on the shore within the Spanish Crab ecotype in our study, further supporting the role of this inversion in adaptation to high-low shore selection gradients. Hence, in Spain, where the two selection gradients coincide, the total divergent selection between ecotypes and the proportion of the genome under divergent selection are likely to be inflated compared to in Sweden. This may result in a stronger barrier to gene flow in the Spanish system.

## Conclusions

Our findings underscore the value of high resolution and multi-dimensional data in natural replicated experiments within the same species in characterizing the intricate nature of speciation. Thanks to a remarkable study system provided by a marine snail in which two parallel ecotypes have advanced to different points on the speciation continuum, we show that reproductive isolation can arise despite a history of gene flow but its progress towards completion revolves around complex ecological and/or evolutionary dynamics. Future studies will clarify the interplay and the relative contributions of extrinsic environmental factors (e.g., spatial configuration, divergent selection) and intrinsic components (e.g., time of divergence, chromosomal rearrangements, genetic incompatibilities, opportunities for reinforcement) in the different levels of reproductive isolation observed between the Swedish and Spanish systems. Differences and similarities between the *L. saxatilis’* Swedish and Spanish pairs together offer an outstanding opportunity to compare barriers to gene flow of diverse strengths without being confounded with differences that might have accumulated in comparisons among distinct taxa, and so contributing to unravelling the mystery of speciation.

## Acknowledgements

We thank Hernan Morales, Jenny Larsson, Pragya Chaube, Isobel Eyres, Samuel Perini, Richard Turney, Graciela Sotelo, and Alison Butlin for help with fieldwork and snail processing. We are grateful to Matteo Fumagalli, Andrea Benazzo, and Sean Stankowski for suggestions on read processing. We also thank Jenny Larsson for guidance on using the ShellShaper program and Luciano Bosso for recommendations on the graphical abstract. The analysis of genomic data was conducted on the University of Sheffield high performance computing cluster, ShARC. Funding was provided by European Research Council grant 693030-BARRIERS to RKB; the Swedish Science Research Council (grant number 2021-04191) to KJ; the Portuguese Foundation for Science and Technology (FCT: 2020.00275.CEECIND and PTDC/BIA-EVL/1614/2021) to RF; Xunta de Galicia (Centro singular de investigación de Galicia accreditation 2019-2022), the European Union (European Regional Development Fund - ERDF), Xunta de Galicia (ED431C 2020-05) and Ministerio de Ciencia, Innovación y Universidades (PID2021-124930NB-I00) to JG and ERA.

## Data Accessibility

Raw sequence reads are deposited in NCBI SRA (BioProject accession ID PRJNA781449 that will be available upon publication). Phenotypic data (maturity, sex, wet weight, shell thickness, shell ridging, shell striping, shell length, shell growth parameters, and boldness scores) will be archived in Zenodo and scripts on Github upon publication.

## Benefit-Sharing

Benefits Generated: A research collaboration was developed with scientists from the countries where fieldwork was conducted, all collaborators are included as co-authors, data and results of research have been shared with the provider communities and the broader scientific community through appropriate biological databases as described above. The research addresses a priority concern regarding how biodiversity is generated and maintained. More broadly, our group is committed to international scientific partnerships, as well as institutional capacity building.

## Author Contributions

RKB, KJ, AMW, FR, and ADJ designed the study. RKB, KJ, AMW, ZZ, RF, JG, ERA collected data. RKB, FR, ADJ, and RF analysed data. FR, ADJ, RKB and KJ drafted the manuscript. All authors revised and agreed to the manuscript.

---

**Cover image.**
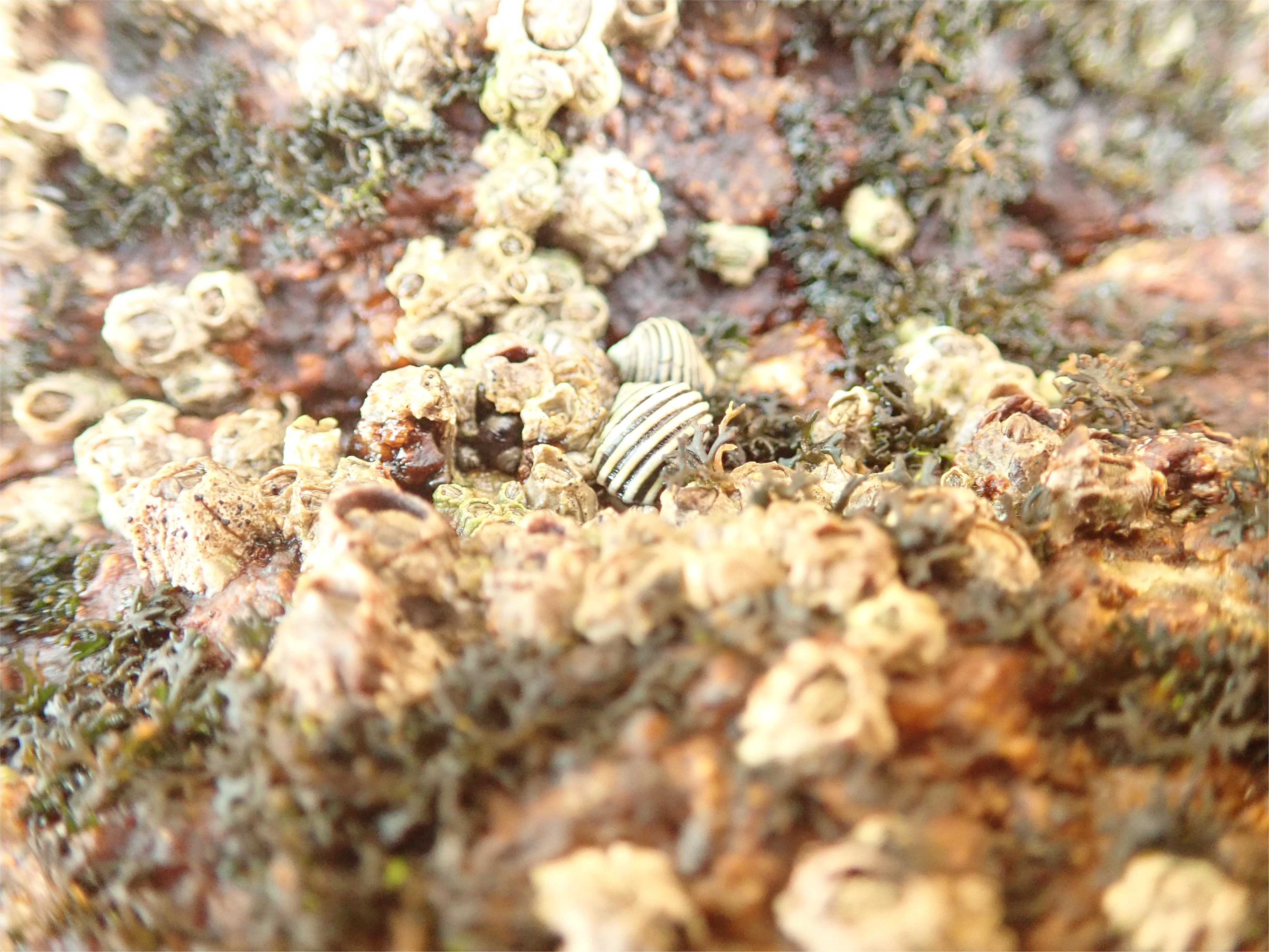
The marine snail *Littorina saxatilis* in the Spanish shore. Photo credits: Aurélien De Jode.

