## Supplementary Information for "Phenotypic divergence and genomic architecture between parallel ecotypes at two different points on the speciation continuum in a marine snail"

### Index

Differentiation between inverted and collinear regions

Inversion results

Tables:

Table S1. Kendall test of concordance

Table S2. Admixture ancestry fraction (separate xls file)

Table S3. Global $F_{ST}$ and $F_{ST}$ by linkage group

Table S4. List of contigs showing higher differentiation in Sweden and Spain

Table S5. Summary of the linear models for phenotypes

Table S6. Linear models phenotypes (separate docx file)

Figures:

Figure S1. Spain - seasonal patterns

Figure S2. Spain and Sweden - number of genetic clusters

Figure S3. Spain - genetic PCA

Figure S4. Spain - genetic PC1 within-group correlation

Figure S5. Spain - habitats association

Figure S6. Joint dataset - genetic patterns

Figure S7. Joint dataset - number of genetic clusters

Figure S8. Sweden - Manhattan plot by LG

Figure S9. Spain - Manhattan plot by LG

Figure S10. Sweden and Spain - average FST per contig

Figure S11. Sweden - genetic PCA by LG

Figure S12. Spain - genetic PCA by LG

Figure S13. PCA per inversion positions in Spain and Sweden

Figure S14. Sweden – Distribution of Fst differences

Figure S15. Spain - Distribution of Fst differences

Figure S16. Arrangement counts per transect position in Spain.

Figure S17. Genomic PC2 against shore position in Spain

Figure S18. Sweden and Spain - genetic patterns without inversions

Figure S19. Sweden and Spain - number of genetic clusters without inversions

Figure S20. Spain - PC1 within-group correlation without inversions

Figure S21. Spain - individual phenotypes

Figure S22. Sweden and Spain - relative phenotypic divergence

Figure S23. Phenotype PCs along the shore.

Figure S24. Joint dataset - phenotypic PCA

Figure S25. Number of abortive embryos between Spanish ecotypes crossings

### **Differentiation between inverted and collinear regions.**

Permutations tests for $F_{ST}$ differences between inverted and collinear regions were conducted in 10 out of the 17 LGs (Fig. S14-S15). $F_{ST}$ in inverted regions were significantly higher than in collinear regions in LG4, LG6, LG7, LG14 and LG17 in both countries, and in LG2 and LG9 in Spain only. In LG6, LG7 and LG14 the differences in $F_{ST}$ between inverted and collinear regions were similar between Spain and Sweden while for LG4 and LG17 the $F_{ST}$ differences were much higher in Spain. For LG1, LG2 and LG9 in Sweden, and LG11 in both countries, $F_{ST}$ were relatively similar along the LGs. The only case where $F_{ST}$ in inverted regions was lower than in collinear regions, was in LG1 in Spain where a large portion around the centre of the LG showed high levels of differentiation (Fig. S9). $F_{ST}$ differences between inverted and collinear regions were substantial though not statistically significant in LG5, probably because the inverted regions covered the majority of the LG.

### **Inversion results**

Nine of the shared inversions (LGC1.1, LGC1.2, LGC2.1, LGC4.1, LGC7.1, LGC9.1, LGC10.2, LGC11.1 and LGC17.1) presented patterns consistent with a single inversion while LGC12.2 and LGC12.3 showed a complex inversion signal in both countries (Fig. 3, Table 1, Fig. S13). The other eight inversions showed contrasting patterns between Spain and Sweden. LGC6.1/2, LGC7.2 and LGC12.4 exhibited clear inversion signals but different inversion types (e.g. simple or complex) in each country. LGC5.1, LGC9.2, LGC14.1.2 and LGC14.3 were identified in Spain but not in this Swedish sample while LGC12.1 was detected in Sweden but not in Spain.

In Spain, a clear association between arrangements and genetic clusters was detected in all inversions, while in Sweden this association was not significant for LGC2.1, LGC4.1, LGC9.1, LGC10.2, LGC11.1 (Table 1). For LGC1.1, LGC1.2, LGC2.1, LGC4.1, LGC7.1, LGC9.1, LGC10.2, LGC11.1, LGC17.1, single inversion pattern were identified in both countries on the projected PCA plot (Fig. S13), allowing us to compare the frequencies of the same arrangements between ecotypes and countries. In LGC1.1, LGC1.2, LGC7.1 and LGC17.1, the arrangement that had a higher frequency in Wave compared to Crab in Spain, was also more frequent in the Wave end of the transect compared to the Crab end in Sweden. The frequency difference was larger in Spain than in Sweden for LGC1.2, and LGC17.1. Conversely, for LGC9.1, LGC10.2 and LGC11.1, the arrangement with a higher frequency in Wave compared to Crab in Spain, had a similar or a lower frequency in the Wave end of the transect compared to the Crab end in Sweden. For those three inversions, arrangement frequency differences between ecotypes were not significant in Sweden.

In Spain, the most common arrangement was fixed in Wave in LGC1.1, LGC1.2, LGC2.1, LGC7.1, LGC9.1, LGC10.2, LGC12.2, LGC12.3 and LGC12.4 while Crab did not show any fixed arrangement in any inversion (Table 1, Fig. S13). Within the Crab cluster, a significant positive correlation between the frequency of the arrangement that is most abundant in Wave and shore position (i.e. Crab individuals in the low shore and in the region of overlap with the Wave distribution were more likely to carry the arrangement typical of the Wave ecotype) was detected in LGC1.1, LGC5.1, LGC6.1/2, LGC7.1, LGC7.2, LGC9.1, LGC9.2, LGC10.2, LGC11.1, LGC12.2, LGC12.3 and LGC17.1 (Table 1, Fig S13-S16). Interestingly, in Spain, arrangement clusters in the inversions LGC9.1, LGC10.2, LGC11.1 and LGC12.4 were split in two sub-clusters along PC2 comprising either individuals belonging to the Wave or Crab genetic group (Fig. S13). This indicates genetic divergence of allelic content between, for example, arrangement A1 for LGC9.1 in Wave compared to the same arrangement in Crab. Conversely, arrangement clusters in the inversions LGC1.1, LGC1.2, LGC2.1, LGC4.1, LGC5.1, LGC7.1, LGC7.2, LGC9.2, LGC17.1 showed a complete to partial overlap between Crab and Wave genetic group (Fig. S13). This indicates genetic similarity between the same arrangement found in each ecotype. Moreover, for LGC1.2, arrangement found in Crab individuals were more wave-like in the lower part of the shore (Fig. S17), suggesting that gene flow might occur in this particular arrangement between the Crab and Wave genetic groups.

The previously known inversions LGC14.1/2 and LGC14.3 were not identified in this Swedish sample while they showed a complex pattern in Spain. Here, the LGC14.1/2 PCA by inversion showed a split into two clusters in Wave (named W1 and W2). W1 included only female individuals while W2 encompassed 37 males and two females (Fig. 3, S15). Similarly, the Wave group in LGC14.3 was split into two clusters with the same sex bias as in 14.1/2. In addition, the Crab group was subdivided in three clusters (named C1, C2 and C3) with 26 males, 33 females plus 23 males, and 17 females plus four males in the C1, C2 and C3 clusters, respectively (Fig. 3, S15). In Sweden, LGC12,2 and LGC12.3 also presented some association with sex, a pattern that was previously described in Hearn et al. (2022).

Table S1. Kendall's test of concordance among the genetic analyses of the three (unthinned) and nine (thinned) random subsets: first two principal components, Hybrid Index, Bayesian information criterion of the discriminant analysis of principal components (BIC), Admixture cluster assignment to either the crab or wave ecotype (Q reported only for K=2) and cross validation error (CV), and $F_{ST}$ values. One genetic cluster was identified in Sweden, but two populations (ends of the transect) were forced here to facilitate comparisons with Spain.

| Dataset | Analysis | χ2 | df | p-value | W |
|---|---|---|---|---|---|
| Spain with inversions | PC1 | 1661 | 187 | < 2.2e-16 | 0.9869496 |
| | PC2 | 1282.5 | 187 | < 2.2e-16 | 0.7620255 |
| | Hybrid Index | 1640.1 | 187 | < 2.2e-16 | 0.9745188 |
| | BIC | 126 | 14 | < 2.2e-16 | 1 |
| | Q - crab | 1552.6 | 187 | < 2.2e-16 | 0.9225197 |
| | Q - wave | 1552.6 | 187 | < 2.2e-16 | 0.9225197 |
| | CV | 36 | 4 | 2.894e-07 | 1 |
| | FST | 946679 | 339615 | < 2.2e-16 | 0.9291691 |
| Sweden with inversions | PC1 | 844.34 | 96 | < 2.2e-16 | 0.9772408 |
| | PC2 | 407.97 | 96 | < 2.2e-16 | 0.9064993 |
| | Hybrid Index | 841.49 | 96 | < 2.2e-16 | 0.9739513 |
| | BIC | 126 | 14 | < 2.2e-16 | 1 |
| | Q - crab | 847.65 | 96 | < 2.2e-16 | 0.9810765 |
| | Q - wave | 847.65 | 96 | < 2.2e-16 | 0.9810765 |
| | CV | 36 | 4 | 2.894e-07 | 1 |
| | FST | 650096 | 294784 | < 2.2e-16 | 0.7351094 |
| Spain without inversions | PC1 | 1654.3 | 187 | < 2.2e-16 | 0.9829257 |
| | PC2 | 621.01 | 187 | < 2.2e-16 | 0.3689915 |
| | Hybrid Index | 1634.9 | 187 | < 2.2e-16 | 0.9714118 |
| | BIC | 126 | 14 | < 2.2e-16 | 1 |

| | | | | | |
|---|---|---|---|---|---|
| | Q - crab | 1581.9 | 187 | < 2.2e-16 | 0.9399105 |
| | Q - wave | 1581.9 | 187 | < 2.2e-16 | 0.9399105 |
| | CV | 36 | 4 | 2.894e-07 | 1 |
| | FST | 670065 | 243857 | < 2.2e-16 | 0.9159265 |
| Sweden without inversions | PC1 | 834.78 | 96 | < 2.2e-16 | 0.966176 |
| | PC2 | 479.02 | 96 | < 2.2e-16 | 0.5544185 |
| | Hybrid Index | 814.81 | 96 | < 2.2e-16 | 0.9430668 |
| | BIC | 126 | 14 | < 2.2e-16 | 1 |
| | Q - crab | 838.71 | 96 | < 2.2e-16 | 0.9707268 |
| | Q - wave | 838.71 | 96 | < 2.2e-16 | 0.9707268 |
| | CV | 36 | 4 | 2.894e-07 | 1 |
| | FST | 442444 | 208655 | < 2.2e-16 | 0.7068194 |
| Joint | PC1 | 2492.3 | 283 | < 2.2e-16 | 0.9785218 |
| | PC2 | 2496.4 | 283 | < 2.2e-16 | 0.98013 |
| | BIC | 126 | 14 | < 2.2e-16 | 1 |

Table S3. Average genome-wide $F_{ST}$ and $F_{ST}$ by linkage group (LG) values in the Spanish (two genetic clusters) and Swedish (ends of the transect) unthinned subsets for whole LG, LG without inversions regions and in inversions. Grey lines indicate LG without inversions. NA values in the “without inversions” columns indicates that no $F_{ST}$ value was computed because the whole LG contained inversion regions. NA values in the “Inversions” columns indicates that no $F_{ST}$ value was computed because there was no inversion in this LG. * “Inversions” columns indicates that $F_{ST}$ difference between inverted regions and none inverted regions was significant according to the permutation test. “NT” in the “Inversions” columns indicates no permutation test was conducted for this LG because the whole LG was affected by inverted regions.

| Linkage group | Spain | | | Sweden | | |
|---|---|---|---|---|---|---|
| | Whole LG | Without inversions | Inversions | Whole LG | Without inversions | Inversions |
| LG1 | 0.11 | 0.12 | 0.08 | 0.08 | 0.08 | 0.08 |
| LG2 | 0.13 | 0.12 | 0.33* | 0.09 | 0.08 | 0.09 |
| LG3 | 0.13 | 0.13 | NA | 0.08 | 0.08 | NA |
| LG4 | 0.09 | 0.07 | 0.22* | 0.06 | 0.06 | 0.07* |
| LG5 | 0.14 | 0.03 | 0.17 | 0.07 | 0.05 | 0.08 |
| LG6 | 0.13 | 0.08 | 0.25* | 0.12 | 0.08 | 0.24* |
| LG7 | 0.09 | 0.08 | 0.13* | 0.09 | 0.06 | 0.13* |
| LG8 | 0.05 | 0.05 | NA | 0.06 | 0.06 | NA |
| LG9 | 0.10 | 0.04 | 0.13* | 0.07 | 0.06 | 0.08 |
| LG10 | 0.08 | NA | 0.08 NT | 0.06 | NA | 0.06 NT |
| LG11 | 0.07 | 0.04 | 0.08 | 0.07 | 0.05 | 0.08 |
| LG12 | 0.19 | NA | 0.19 NT | 0.12 | NA | 0.12 NT |
| LG13 | 0.06 | 0.06 | NA | 0.06 | 0.06 | NA |
| LG14 | 0.19 | 0.05 | 0.20* | 0.17 | 0.03 | 0.18* |
| LG15 | 0.06 | 0.06 | NA | 0.07 | 0.07 | NA |
| LG16 | 0.05 | 0.05 | NA | 0.05 | 0.05 | NA |
| LG17 | 0.11 | 0.08 | 0.31* | 0.08 | 0.07 | 0.13* |
| Genome-wide | 0.11 | 0.09 | 0.16 | 0.08 | 0.07 | 0.11 |

Table S4. List of contigs showing higher differentiation (average $F_{ST}$ higher than 0.3) between the two genetic clusters (Spain) or ends of the transect (Sweden). Inversions identified in previous studies are highlighted in grey.

| Contig | Mean $F_{ST}$ Sweden | Mean $F_{ST}$ Spain | Linkage group | AvgMP |
|---|---|---|---|---|
| Contig3429 | 0.39 | 0.50 | LG1 | 24.59 |
| Contig40025 | 0.31 | 0.61 | LG1 | 45.03 |
| Contig39800 | 0.62 | 0.43 | LG1 | 45.92 |
| Contig55764 | 0.33 | 0.43 | LG2 | 42.16 |
| Contig47588 | 0.31 | 0.44 | LG2 | 52.04 |
| Contig44328 | 0.31 | 0.37 | LG2 | 53.06 |
| Contig6656 | 0.36 | 0.42 | LG3 | 30.44 |
| Contig3407 | 0.41 | 0.35 | LG3 | 30.51 |
| Contig54177 | 0.58 | 0.52 | LG3 | 30.51 |
| Contig3544 | 0.32 | 0.36 | LG5 | 22.53 |
| Contig6361 | 0.36 | 0.32 | LG6 | 0.00 |
| Contig7009 | 0.48 | 0.39 | LG6 | 0.00 |
| Contig44587 | 0.51 | 0.36 | LG6 | 0.00 |
| Contig4622 | 0.30 | 0.30 | LG6 | 1.37 |
| Contig40766 | 0.45 | 0.39 | LG6 | 3.20 |
| Contig6653 | 0.42 | 0.40 | LG6 | 3.90 |
| Contig46033 | 0.42 | 0.31 | LG6 | 5.11 |
| Contig38260 | 0.45 | 0.42 | LG6 | 5.39 |
| Contig7695 | 0.57 | 0.34 | LG6 | 6.68 |
| Contig48459 | 0.41 | 0.38 | LG6 | 6.68 |
| Contig56105 | 0.43 | 0.52 | LG6 | 8.15 |
| Contig4701 | 0.32 | 0.51 | LG6 | 11.19 |
| Contig38821 | 0.37 | 0.33 | LG6 | 17.44 |
| Contig43679 | 0.31 | 0.42 | LG6 | 18.60 |
| Contig49086 | 0.32 | 0.36 | LG6 | 19.76 |
| Contig5843 | 0.40 | 0.36 | LG6 | 19.76 |
| Contig2899 | 0.45 | 0.39 | LG6 | 19.76 |
| Contig985 | 0.45 | 0.47 | LG6 | 23.63 |
| Contig3228 | 0.47 | 0.47 | LG6 | 23.63 |
| Contig354 | 0.39 | 0.52 | LG6 | 23.63 |
| Contig376 | 0.37 | 0.35 | LG6 | 23.63 |
| Contig39724 | 0.39 | 0.43 | LG6 | 23.63 |
| Contig43191 | 0.66 | 0.46 | LG6 | 23.63 |
| Contig45103 | 0.33 | 0.43 | LG6 | 23.63 |
| Contig46665 | 0.34 | 0.33 | LG6 | 23.63 |
| Contig48389 | 0.34 | 0.32 | LG6 | 23.63 |
| Contig3509 | 0.39 | 0.35 | LG8 | 33.47 |
| Contig8254 | 0.50 | 0.42 | LG9 | 28.52 |
| Contig41383 | 0.30 | 0.32 | LG12 | 44.83 |
| Contig52467 | 0.32 | 0.42 | LG12 | 49.73 |
| Contig9802 | 0.32 | 0.38 | LG12 | 53.37 |
| Contig47257 | 0.32 | 0.31 | LG14 | 1.55 |
| Contig47125 | 0.30 | 0.30 | LG14 | 6.78 |
| Contig46983 | 0.32 | 0.31 | LG14 | 8.52 |
| Contig46460 | 0.30 | 0.42 | LG14 | 10.24 |
| Contig46795 | 0.35 | 0.33 | LG14 | 10.24 |
| Contig479 | 0.31 | 0.40 | LG14 | 10.24 |
| Contig48013 | 0.35 | 0.35 | LG14 | 10.24 |
| Contig50936 | 0.51 | 0.50 | LG14 | 10.24 |
| Contig51020 | 0.37 | 0.39 | LG14 | 10.24 |
| Contig991 | 0.49 | 0.32 | LG14 | 10.24 |
| Contig3777 | 0.47 | 0.39 | LG17 | 25.03 |
| Contig39278 | 0.30 | 0.51 | LG17 | 32.02 |

Table S5. Summary of the results for the linear model testing for the effect of genetic group and transect position on individual phenotypic traits. Cells shaded in grey represent phenotype trait that are independent of size.

| **Phenotypic trait** | **Significant term** |
|---|---|
| PC1 phenotype | Genetic group – Transect position |
| PC2 phenotype | Genetic group |
| Wet weight | Genetic group – Transect position – Interaction term |
| Width growth | Genetic group – Transect position |
| Height growth | Genetic group |
| r0 | Genetic group – Transect position |
| z0 | Genetic group – Transect position |
| a0 | Genetic group – Transect position |
| eccentricity | Genetic group – Transect position |
| apAngle | Genetic group |
| Shell length | Genetic group – Transect position |
| Apex1 | NONE |
| Apex2 | Genetic group – Transect position |
| Scale factor | Genetic group – Transect position |
| Thickness | Genetic group – Transect position |
| Extension factor | NONE |
| a0 scaled | Genetic group – Transect position |
| z0 scaled | Transect position |
| Log height growth | Genetic group |
| Log width growth | Genetic group – Transect position |
| Relative thickness | Genetic group |
| Convexity | NONE |

Figure S1. Seasonal patterns in Spain as shown by the first two genetic principal components of the thinned datasets (only one subset is shown here).

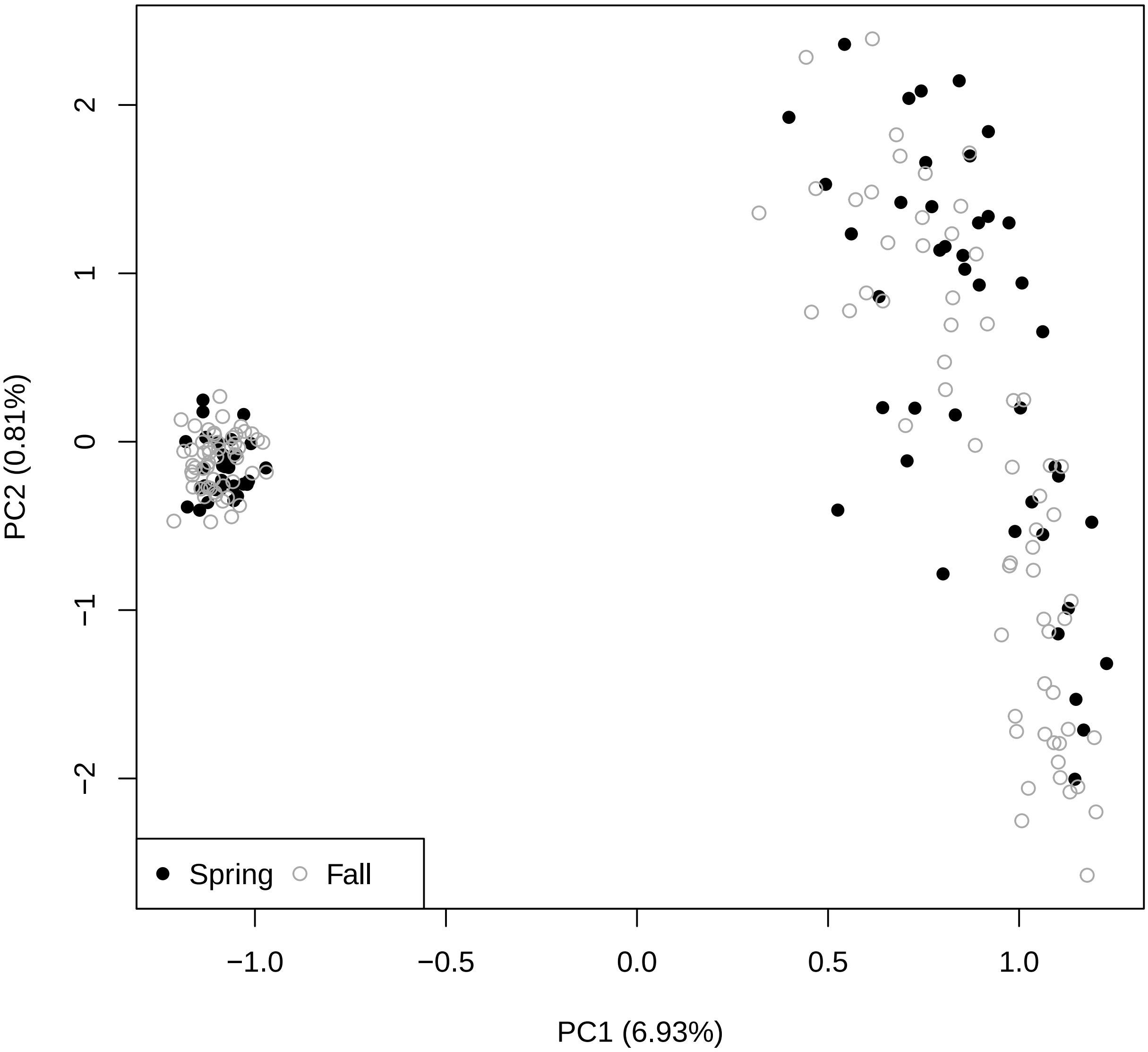

Figure S2. Number of genetic clusters (K) in Spain (a, c) and Sweden (b, d) as shown by the discriminant analyses of principal components (DAPCs, a, b) and Admixture cross-validation error (c, d).

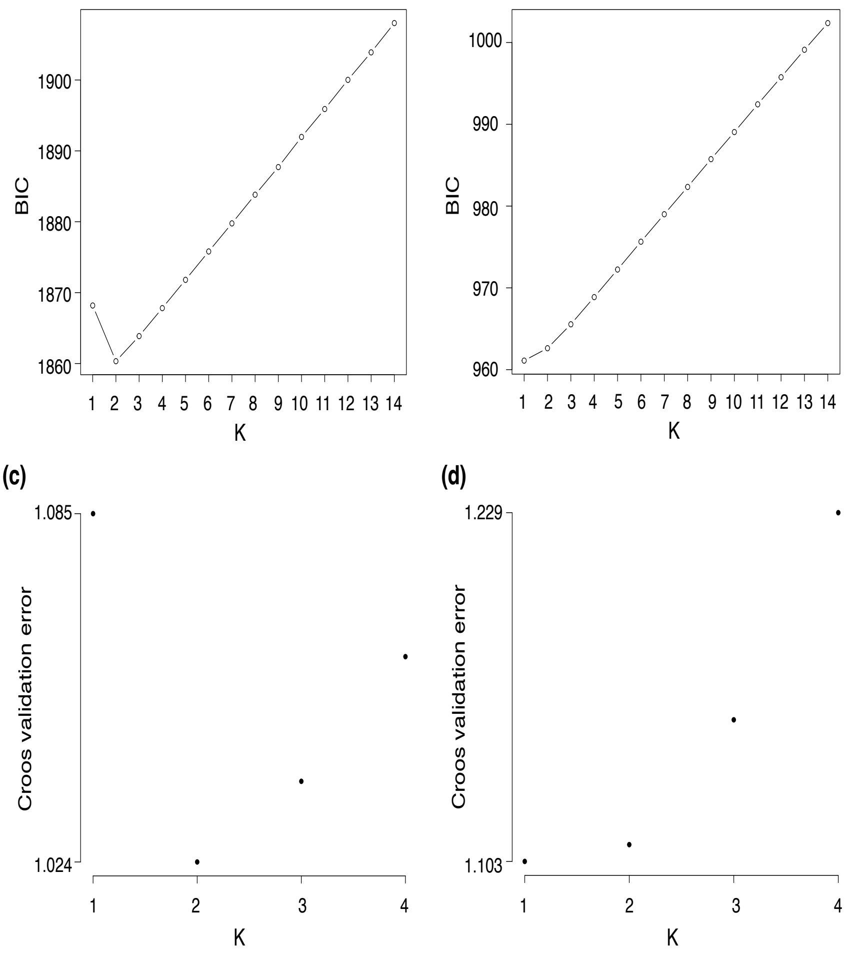

Figure S3. Pattern consistency of genetic differentiation in Spain as shown by the first two principal components in the nine thinned subsets. The Red genetic group and Blue genetic group are represented in blue and red, respectively. Abbreviations: s=random SNP subsampling, r= random read subsampling.

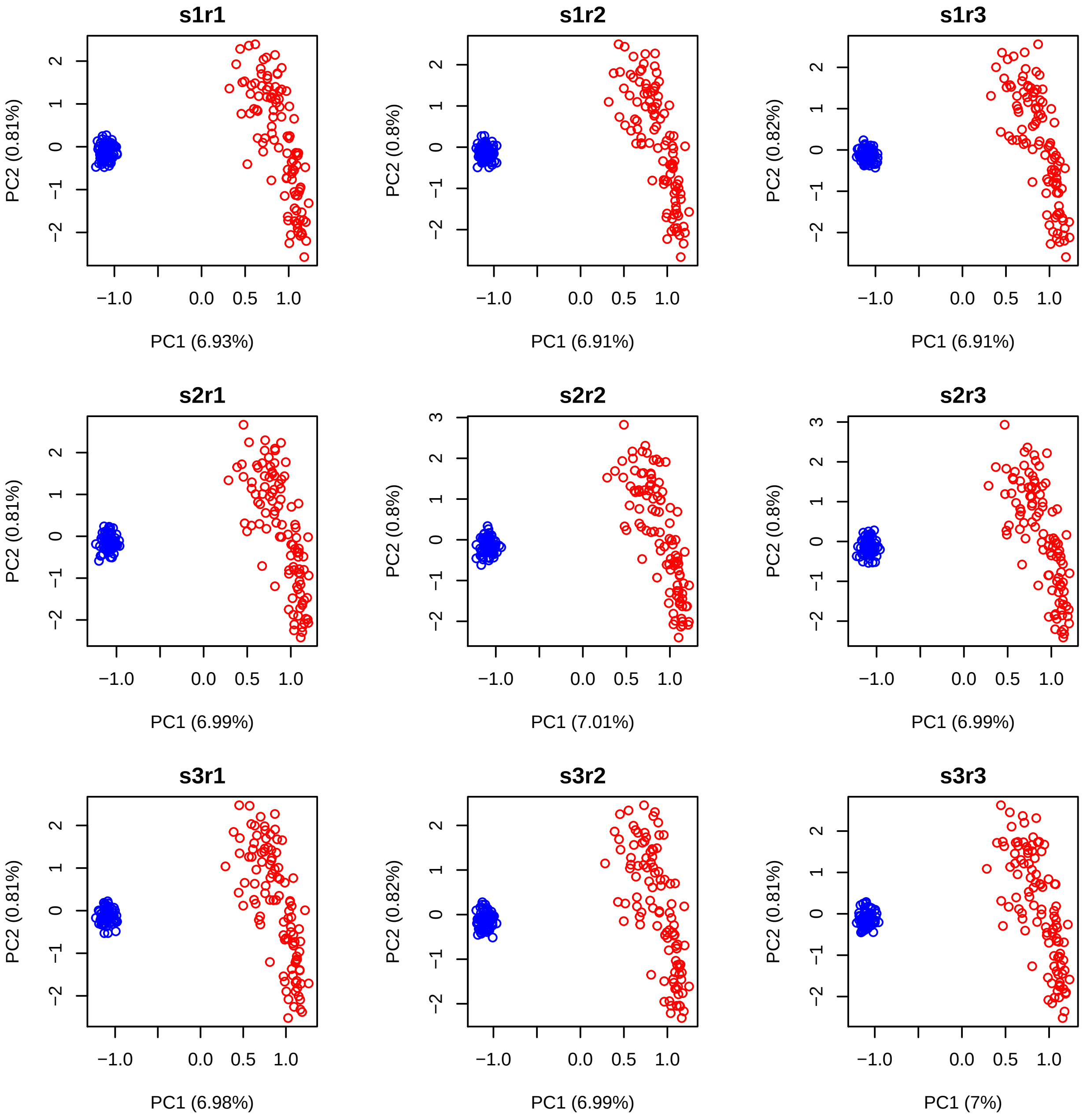

Figure S4. Correlation of the genetic within-cluster PC1 scores in the Red genetic group (a) and Blue genetic (b) group among the nine Spanish thinned subsets. Abbreviations: s=random SNP subsampling, r= random read subsampling.

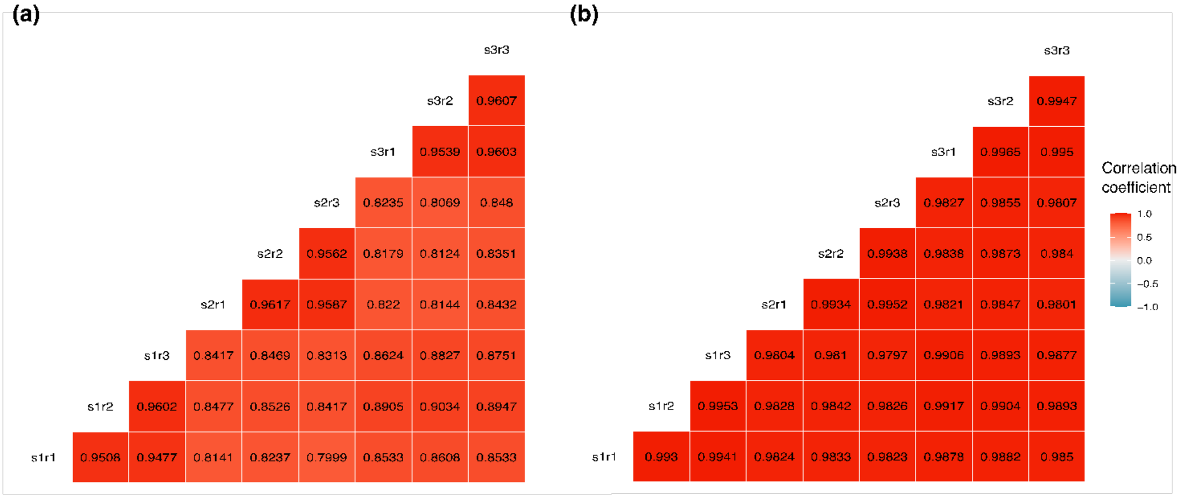

Figure S5. Association between the genetic clusters and habitat features along the Spanish transect: presence (filled dots) or absence (empty circles) of barnacles (Chthamalus, a), goose barnacles (Pollicipes, b) and mussels (Mytilus, c); and shore height (d). Transect positions closer to zero correspond to the high shore while higher values represent the low shore. The Crab and Wave group are represented in blue and red, respectively.

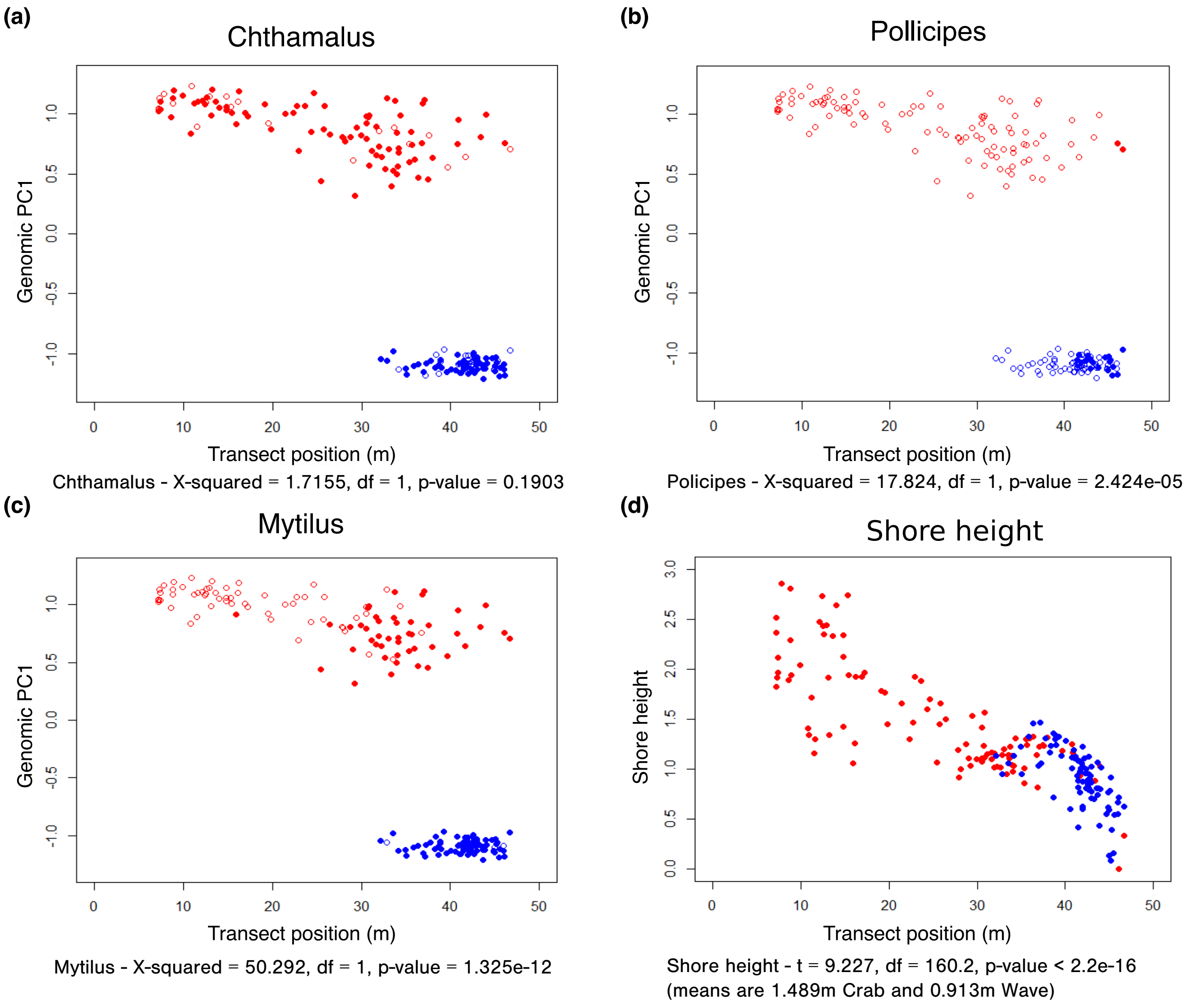

Figure S6. Genetic differentiation between countries as shown by the first two principal components in the joint dataset.

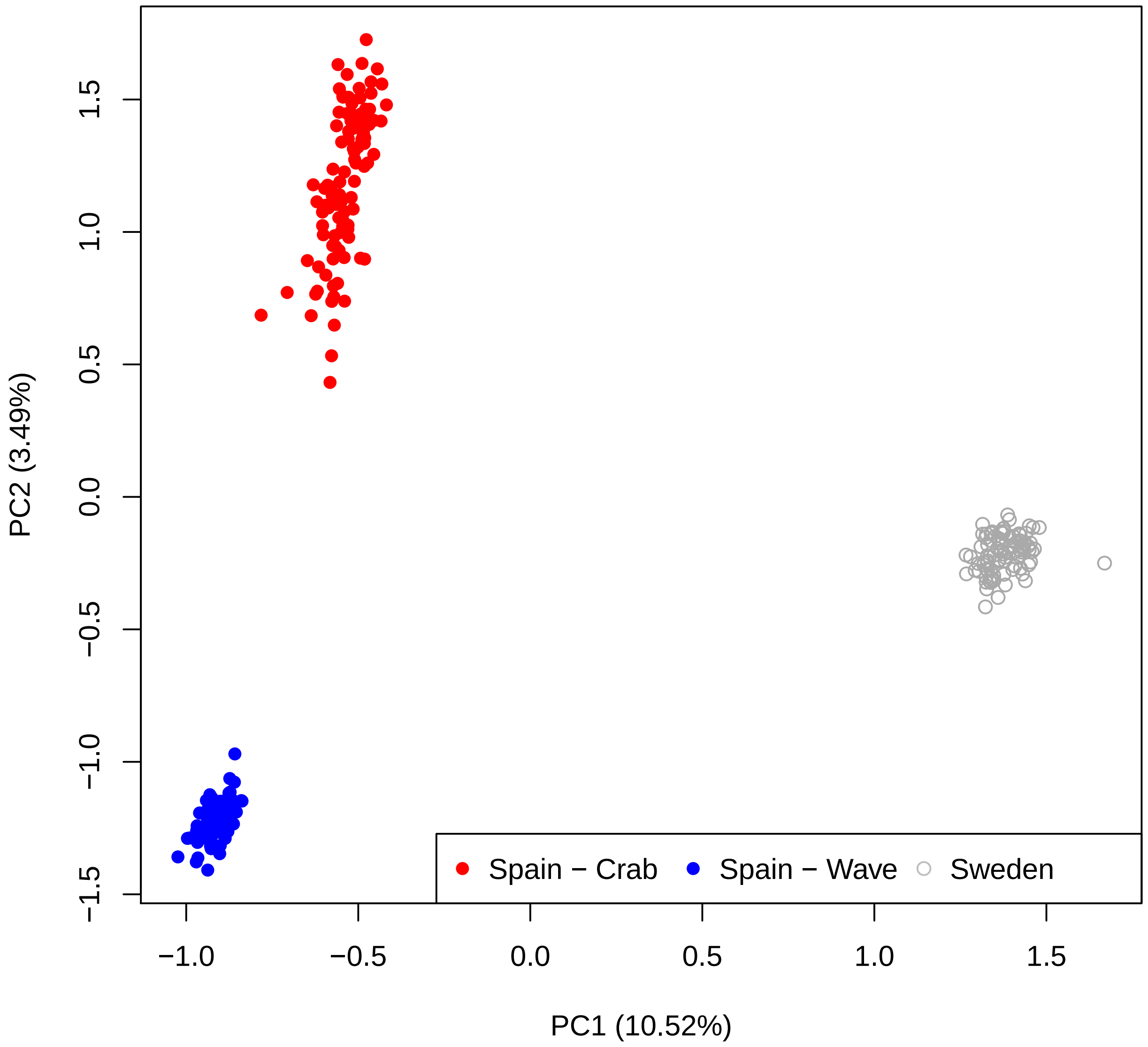

Figure S7. Number of genetic clusters in the joint dataset (Sweden and Spain) as shown by the discriminant analyses of principal components (DAPCs).

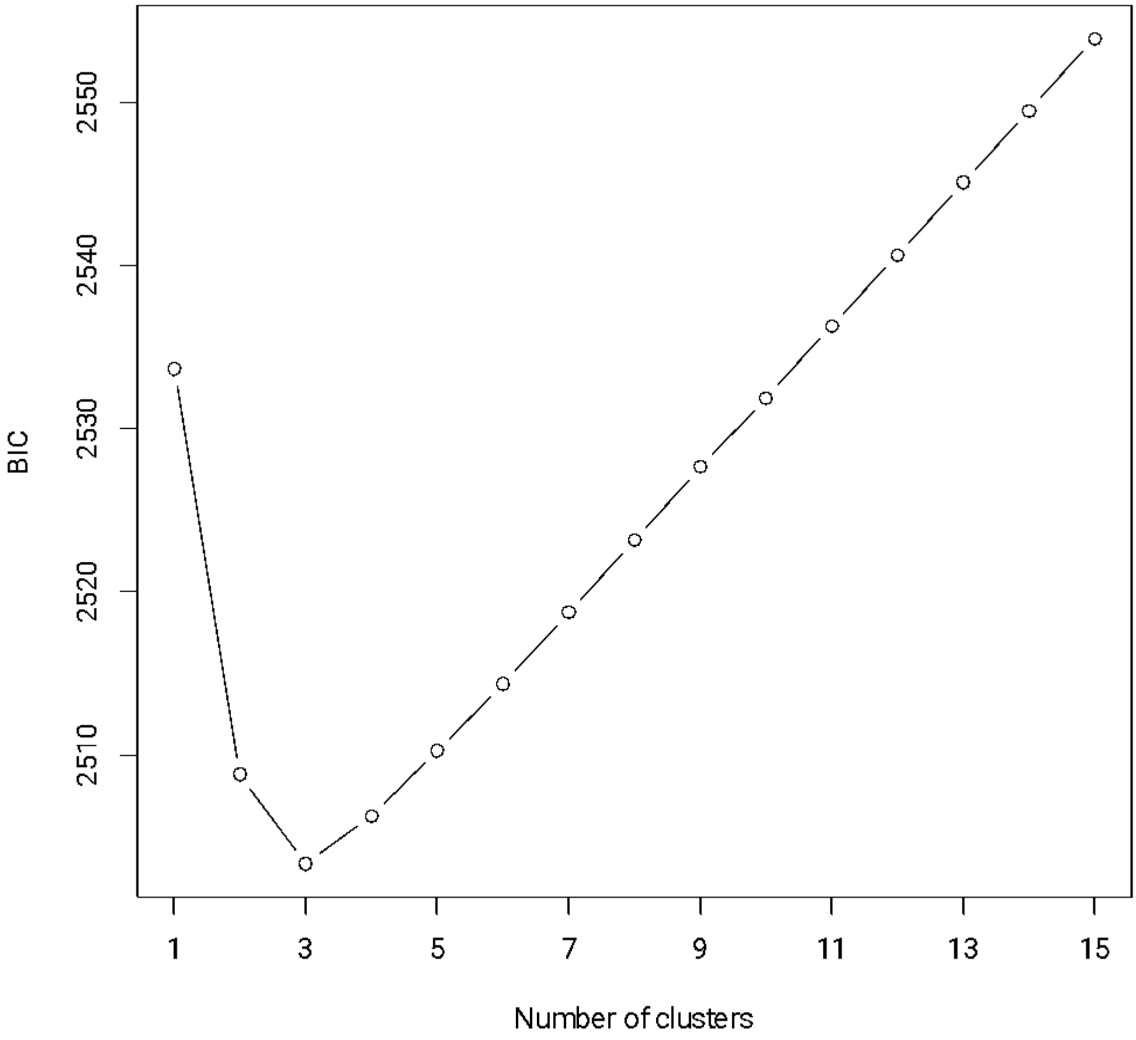

Figure S8. Differentiation between the Crab and Wave ecotypes along genome and chromosomal inversions as shown by Manhattan plots in Sweden. Grey rectangles indicate the position of known genomic inversions (Reeve et al. 2023). Horizontal black and red bars represent the average and 95th percentile of the Fst per map position.

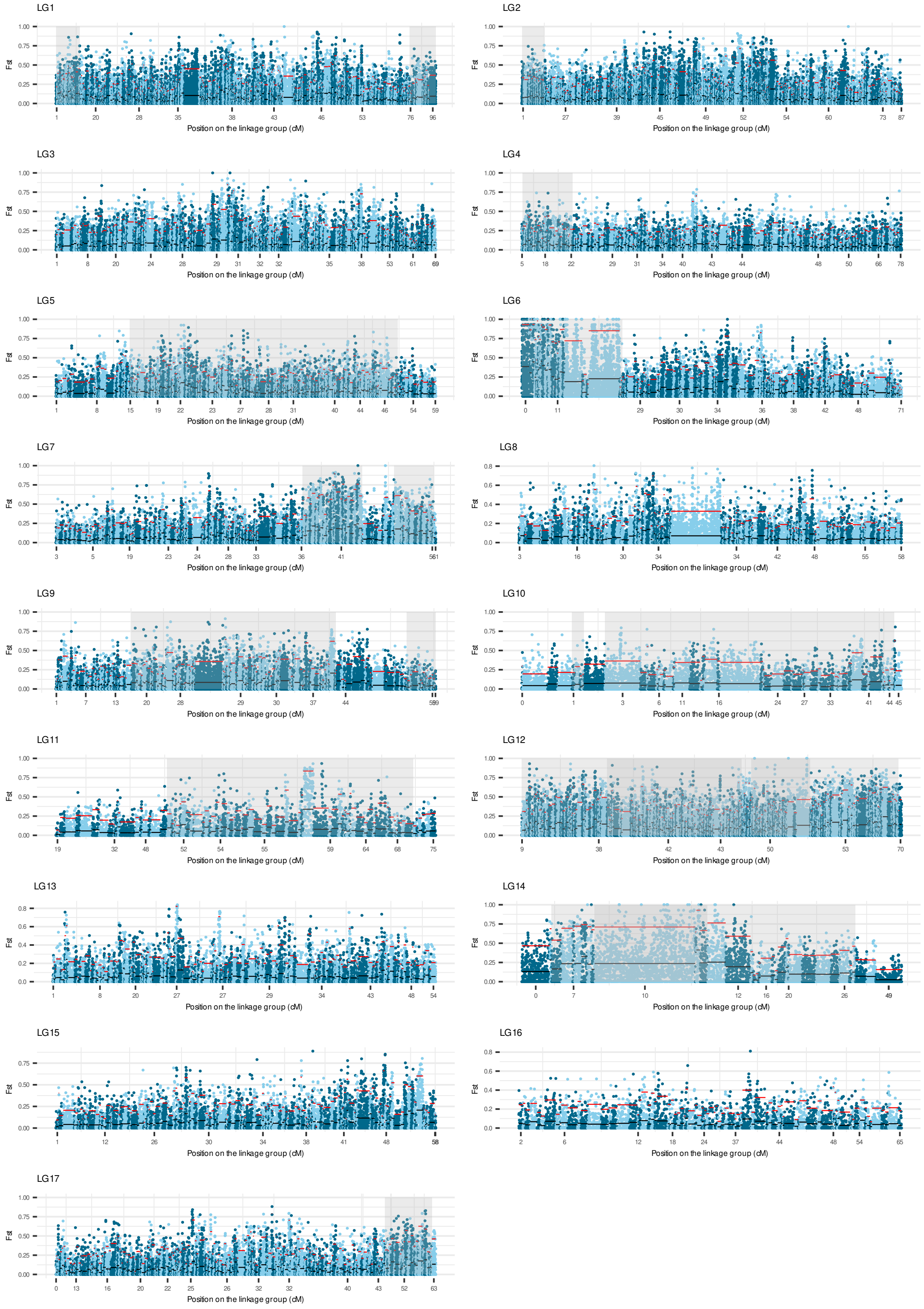

Figure S9. Differentiation between the Wave and Crab ecotypes along genome and chromosomal inversions as shown by Manhattan plots in Spain. Grey rectangles indicate the position of known genomic inversions (Reeve et al. 2023). Horizontal black and red bars represent the average and 95th percentile of the Fst per map position.

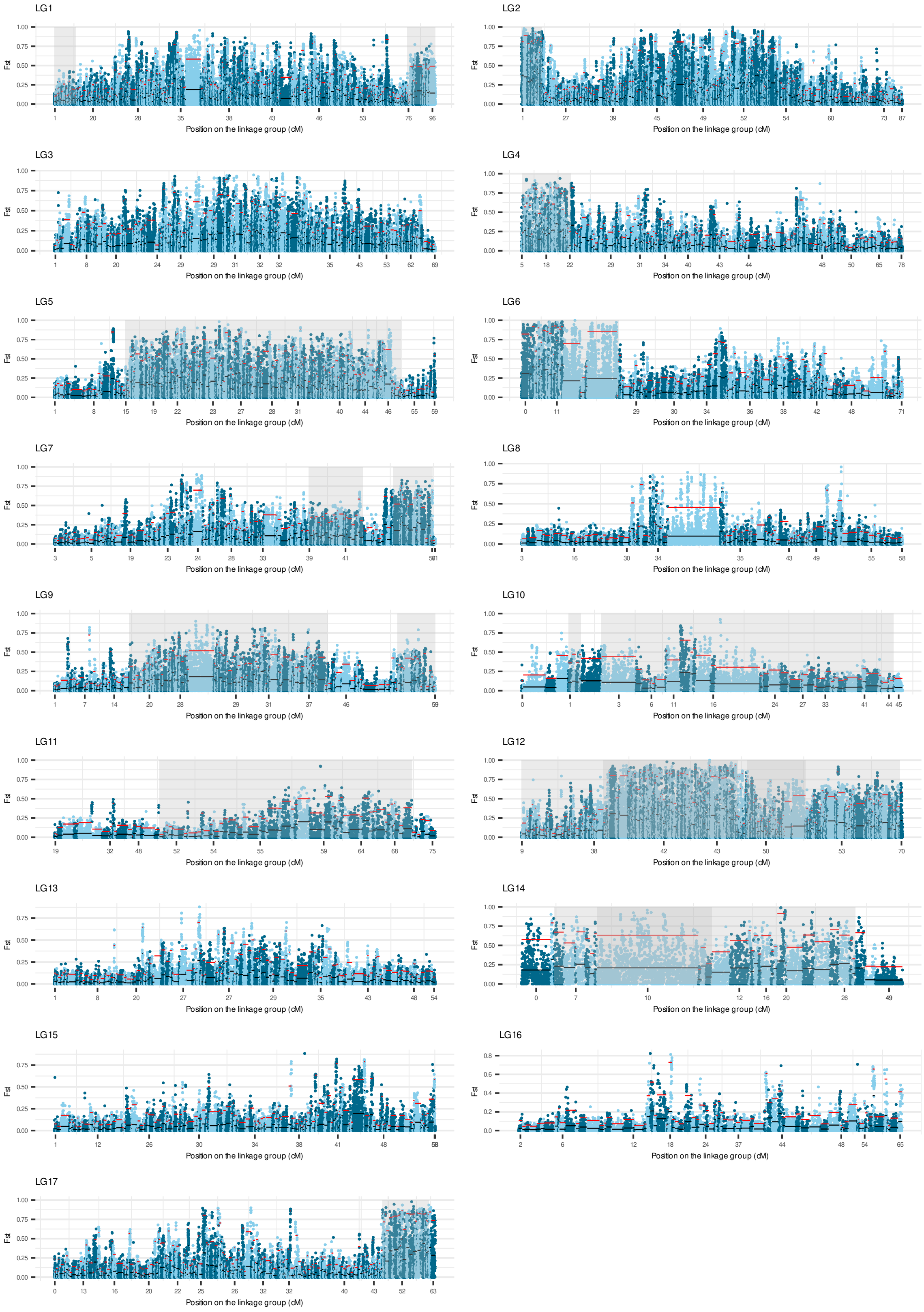

Figure S10. Correlation (Pearson coefficient) between divergence among the two genetic groups in Spain and the ends of the transect in Sweden as shown by average FST per contig. Coloured dots are in genomic regions containing inversions.

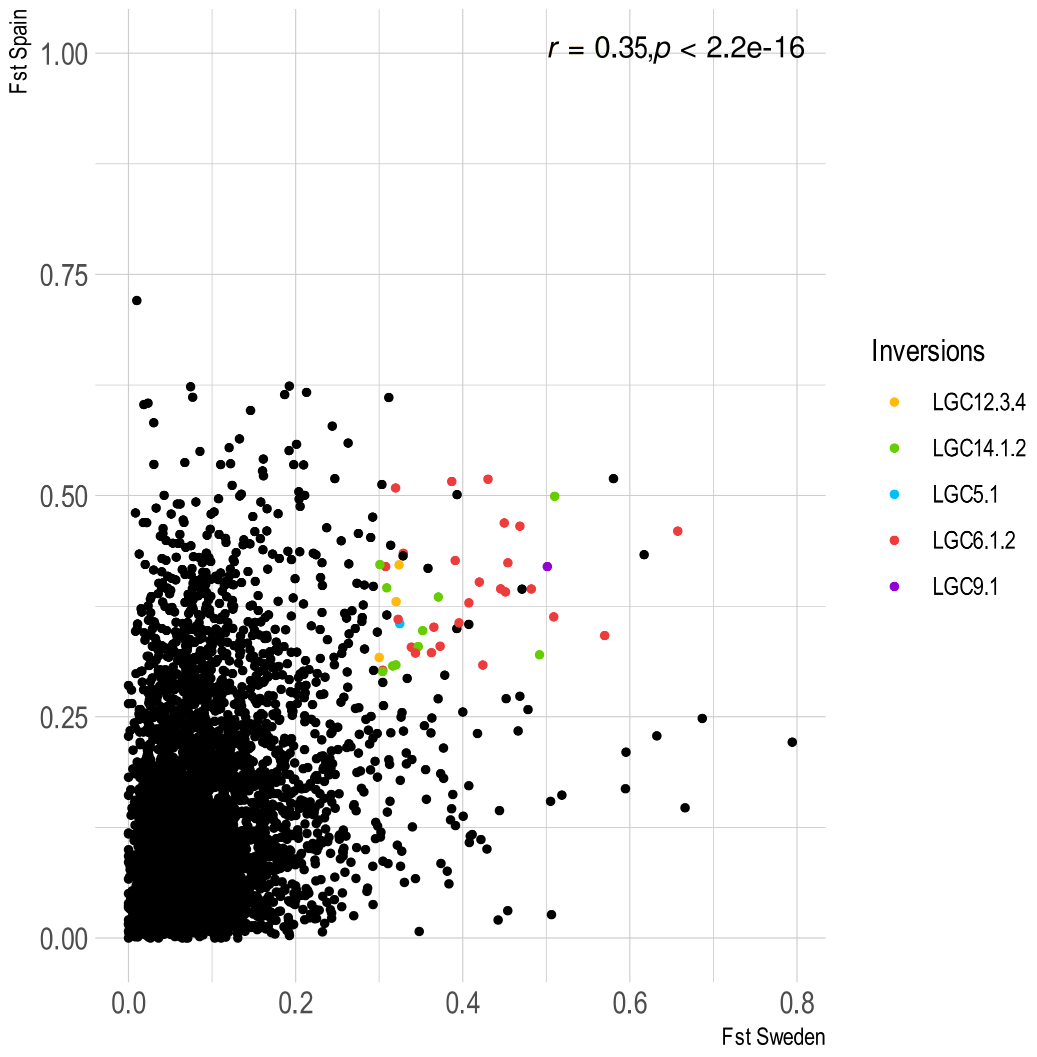

Figure S11. Divergence along genome and chromosomal inversions as shown by PCA by map position in Sweden. Individuals belonging to the Crab and Wave ecotype are represented by red and blue lines, respectively. Grey rectangles indicate the position of known genomic inversions (Reeve et al. 2023).

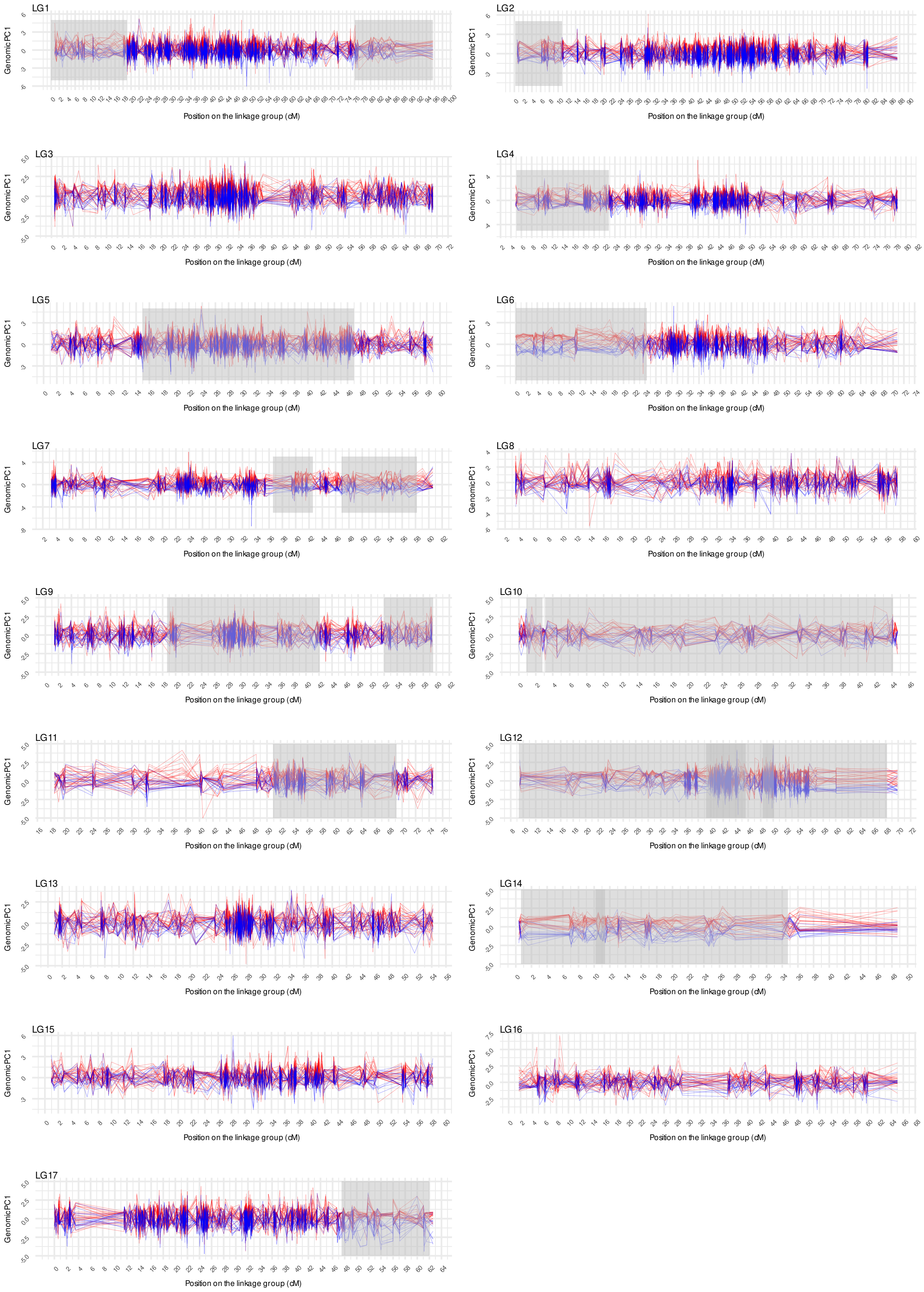

Figure S12. Divergence along genome and chromosomal inversions as shown by PCA by map position in Spain. Individuals belonging to the Crab and Wave ecotype are represented by red and blue lines, respectively. Grey rectangles indicate the position of known genomic inversions (Reeve et al. 2023).

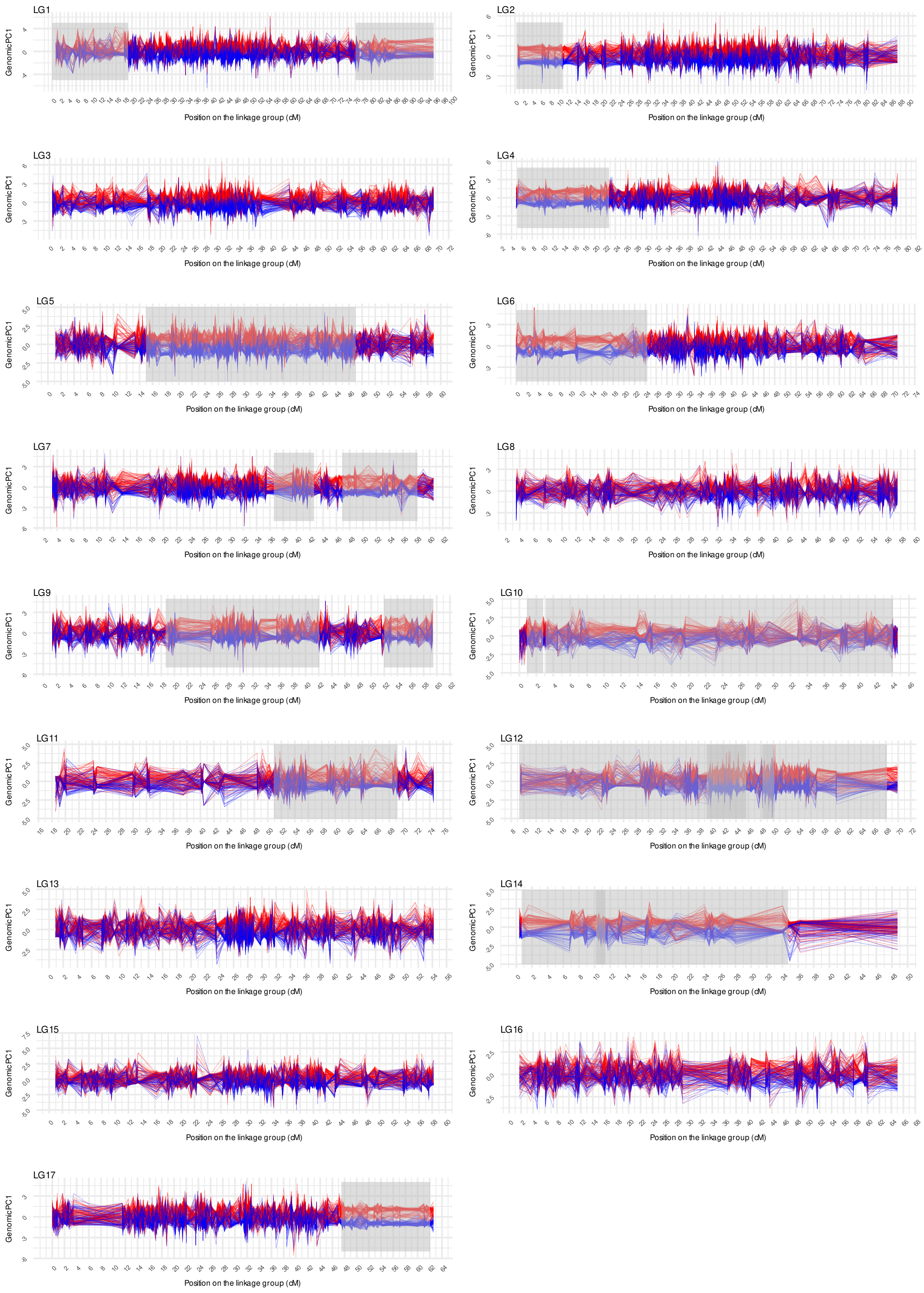

Figure S13. Genetic PCA Plots (PC1 vs PC2) by inverted regions of *L. saxatilis* (Reeve et al. 2023). Each row contains three PCA plots for each inversion. Column A1 to A20: PC1 vs PC2 in the Spanish transect. Column B1 to B20: PC1 vs PC2 in the Swedish transect. Column C1 to C20: PC1 vs PC2 in both countries (projected PCA approach). Spanish individuals are represented with filled dots and Swedish individuals with circles.

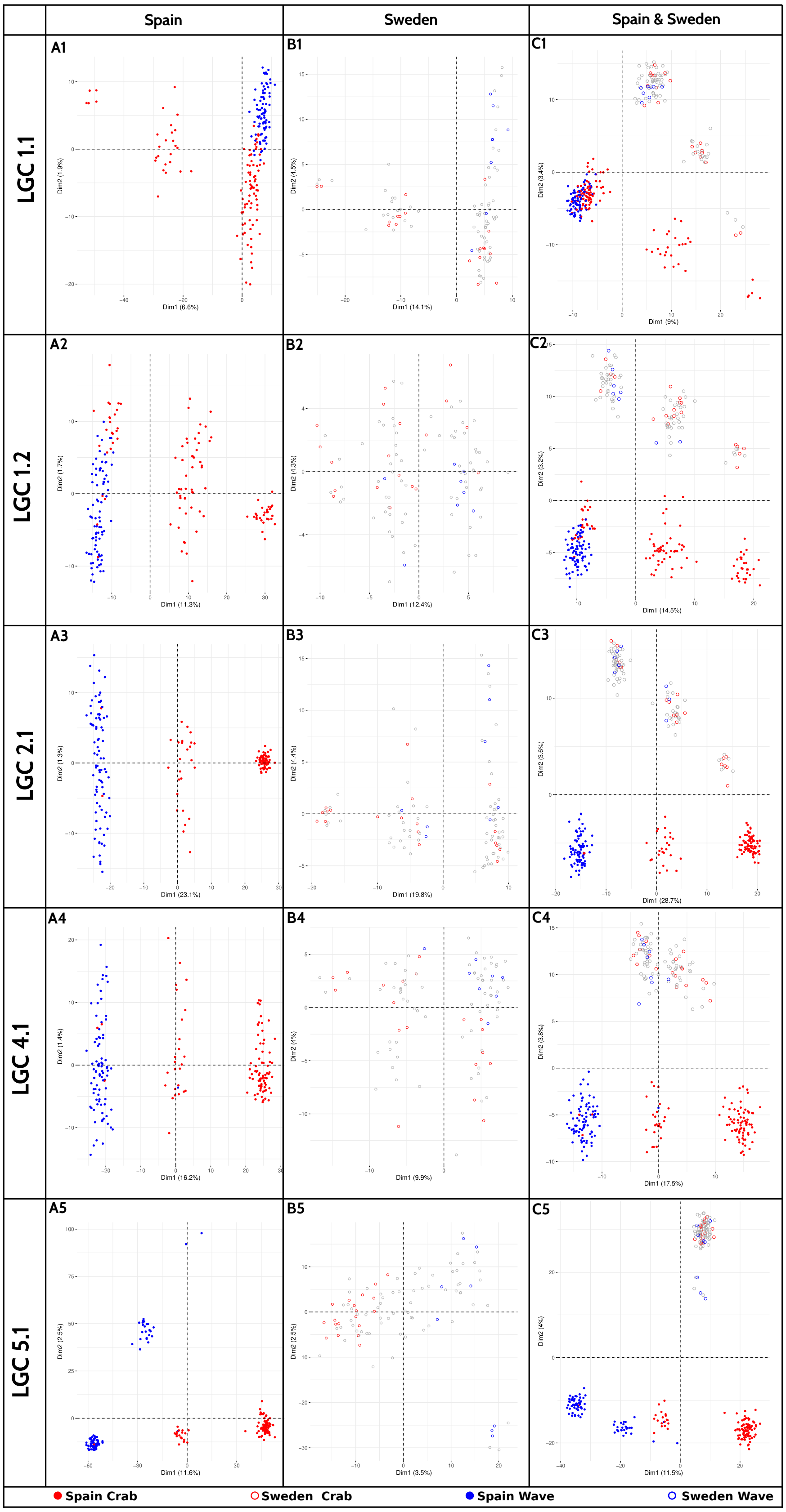

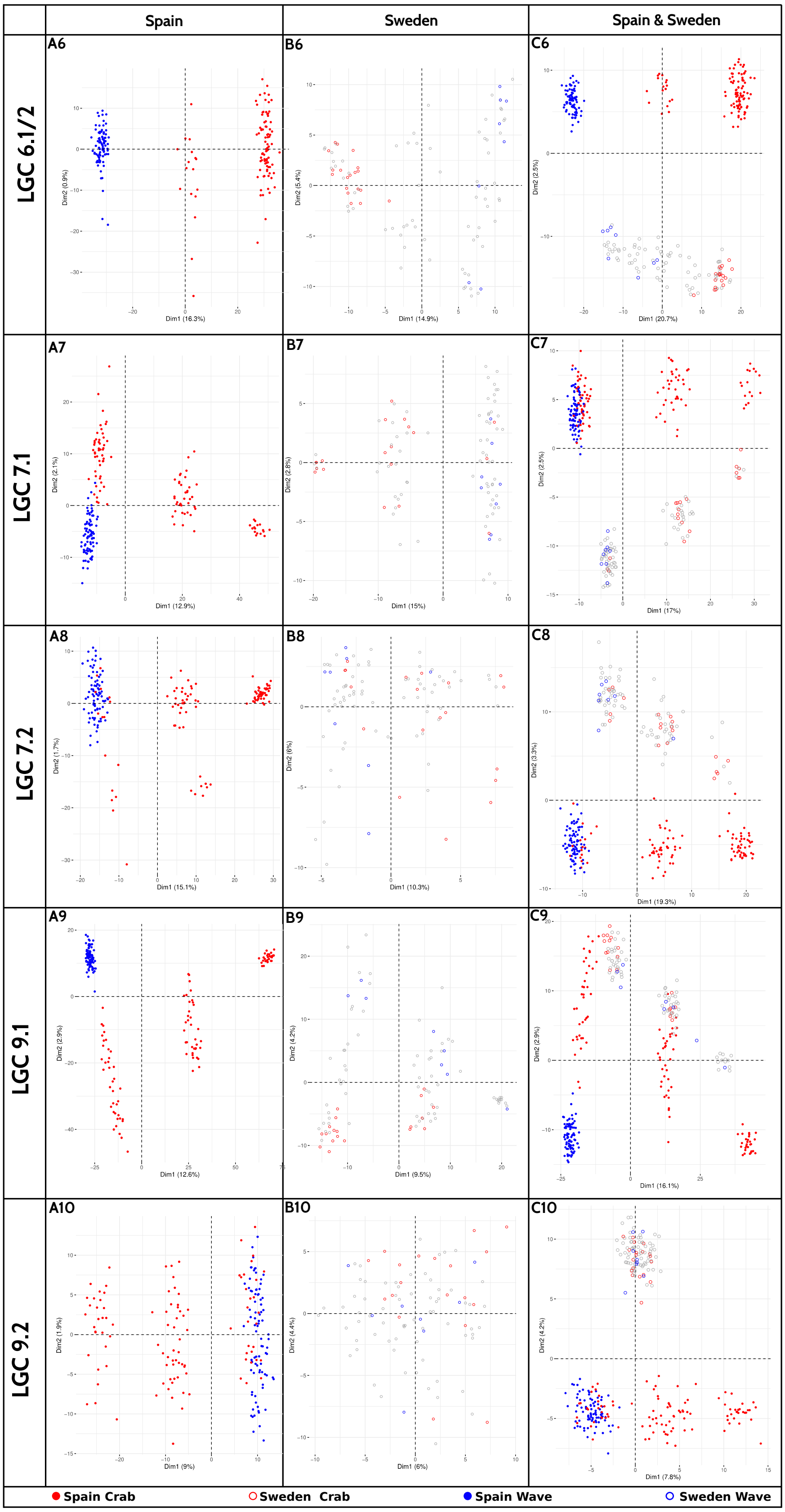
Spain
Sweden
Spain & Sweden
LGC 6.1/2
A6
B6
C6
LGC 7.1
A7
B7
C7
LGC 7.2
A8
B8
C8
LGC 9.1
A9
B9
C9
LGC 9.2
A10
B10
C10
Spain Crab
Sweden Crab
Spain Wave
Sweden Wave

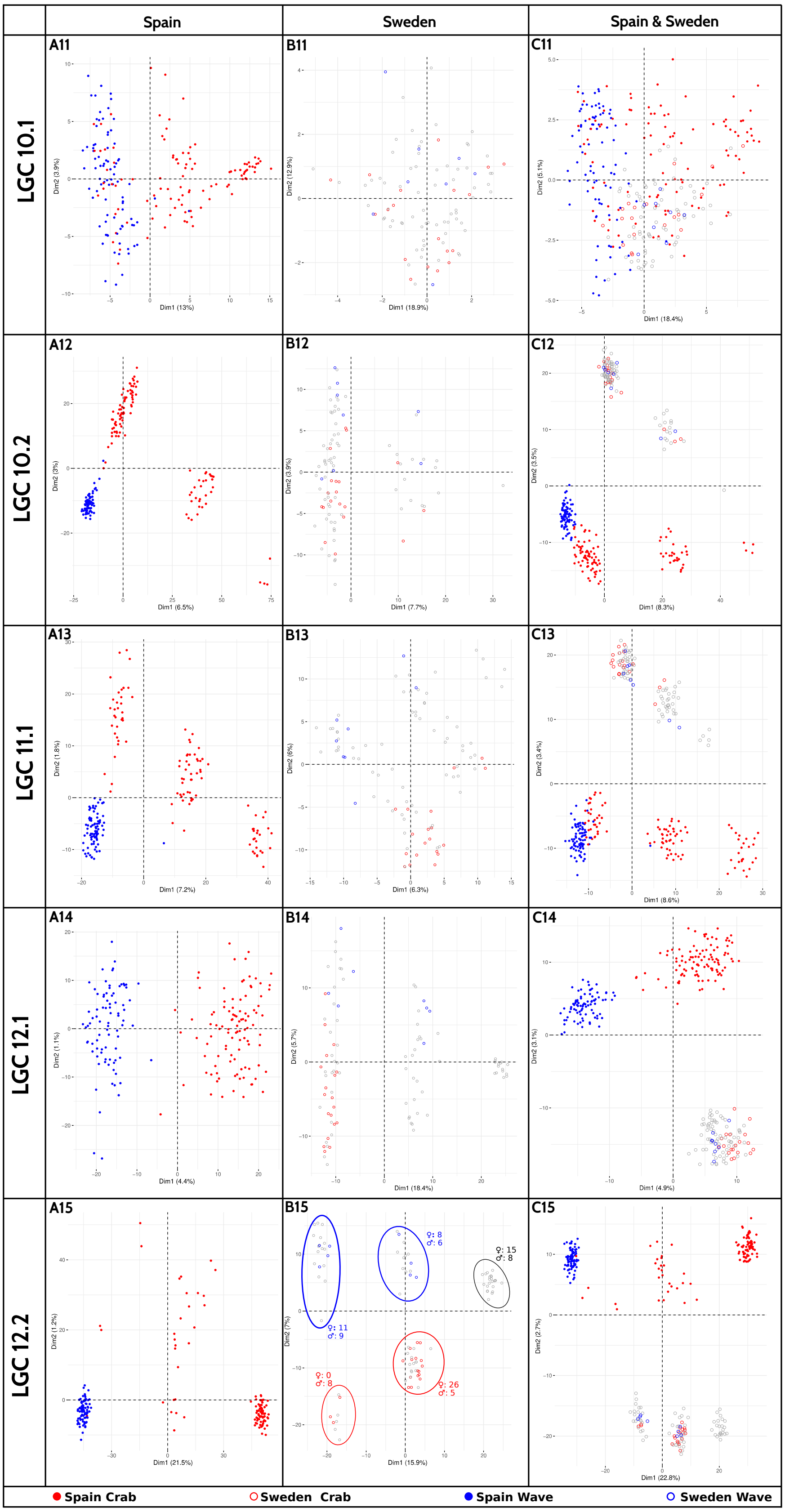
Spain
Sweden
Spain & Sweden
LGC 10.1
LGC 10.2
LGC 11.1
LGC 12.1
LGC 12.2
A11
B11
C11
A12
B12
C12
A13
B13
C13
A14
B14
C14
A15
B15
C15
♀: 11
♂: 9
♀: 8
♂: 6
♀: 15
♂: 8
♀: 0
♂: 8
♀: 26
♂: 5
Spain Crab
Sweden Crab
Spain Wave
Sweden Wave

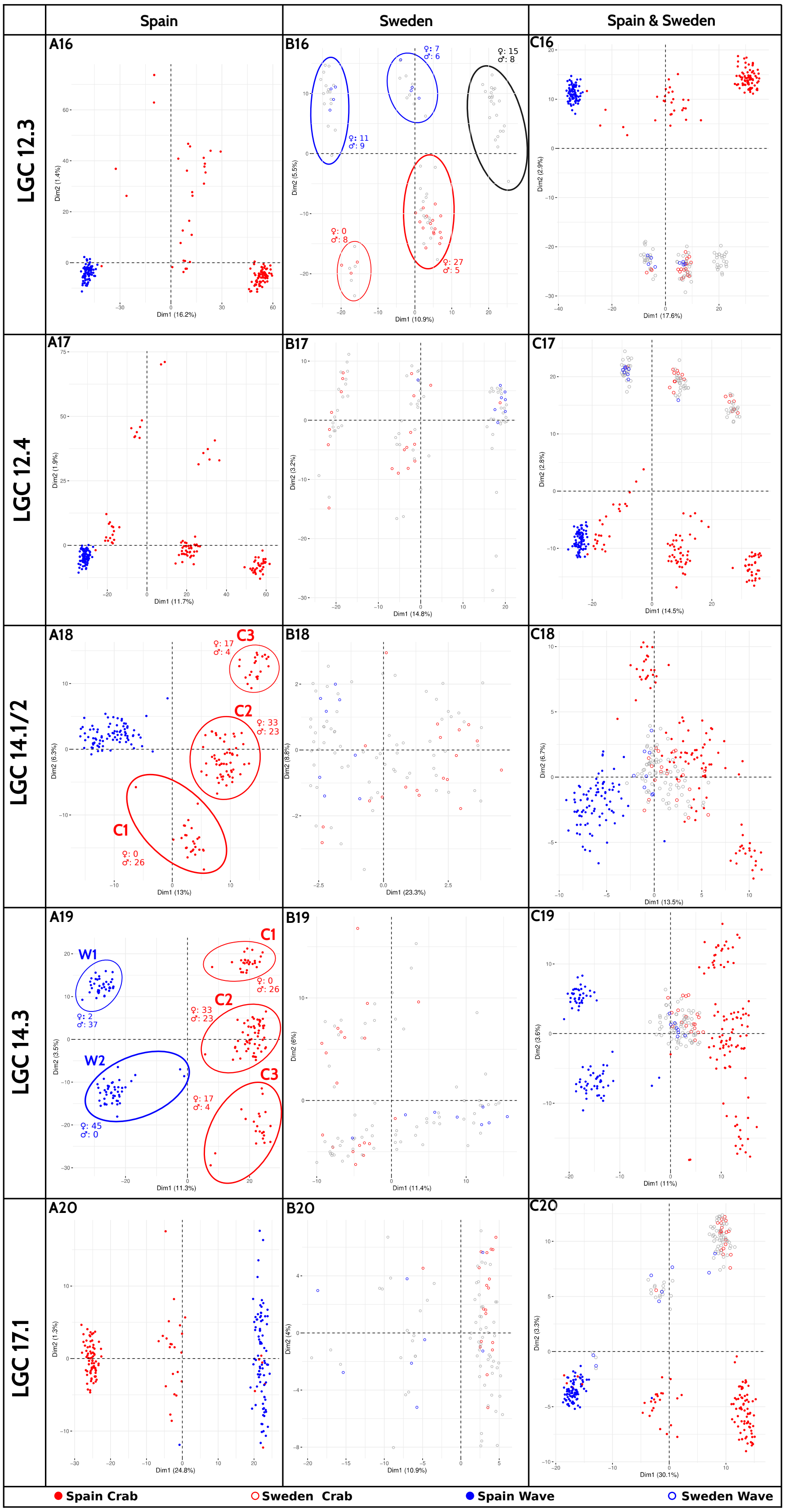
Spain
Sweden
Spain & Sweden
LGC 12.3
LGC 12.4
LGC 14.1/2
LGC 14.3
LGC 17.1
A16
B16
C16
A17
B17
C17
A18
B18
C18
A19
B19
C19
A20
B20
C20
C1
C2
C3
W1
W2
Dim1 (16.2%)
Dim2 (1.4%)
Dim1 (10.9%)
Dim2 (5.5%)
Dim1 (17.6%)
Dim2 (2.9%)
Dim1 (11.7%)
Dim2 (1.9%)
Dim1 (14.8%)
Dim2 (3.2%)
Dim1 (14.5%)
Dim2 (2.8%)
Dim1 (13%)
Dim2 (6.3%)
Dim1 (23.3%)
Dim2 (8.8%)
Dim1 (13.5%)
Dim2 (6.7%)
Dim1 (11.3%)
Dim2 (3.5%)
Dim1 (11.4%)
Dim2 (6%)
Dim1 (11%)
Dim2 (3.6%)
Dim1 (24.8%)
Dim2 (1.3%)
Dim1 (10.9%)
Dim2 (4%)
Dim1 (30.1%)
Dim2 (3.3%)
Spain Crab
Sweden Crab
Spain Wave
Sweden Wave

Figure S14. Distribution of Fst differences between block of SNPs of the same size as an inversion and the rest of the LG in Sweden based on 1000 permutations. Vertical red lines indicates the observed Fst difference between inverted and collinear regions. Vertical grey lines represent quantiles of the distribution.

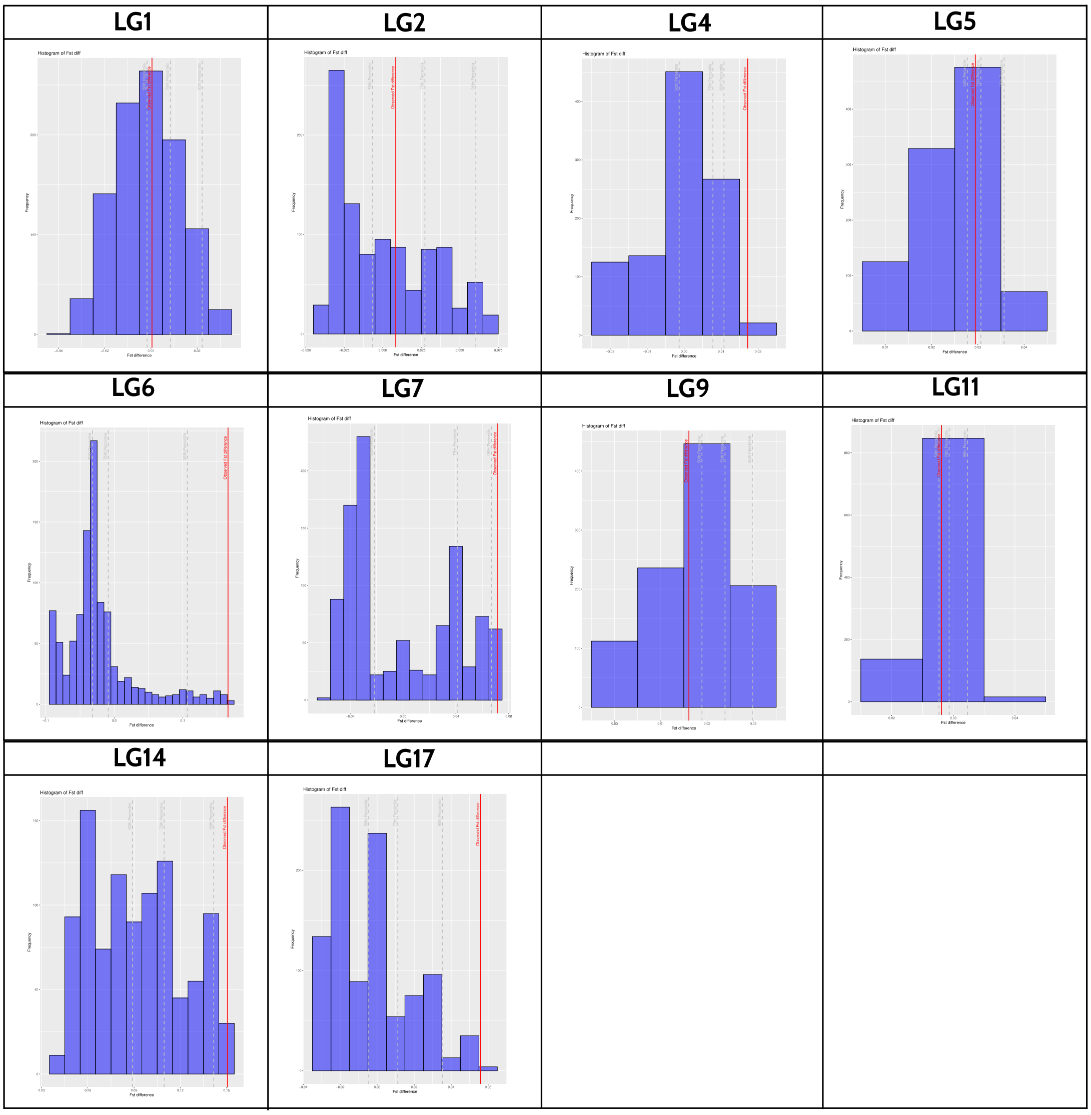

Figure S15. Distribution of Fst differences between block of SNPs of the same size as an inversion and the rest of the LG in Spain based on 1000 permutations. Vertical red lines indicates the observed Fst difference between inverted and collinear regions. Vertical grey lines represent quantiles of the distribution.

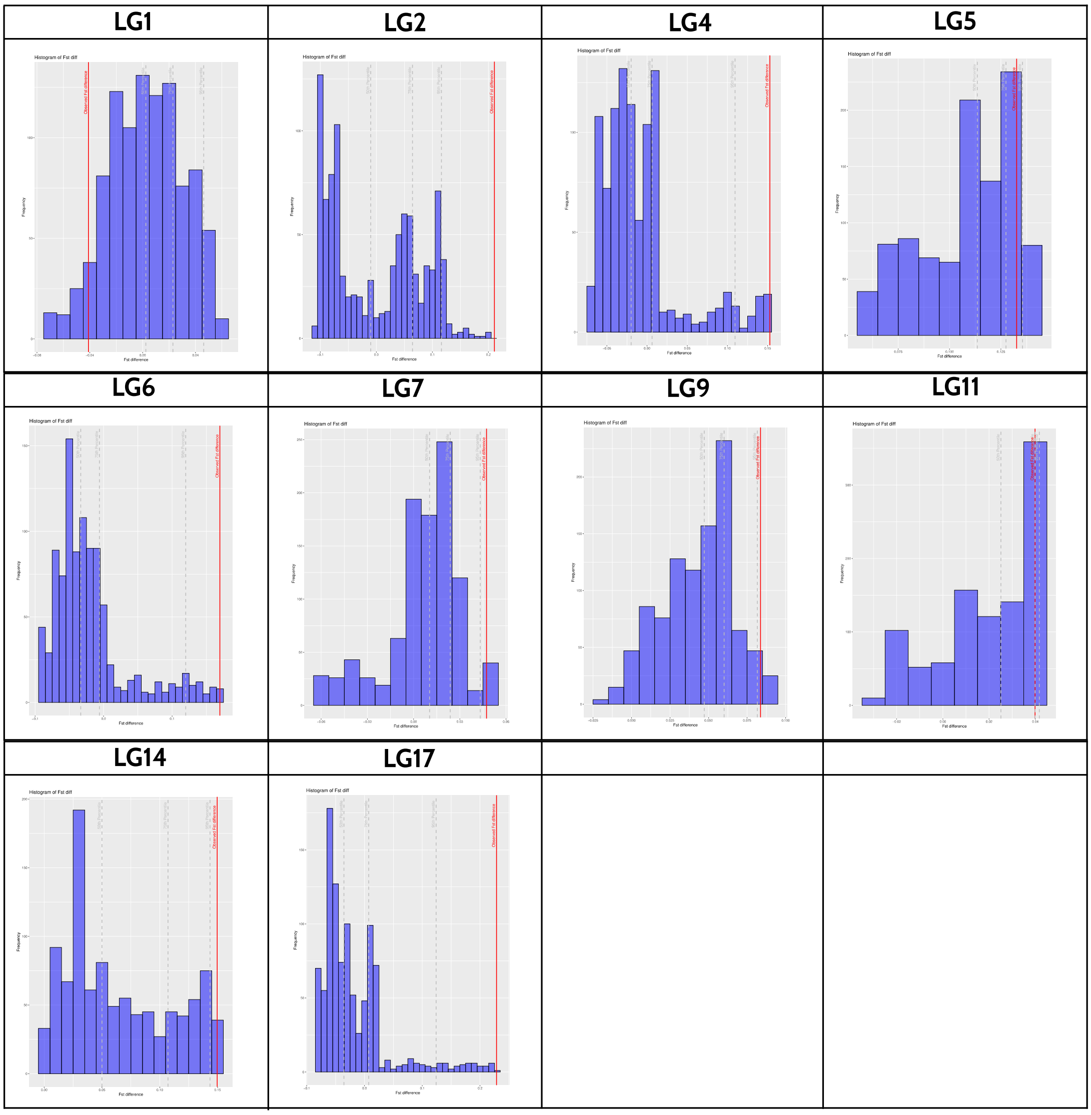

Figure S16. Most abundant arrangement counts against transect position in Spain for each inversion. Red and blue dots represent individual belonging to the Crab and Wave ecotype, respectively.

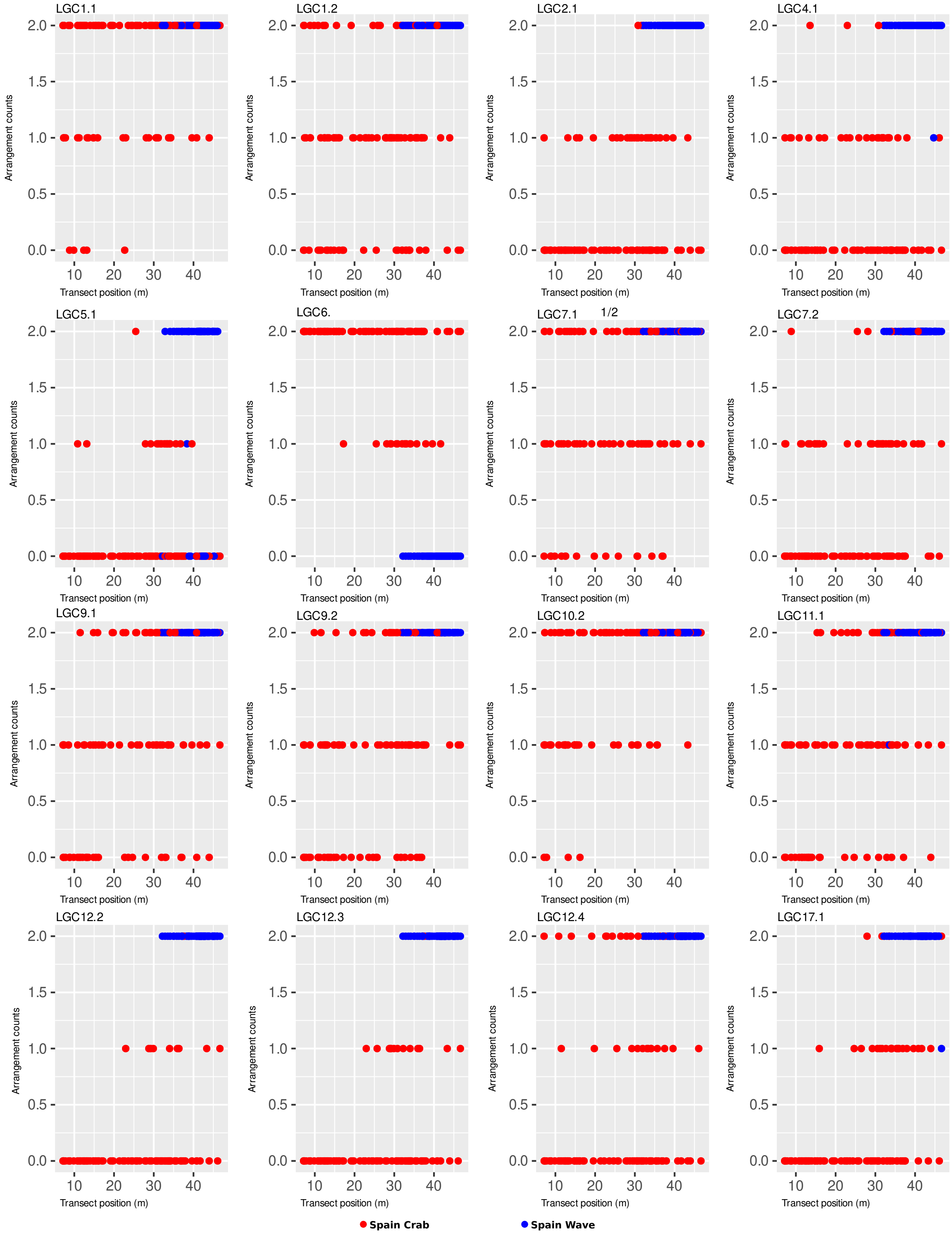

Figure S17. Genomic PC2 scores against transect position in Spain for inversions showing a complete to partial overlap between Crab and Wave genetic group for homozygote individuals carrying the arrangement most abundant in wave. Red and blue dots represent individual belonging to the Crab and Wave ecotype, respectively.

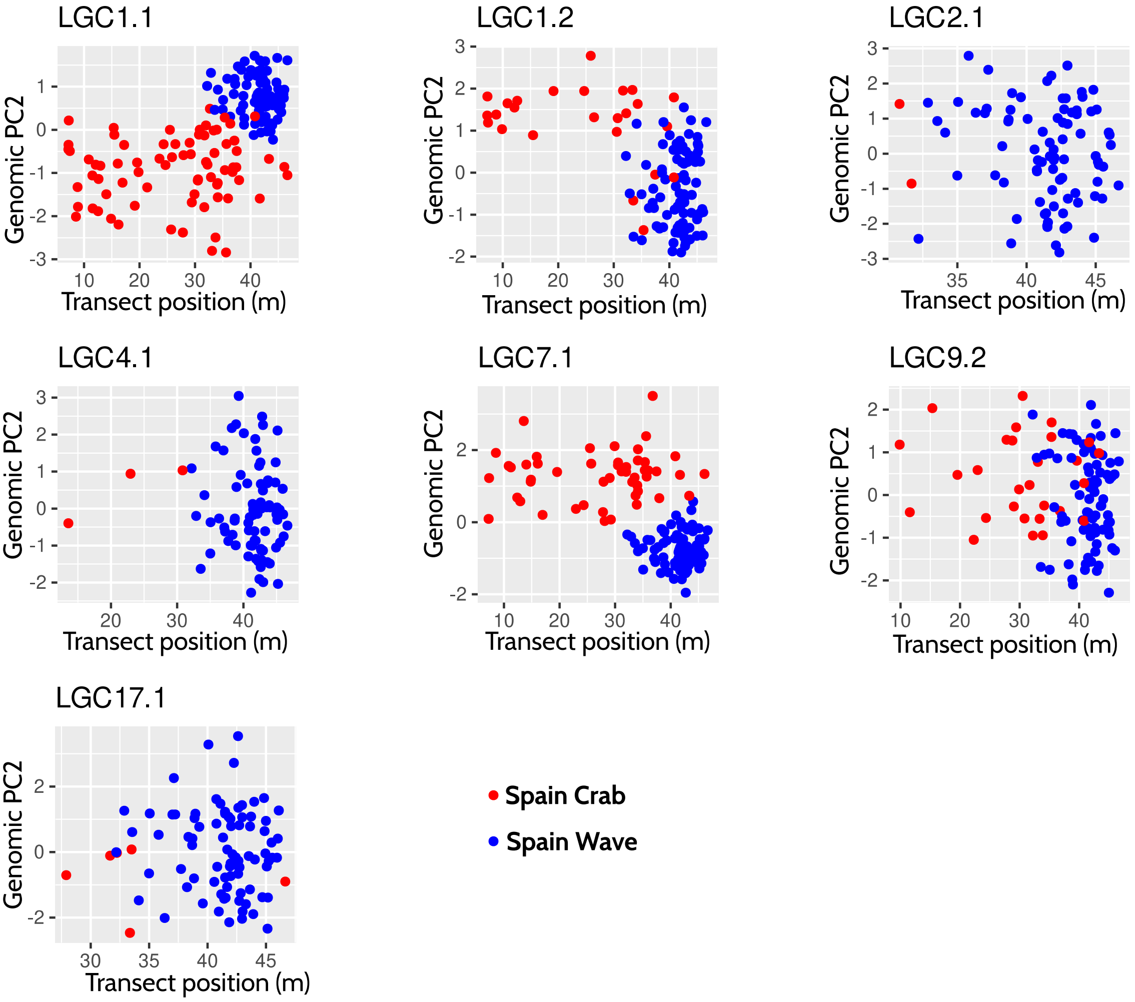

Figure S18. Patterns of genomic divergence in Spain (a, c, e, g, i) and Sweden (b, d, f, h, l) after removal of loci within known inversions as shown by PCA (a, b, c, d), admixture (e, f) and the Hybrid Index (g-l). Transect positions closer to zero correspond to the high shore (Spain) or boulder field (Sweden) while higher values represent the low shore (Spain) or rocky headland (Sweden). Snails in the admixture plots (e, f) were ordered according to their position along shore. In Sweden, a sampling gap occurred at the transect positions 90-120 as seen in the plot of PC1 and Hybrid Index along shore (d, h) and a single genetic cluster was identified but two groups are shown in the admixture plot (e) to facilitate comparisons with Spain. The Crab and Wave group are coloured in blue and red, respectively.

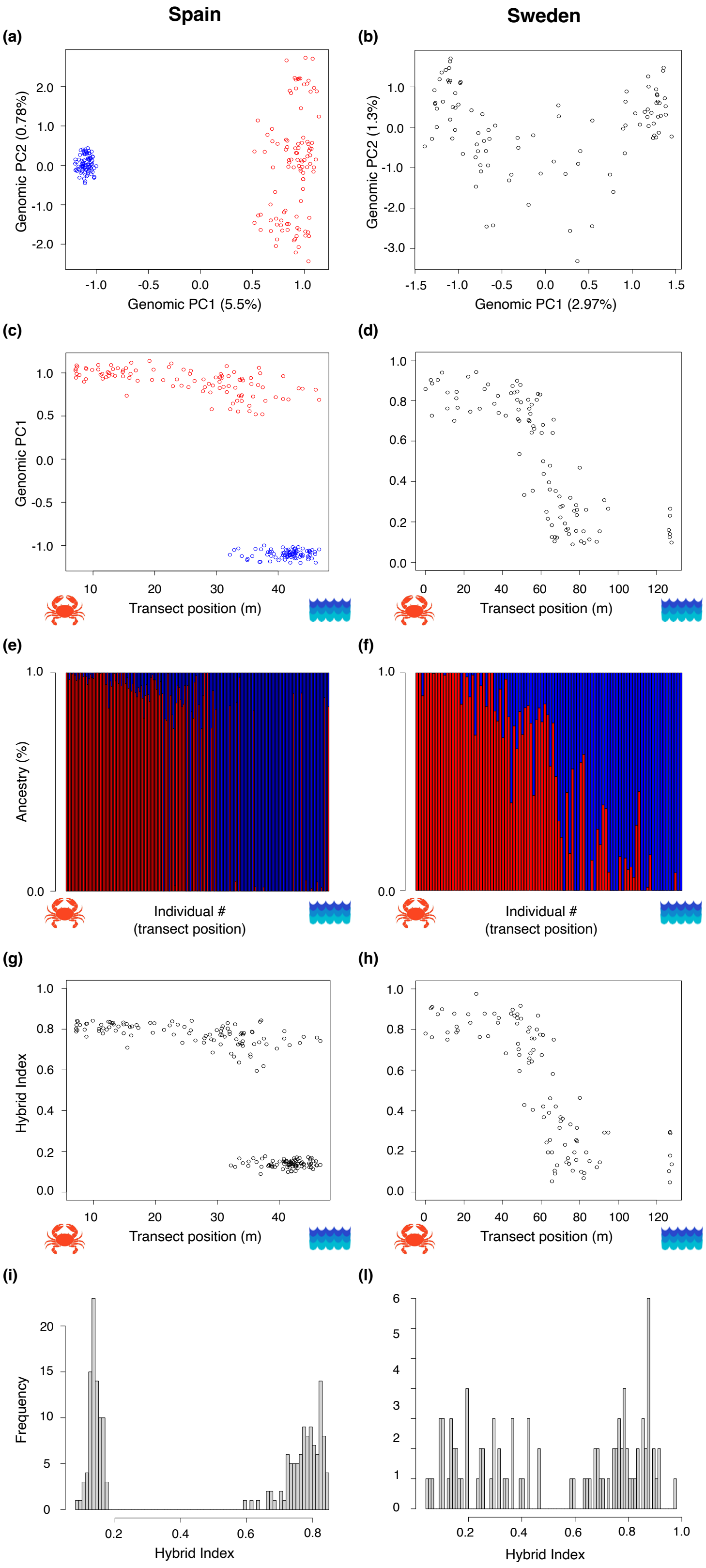

Figure S19. Number of genetic clusters (K) in Spain (a, c) and Sweden (b, d) after removal of loci within known inversions as shown by the discriminant analyses of principal components (DAPCs; a, b) and Admixture cross-validation error (c, d).

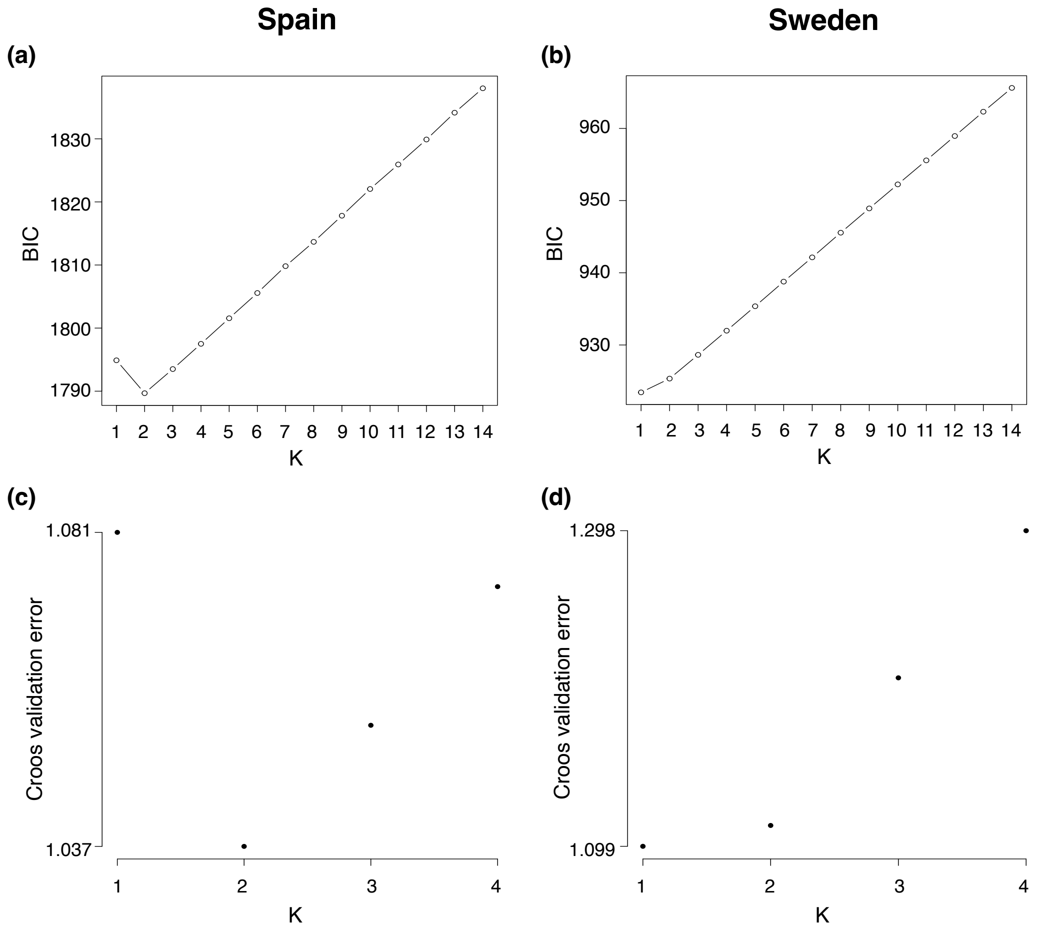

Figure S20. Correlation of the genetic within-cluster PC1 scores in the red genetic group (a) and Blue genetic group (b) among the nine Spanish thinned subsets after removing sites falling within known chromosomal inversions. Abbreviations: s=random SNP subsampling, r= random read subsampling.

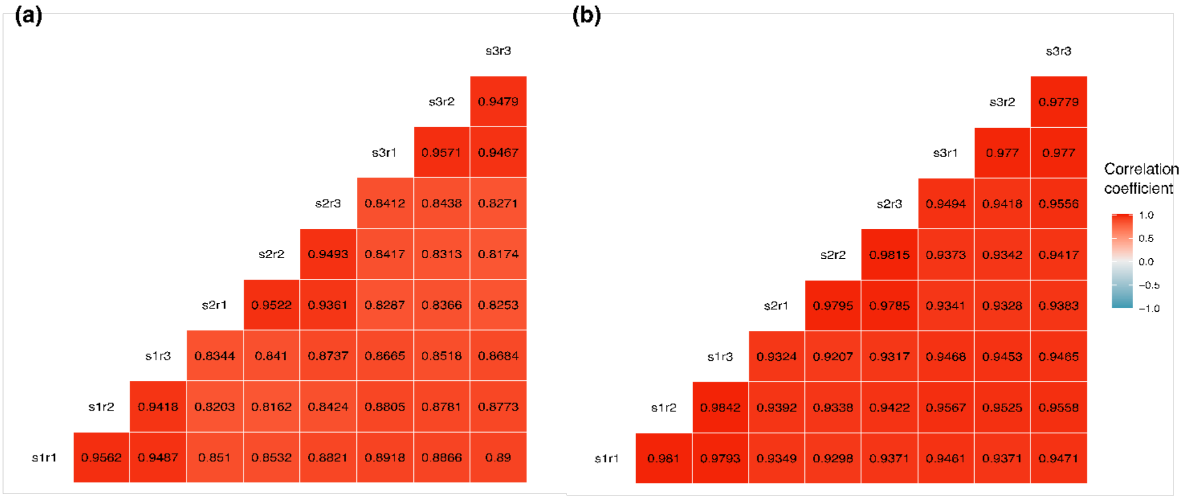

Figure S21. Phenotypic divergence among the Spanish ecotypes as described by the individual traits. Blue and red dots represent individuals belonging to the Wave and Crab ecotype, respectively. With a0_scaled and z_0 corresponding to the two aperture position parameters and convexity = log(g_h) – log(g_w).

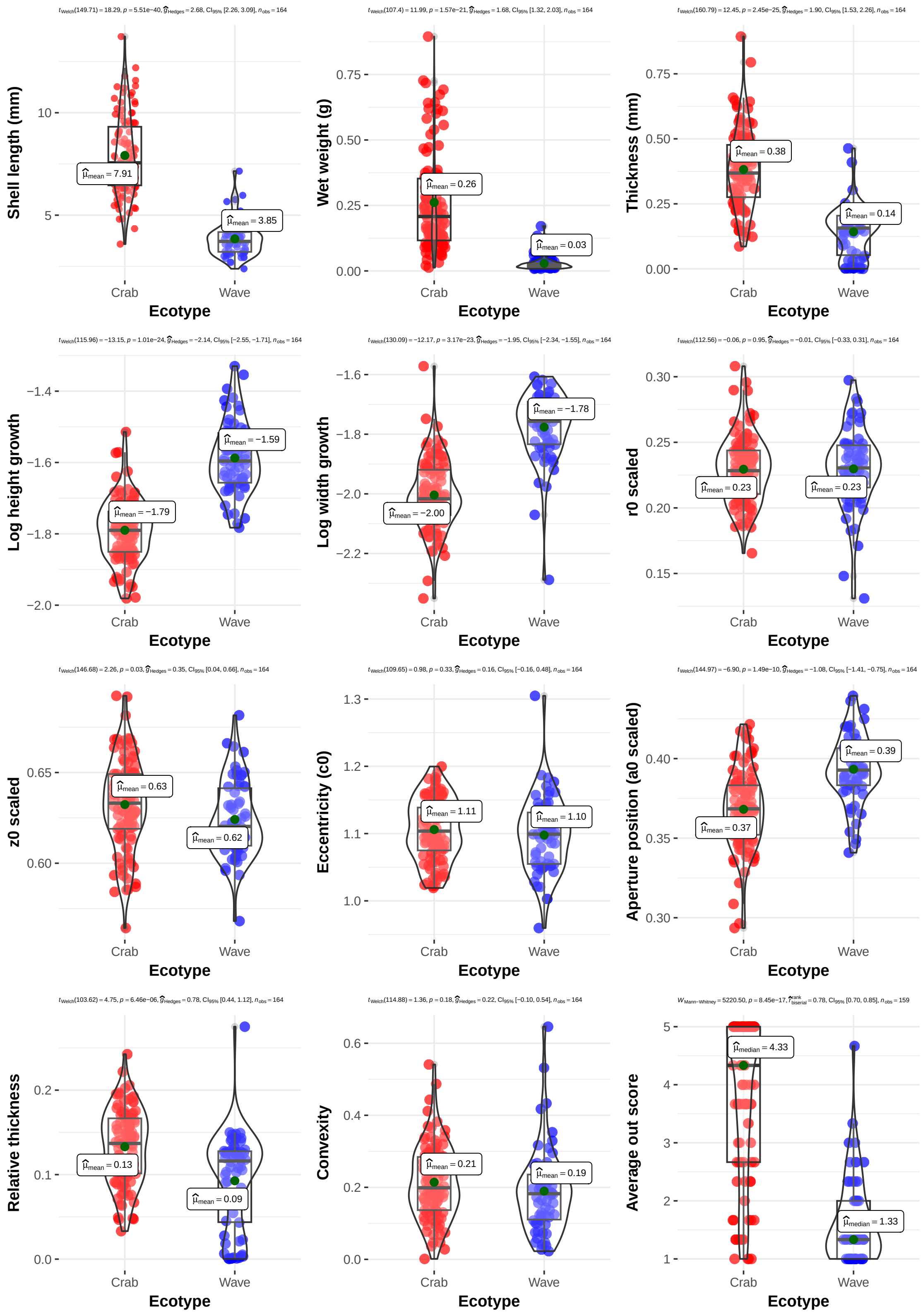

Figure S22. Relative phenotypic differences in individual traits (width growth (gw), height growth (hw), weight, shell length, aperture size (a0_scaled), aperture position (r0_scaled), relative thickness and aperture shape) between ecotypes in Spain (transect from this study)
and Sweden (CZA from Koch *et al.* 2022 *Evolution*). In Spain, these relative values were computed as the difference between the Crab ecotype average and the Wave ecotype average
divided by the Crab ecotype average. In Sweden, these relative values were computed as the difference between the average boulder field end of the transect and the rocky headlands end of the transect average divided by the boulder field end of the transect average.

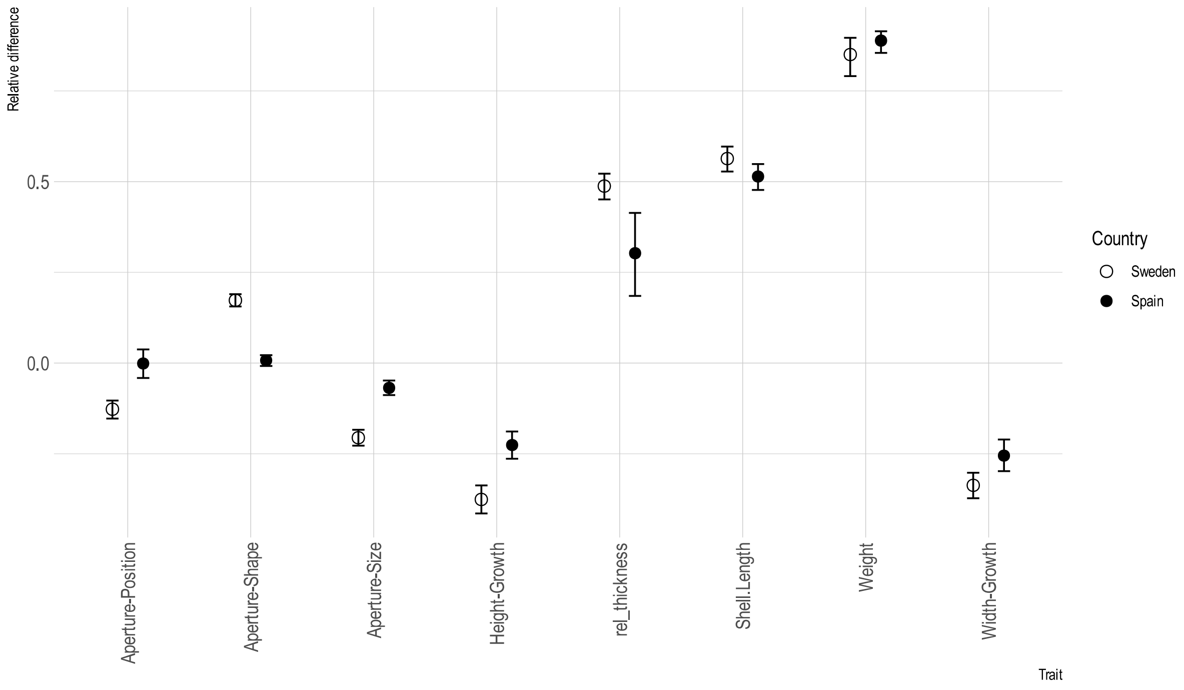

Figure S23. Phenotype PCs along the shore in Spain. PC1 phenotype along transect position (A). PC2 phenotype along transect position (B). In Panel A and B, red and blue colors represent the Crab and Wave ecotype, respectively. PC1 phenotype along transect position in the Crab (C) and Wave (D) ecotype. Grey shaded areas around regressions lines represent the confidence intervals for the regression lines.

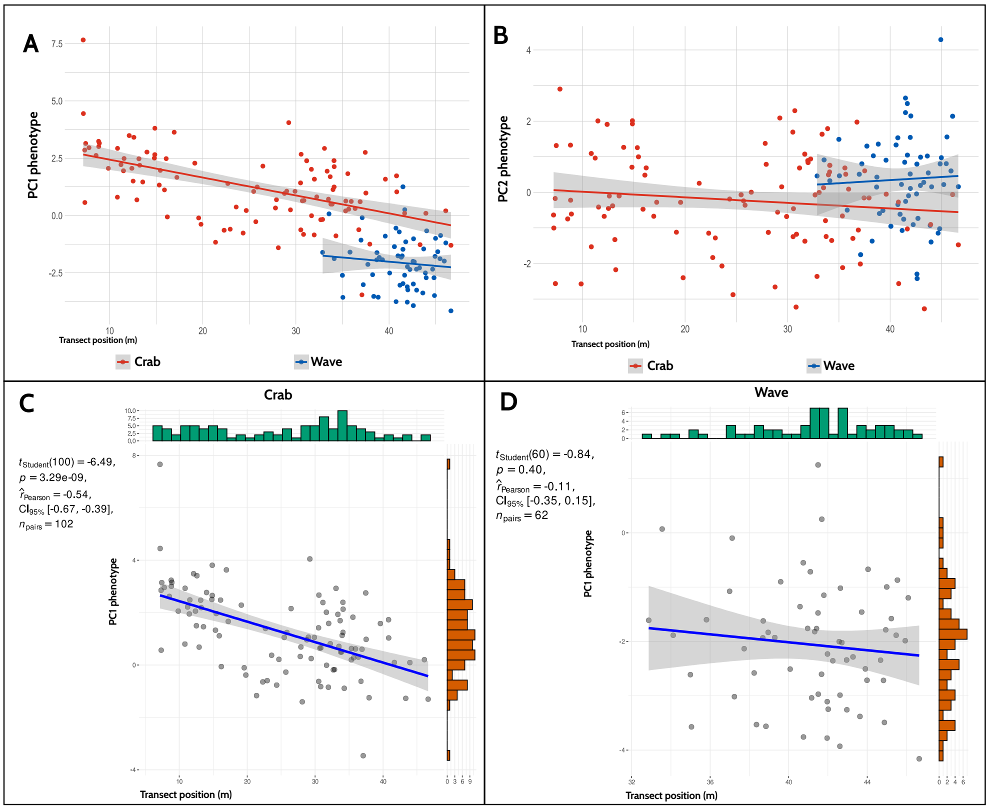

Figure S24. Phenotypic multivariate divergence in the joint dataset including Spain (transect from this study) and Sweden (CZA transect from Koch *et al.* 2022 *Evolution*) as shown by the first two principal components of the phenotypic PCA. The following variables were scaled in each country and included in the PCA: width growth (gw), height growth (gh), weight, shell length, aperture size (a0_scaled), aperture position (r0_scaled), relative thickness and aperture shape. The Spanish Crab and Wave (the two genetic clusters), and as well as the Swedish Crab and Wave (ends of transect) ecotype are represented by blue, red, turquoise, and orange dots, respectively. Grey dots correspond to individuals in or close to the Swedish contact zone (middle portion of the CZA transect). Triangles indicate centroid positions in each ecotype and country. Dotted arrows indicate the shift of the centroids from Wave to Crab ecotype in each country.

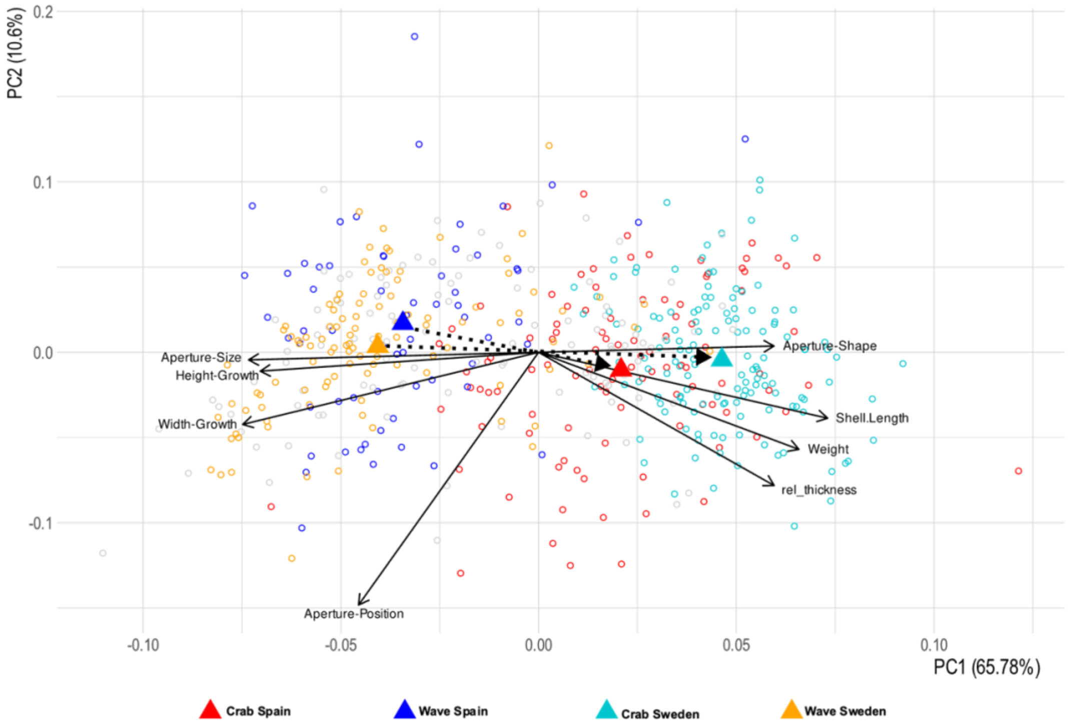

Figure S25. Number of abortive embryos in females following laboratory crosses with single males. A) Females of Spanish Wave ecotype mated with males of Spanish Crab ecotype. B) Females of Spanish Crab ecotype mated with males of same ecotype. C) Females of Spanish Wave ecotype mated with males of same ecotype. In A females and males were from different sites (females; Faro do Roncudo N 43.2747°, W 8.9908°, males; Faro de San Cibrao N 43.6995°, W 7.4349°). In B and C females and males were from the same site (B; Centinela N 42.0772°, W 8.8965°, and C; Faro do Roncudo). Variation among females in abortion rates are high, but in A (between ecotype) the distribution is highly skewed towards very high rates of abortion, while in B and C (within ecotype) the distributions are skewed towards lower abortion rates. All crosses were undertaken in Tjärnö Marine Laboratory.

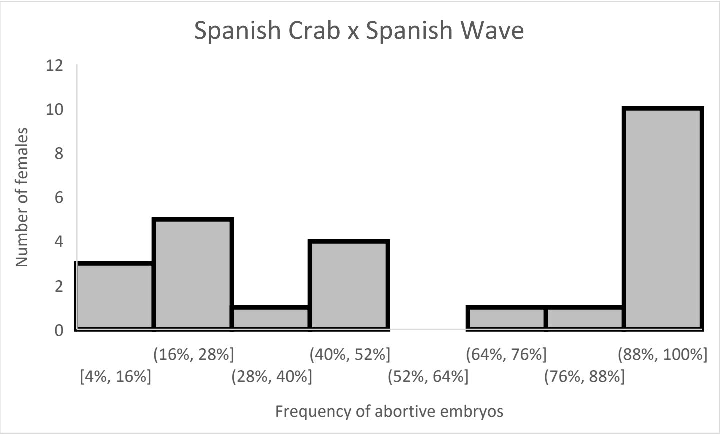


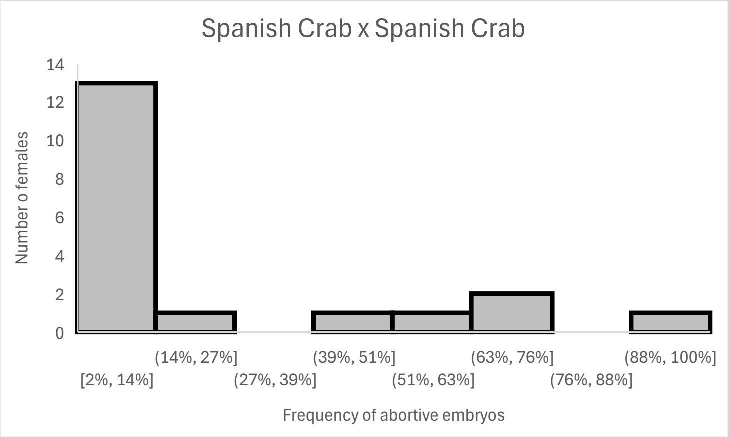

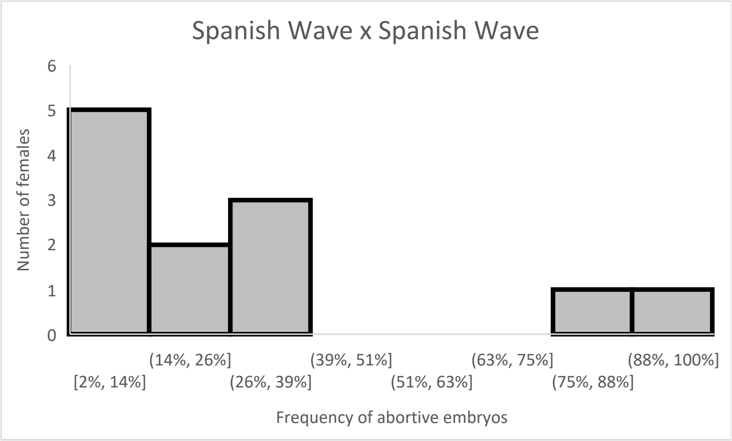
Spanish Wave x Spanish Wave
Number of females
0
1
2
3
4
5
6
[2%, 14%]
(14%, 26%]
(26%, 39%]
(39%, 51%]
(51%, 63%]
(63%, 75%]
(75%, 88%]
(88%, 100%]
Frequency of abortive embryos
