## Supplementary material for "Phenotypic divergence and genomic architecture between parallel ecotypes at two different points on the speciation continuum in a marine snail": Table S6

**Table S6. Details of the linear model results testing for the effect of genetic group and transect position on phenotypical traits.**

**PC1 phenotype**

**Model: PC1 phenotype ~ Genetic group + Position on the shore.**

```
Residuals:
    Min      1Q  Median      3Q     Max
-3.8076 -0.8408  0.0097  0.6652  5.0397

Coefficients:
                 Estimate Std. Error t value Pr(>|t|)
(Intercept)       3.16458    0.30401  10.409  < 2e-16 ***
Genetic_groupRed -2.09161    0.26799  -7.805 7.13e-13 ***
lineD            -0.07601    0.01104  -6.887 1.21e-10 ***
---
Signif. codes:  0 ‘***’ 0.001 ‘**’ 0.01 ‘*’ 0.05 ‘.’ 0.1 ‘ ’ 1

Residual standard error: 1.249 on 161 degrees of freedom
Multiple R-squared:  0.6643,    Adjusted R-squared:  0.6601
F-statistic: 159.3 on 2 and 161 DF,  p-value: < 2.2e-16
```

**PC2 phenotype.**

**Model: PC2 phenotype ~ Genetic group + Position on the shore.**

```
Residuals:
    Min      1Q  Median      3Q     Max
-3.0505 -0.9013  0.0389  0.9062  3.9251

Coefficients:
                 Estimate Std. Error t value Pr(>|t|)
(Intercept)       -0.2212     0.1284  -1.723   0.0868 .
Genetic_groupRed   0.5851     0.2088   2.802   0.0057 **
---
Signif. codes:  0 ‘***’ 0.001 ‘**’ 0.01 ‘*’ 0.05 ‘.’ 0.1 ‘ ’ 1

Residual standard error: 1.297 on 162 degrees of freedom
Multiple R-squared:  0.04622,   Adjusted R-squared:  0.04034
F-statistic: 7.851 on 1 and 162 DF,  p-value: 0.005698
```

**Wet weight.**

**Model: Wet Weight ~ Genetic group * Position on the shore.**

```
Residuals:
     Min       1Q   Median       3Q      Max
-0.35092 -0.04260 -0.00328  0.03793  0.38929

Coefficients:
                         Estimate Std. Error t value Pr(>|t|)
(Intercept)             0.6028788  0.0244998  24.608  < 2e-16 ***
Genetic_groupRed       -0.4765440  0.1613735  -2.953  0.00362 **
lineD                  -0.0135711  0.0008933 -15.193  < 2e-16 ***
Genetic_groupRed:lineD  0.0112112  0.0039607   2.831  0.00524 **
---
Signif. codes:  0 ‘***’ 0.001 ‘**’ 0.01 ‘*’ 0.05 ‘.’ 0.1 ‘ ’ 1

Residual standard error: 0.09847 on 160 degrees of freedom
Multiple R-squared:  0.7359,    Adjusted R-squared:  0.731
F-statistic: 148.6 on 3 and 160 DF,  p-value: < 2.2e-16
```

**Width growth.**

**Model: Width growth ~ Genetic group + Position on the shore.**

```
Residuals:
      Min        1Q    Median        3Q       Max
-0.069068 -0.009958 -0.000427  0.010049  0.067994

Coefficients:
                  Estimate Std. Error t value Pr(>|t|)
(Intercept)      0.1274136  0.0040864  31.180  < 2e-16 ***
Genetic_groupRed 0.0293294  0.0036022   8.142 1.01e-13 ***
lineD            0.0003301  0.0001484   2.225   0.0275 *
---
Signif. codes:  0 ‘***’ 0.001 ‘**’ 0.01 ‘*’ 0.05 ‘.’ 0.1 ‘ ’ 1

Residual standard error: 0.01679 on 161 degrees of freedom
Multiple R-squared:  0.5122,    Adjusted R-squared:  0.5061
F-statistic: 84.51 on 2 and 161 DF,  p-value: < 2.2e-16
```

**Height growth.**

**Model: Height growth ~ Genetic group.**

ONE FACTOR

**r0.**

**Model: r0 ~ Genetic group + Position on the shore.**

```
Residuals:
    Min      1Q  Median      3Q     Max
-0.6947 -0.1994 -0.0321  0.1687  0.8888

Coefficients:
                  Estimate Std. Error t value Pr(>|t|)
(Intercept)       2.419681   0.068705  35.218  < 2e-16 ***
Genetic_groupRed -0.514904   0.060563  -8.502 1.21e-14 ***
lineD            -0.024771   0.002494  -9.931  < 2e-16 ***
---
Signif. codes:  0 ‘***’ 0.001 ‘**’ 0.01 ‘*’ 0.05 ‘.’ 0.1 ‘ ’ 1

Residual standard error: 0.2822 on 161 degrees of freedom
Multiple R-squared:  0.757,     Adjusted R-squared:  0.754
F-statistic: 250.8 on 2 and 161 DF,  p-value: < 2.2e-16
```

**z0.**

**Model: z0 ~ Genetic group + Position on the shore.**

```
Residuals:
     Min       1Q   Median       3Q      Max
-2.19585 -0.44115  0.02447  0.40392  2.89517

Coefficients:
                  Estimate Std. Error t value Pr(>|t|)
(Intercept)       7.222040   0.193841  37.258  < 2e-16 ***
Genetic_groupRed -1.217621   0.170870  -7.126 3.28e-11 ***
lineD            -0.087418   0.007038 -12.422  < 2e-16 ***
---
Signif. codes:  0 ‘***’ 0.001 ‘**’ 0.01 ‘*’ 0.05 ‘.’ 0.1 ‘ ’ 1

Residual standard error: 0.7963 on 161 degrees of freedom
Multiple R-squared:  0.7803,    Adjusted R-squared:  0.7776
F-statistic: 285.9 on 2 and 161 DF,  p-value: < 2.2e-16
```

**a0.**

**Model: a0 ~ Genetic group + Position on the shore.**

```
Residuals:
     Min       1Q   Median       3Q      Max
-1.27715 -0.24126  0.02492  0.25046  1.09431

Coefficients:
                  Estimate Std. Error t value Pr(>|t|)
(Intercept)       3.973277   0.097584  40.716  < 2e-16 ***
Genetic_groupRed -0.694802   0.086020  -8.077 1.47e-13 ***
lineD            -0.042946   0.003543 -12.122  < 2e-16 ***
---
Signif. codes:  0 ‘***’ 0.001 ‘**’ 0.01 ‘*’ 0.05 ‘.’ 0.1 ‘ ’ 1

Residual standard error: 0.4009 on 161 degrees of freedom
Multiple R-squared:  0.7902,    Adjusted R-squared:  0.7876
F-statistic: 303.3 on 2 and 161 DF,  p-value: < 2.2e-16
```

**Eccentricity.**

**Model: Eccentricity ~ Genetic group + Position on the shore.**

```
Residuals:
    Min      1Q  Median      3Q     Max
-1.5828 -0.2659 -0.0125  0.3065  1.2837

Coefficients:
                  Estimate Std. Error t value Pr(>|t|)
(Intercept)       4.438562   0.112631  39.408  < 2e-16 ***
Genetic_groupRed -0.759235   0.099284  -7.647 1.76e-12 ***
lineD            -0.049085   0.004089 -12.004  < 2e-16 ***
---
Signif. codes:  0 ‘***’ 0.001 ‘**’ 0.01 ‘*’ 0.05 ‘.’ 0.1 ‘ ’ 1

Residual standard error: 0.4627 on 161 degrees of freedom
Multiple R-squared:  0.7813,    Adjusted R-squared:  0.7786
F-statistic: 287.6 on 2 and 161 DF,  p-value: < 2.2e-16
```

**apAngle.**

**Model: apAngle ~ Genetic group .**

ONE FACTOR

**Shell length.**

**Model: Shell length ~ Genetic group + Position on the shore.**

```
Residuals:
    Min      1Q  Median      3Q     Max
-3.5808 -0.6375 -0.0178  0.6716  3.4528

Coefficients:
                 Estimate Std. Error t value Pr(>|t|)
(Intercept)       11.2039     0.2809  39.890  < 2e-16 ***
Genetic_groupRed  -1.9701     0.2476  -7.957 2.96e-13 ***
lineD             -0.1308     0.0102 -12.823  < 2e-16 ***
---
Signif. codes:  0 ‘***’ 0.001 ‘**’ 0.01 ‘*’ 0.05 ‘.’ 0.1 ‘ ’ 1

Residual standard error: 1.154 on 161 degrees of freedom
Multiple R-squared:    0.8,     Adjusted R-squared:  0.7975
F-statistic: 321.9 on 2 and 161 DF,  p-value: < 2.2e-16
```

**Thickness.**

**Model: Thickness ~ Genetic group + Position on the shore.**

```
Residuals:
     Min       1Q   Median       3Q      Max
-0.30410 -0.09602  0.01078  0.06629  0.44691

Coefficients:
                  Estimate Std. Error t value Pr(>|t|)
(Intercept)       0.472026   0.031211  15.124  < 2e-16 ***
Genetic_groupRed -0.182458   0.027512  -6.632 4.78e-10 ***
lineD            -0.003572   0.001133  -3.152  0.00193 **
---
Signif. codes:  0 ‘***’ 0.001 ‘**’ 0.01 ‘*’ 0.05 ‘.’ 0.1 ‘ ’ 1

Residual standard error: 0.1282 on 161 degrees of freedom
Multiple R-squared:  0.4735,    Adjusted R-squared:  0.467
F-statistic:  72.4 on 2 and 161 DF,  p-value: < 2.2e-16
```

**a0 scaled.**

**Model: a0 scaled ~ Genetic group + Position on the shore.**

```
Residuals:
      Min        1Q    Median        3Q       Max
-0.074350 -0.013738  0.000323  0.014891  0.045201

Coefficients:
                  Estimate Std. Error t value Pr(>|t|)
(Intercept)      0.3519138  0.0055872  62.985  < 2e-16 ***
Genetic_groupRed 0.0148380  0.0049251   3.013  0.00301 **
lineD            0.0006453  0.0002028   3.181  0.00176 **
---
Signif. codes:  0 ‘***’ 0.001 ‘**’ 0.01 ‘*’ 0.05 ‘.’ 0.1 ‘ ’ 1

Residual standard error: 0.02295 on 161 degrees of freedom
Multiple R-squared:   0.26,     Adjusted R-squared:  0.2508
F-statistic: 28.29 on 2 and 161 DF,  p-value: 2.966e-11
```

**z0 scaled.**

**Model: z0 scaled ~ Position on the shore.**

```
Residuals:
      Min        1Q    Median        3Q       Max
-0.057570 -0.015677 -0.001155  0.015541  0.062279

Coefficients:
                   Estimate Std. Error t value Pr(>|t|)
(Intercept)       0.6468767  0.0056053 115.404  < 2e-16 ***
Genetic_groupRed  0.0010694  0.0049411   0.216  0.82893
lineD            -0.0005778  0.0002035  -2.839  0.00511 **
---
Signif. codes:  0 ‘***’ 0.001 ‘**’ 0.01 ‘*’ 0.05 ‘.’ 0.1 ‘ ’ 1

Residual standard error: 0.02303 on 161 degrees of freedom
Multiple R-squared:  0.07446,   Adjusted R-squared:  0.06296
F-statistic: 6.476 on 2 and 161 DF,  p-value: 0.001971
```

**Log height growth.**

**Model: Log height growth ~ Genetic group.**

```
Residuals:
      Min        1Q    Median        3Q       Max
-0.198767 -0.062138 -0.002536  0.055177  0.279545

Coefficients:
                   Estimate Std. Error t value Pr(>|t|)
(Intercept)      -1.8070473  0.0225810 -80.025   <2e-16 ***
Genetic_groupRed  0.1916986  0.0199051   9.631   <2e-16 ***
lineD             0.0006805  0.0008198   0.830    0.408
---
Signif. codes:  0 ‘***’ 0.001 ‘**’ 0.01 ‘*’ 0.05 ‘.’ 0.1 ‘ ’ 1

Residual standard error: 0.09276 on 161 degrees of freedom
Multiple R-squared:  0.5342,    Adjusted R-squared:  0.5285
F-statistic: 92.33 on 2 and 161 DF,  p-value: < 2.2e-16
```

**Log width growth.**

**Model: Log width growth ~ Genetic group + Position on the shore.**

```
Residuals:
     Min       1Q   Median       3Q      Max
-0.51334 -0.06512  0.00441  0.07588  0.40367

Coefficients:
                  Estimate Std. Error t value Pr(>|t|)
(Intercept)      -2.064058   0.027995 -73.730  < 2e-16 ***
Genetic_groupRed  0.189610   0.024677   7.684 1.43e-12 ***
lineD             0.002386   0.001016   2.348   0.0201 *
---
Signif. codes:  0 ‘***’ 0.001 ‘**’ 0.01 ‘*’ 0.05 ‘.’ 0.1 ‘ ’ 1

Residual standard error: 0.115 on 161 degrees of freedom
Multiple R-squared:  0.4936,    Adjusted R-squared:  0.4873
F-statistic: 78.47 on 2 and 161 DF,  p-value: < 2.2e-16
```

**Relative thickness.**

**Model: Relative thickness ~ Genetic group.**

```
Residuals:
     Min       1Q   Median       3Q      Max
-0.09586 -0.03776  0.01076  0.03342  0.18119

Coefficients:
                   Estimate Std. Error t value Pr(>|t|)
(Intercept)       0.1122762  0.0118844   9.447  < 2e-16 ***
Genetic_groupRed -0.0534488  0.0104761  -5.102 9.36e-07 ***
lineD             0.0008229  0.0004315   1.907   0.0583 .
---
Signif. codes:  0 ‘***’ 0.001 ‘**’ 0.01 ‘*’ 0.05 ‘.’ 0.1 ‘ ’ 1

Residual standard error: 0.04882 on 161 degrees of freedom
Multiple R-squared:  0.1564,    Adjusted R-squared:  0.1459
F-statistic: 14.92 on 2 and 161 DF,  p-value: 1.136e-06
```
